# Monitoring population extinction risk with community science data

**DOI:** 10.1101/2025.06.19.660569

**Authors:** Orlando Acevedo-Charry, José M. Ponciano, Caroline L. Poli, Brian M. Jeffery, Robert J. Fletcher, María Ángela Echeverry-Galvis, Bette A. Loiselle, Scott K. Robinson, Miguel A. Acevedo

**Affiliations:** School of Natural Resources and Environment, IFAS, University of Florida, Gainesville, FL 32611, USA; Department of Wildlife Ecology and Conservation, IFAS, University of Florida, Gainesville, FL 32611, USA; Florida Museum of Natural History, University of Florida, Gainesville, FL 32611, USA; Center for Latin American Studies and Tropical Conservation Development Program, University of Florida, Gainesville, FL 32611, USA; Department of Biology, University of Florida, Gainesville, FL 32611, USA; Fish and Wildlife Research Institute, Florida Fish and Wildlife Conservation Commission, 1105 SW Williston Rd, Gainesville, FL 32601, USA; Department of Zoology, Conservation Research Institute, University of Cambridge, Cambridge, UK; Departamento de Ecología y Territorio, Pontificia Universidad Javeriana, Bogotá D.C., Colombia

**Keywords:** citizen science, diffusion process, eBird, exponential stochastic population model, Gompertz stochastic population model, risk-based population viability analysis, state-space population models

## Abstract

The robust estimation of local extinction risk is central to inform management and conservation efforts. Still, estimating this key demographic parameter requires standardized monitoring data that are lacking for most species and systems. The analysis of community science data is emerging as a promising alternative. These expansive datasets leverage observations from multiple volunteers that provide higher temporal and spatial resolution. Nevertheless, the proper analysis of community science data is challenging because it requires accounting for additional complexities in the intrinsic ecological and observational processes.

To address this issue, we describe and test a quantitative approach that fits continuous state-space models iteratively to eBird data with the ultimate goal of estimating local persistence probability through time.

We evaluated model accuracy by comparing estimates and trends from eBird with those from the endangered Everglades’ snail kite long-term, standardized monitoring project. We also performed two separate sensitivity analyses (temporal and sampling thinning) to assess how robust the persistence estimates are to a reduction in the number of eBird observations available.

Our results showed that the temporal trend trajectory of local population persistence estimated from eBird closely matched that from standardized monitoring. Moreover, the trend remained similar even when reducing the amount of eBird data available to 5% of the original data set – a reduction from 258 to 13 weeks or from 7,714 to 385 lists of observations across 5 years of monitoring.

**Synthesis and applications:** Our modeling framework provides a robust, computationally efficient, and easy-to-apply tool for monitoring local persistence probability that can support global conservation efforts. This will complement the monitoring of species population viability in places where standardized monitoring is still lacking, but community science observations are common.

## 1 | INTRODUCTION

Monitoring population trends and assessing extinction risk is a priority in applied ecology and conservation (Bakker & Doak, 2009; Callaghan et al., 2024; Dennis et al., 1991; Fink, Johnston, et al., 2023; Neate-Clegg et al., 2020; Staples et al., 2005). However, accurate trend predictions are often constrained by the limited availability of standardized biodiversity monitoring data that are costly and logistically challenging to gather, particularly for species in understudied regions (Lees et al., 2022; Robinson et al., 2021; Simmons et al., 2019). Community science data, which is gathered by volunteers and often referred to as citizen or participatory science, has emerged as a promising alternative to address these data gaps (Amano et al., 2016; Crawford et al., 2020; Dennis et al., 2017; Johnston et al., 2025; Kelling et al., 2019). Yet, these opportunistic datasets present multiple complexities, requiring advanced modeling approaches to ensure they accurately reflect ecological processes (Boyd et al., 2023; Dennis et al., 2006; Johnston et al., 2023, 2025).

Community science data has the potential to inform population viability – the minimum conditions required for the long-term persistence of a population (Bakker & Doak, 2009; Crawford et al., 2020; Kéry et al., 2010; Staples et al., 2005; Trouillier et al., 2023). However, challenges remain to use community science data for biodiversity monitoring appropriately (Johnston et al., 2023). The opportunistic nature of most community science data initiatives has inherent spatial, observer, and reporting preference biases (Johnston et al., 2021), leading to heterogeneous data that are challenging to analyze (Johnston et al., 2023). Therefore, since more data does not necessarily mean correct representation of true ecological processes (Boyd et al., 2023; Meng, 2018), modeling approaches accounting for multiple data complexities are needed to analyze community science data (Dennis et al., 2006; Fink, Johnston, et al., 2023; Johnston et al., 2023, 2025). Although recent analytical advances have addressed some of these challenges (Fink, Johnston, et al., 2023; Johnston et al., 2021, 2023; Stillman et al., 2023; Zhao et al., 2024), crucial limitations persist. For example, most of these approaches remain resource-intensive and infeasible to apply in many regions with reduced community science data, scarcity of standardized monitoring data, or limited computer power; thus, limited accessibility to powerful but complex tools continues to impede their broader use (Johnston et al., 2023). In addition, the application of extinction theory to assess population risk from community science data is still conspicuously uncommon (Bakker & Doak, 2009; Ferguson & Ponciano, 2014; Staples et al., 2005), despite its potential to provide key insights about population trends (Crawford et al., 2020; Dennis et al., 2017; Fink, Johnston, et al., 2023; Johnston et al., 2025; Kéry et al., 2010). Therefore, to appropriately leverage community science data, there is still a need for modeling approaches that provide a balance between robustness during the modeling, disentangle observation and ecological processes, ease of application, and computational efficiency.

Here we propose and test a risk-based Viable Population Monitoring framework (VPM; Figure 1; Staples et al., 2005) as a robust and computationally efficient framework to estimate local population persistence from community science data. VPM is a promising tool to estimate the risk of quasi-extinction (Staples et al., 2005), which is defined as the species probability to decrease below pre-established abundance thresholds and become effectively extirpated (Ginzburg et al., 1982; Trouillier et al., 2023). In addition, VPM can integrate state-space models that help address the complexities from community science data, including disentangling observation and environmental noise. State-space population dynamic models link observed time series data to latent population processes (Dennis & Ponciano, 2014; Humbert et al., 2009; Newman et al., 2014, 2023). These state-space models provide robust estimates of population growth rates, density dependence (when present), and independent variability in ecological and observational processes (Dennis et al., 2006; Dennis & Otten, 2000; Evans et al., 2023; Hostetler & Chandler, 2015; Ponciano et al., 2009). The VPM framework uses those estimated population parameters from the state-space models to simulate near-future trajectories and estimate population quasi-extinction probabilities and its mathematical complement, the local persistence probability (φ). Furthermore, the VPM iteratively updates these estimates with new data collection, monitoring the population extinction risk (Figure 1).

**Figure 1.**
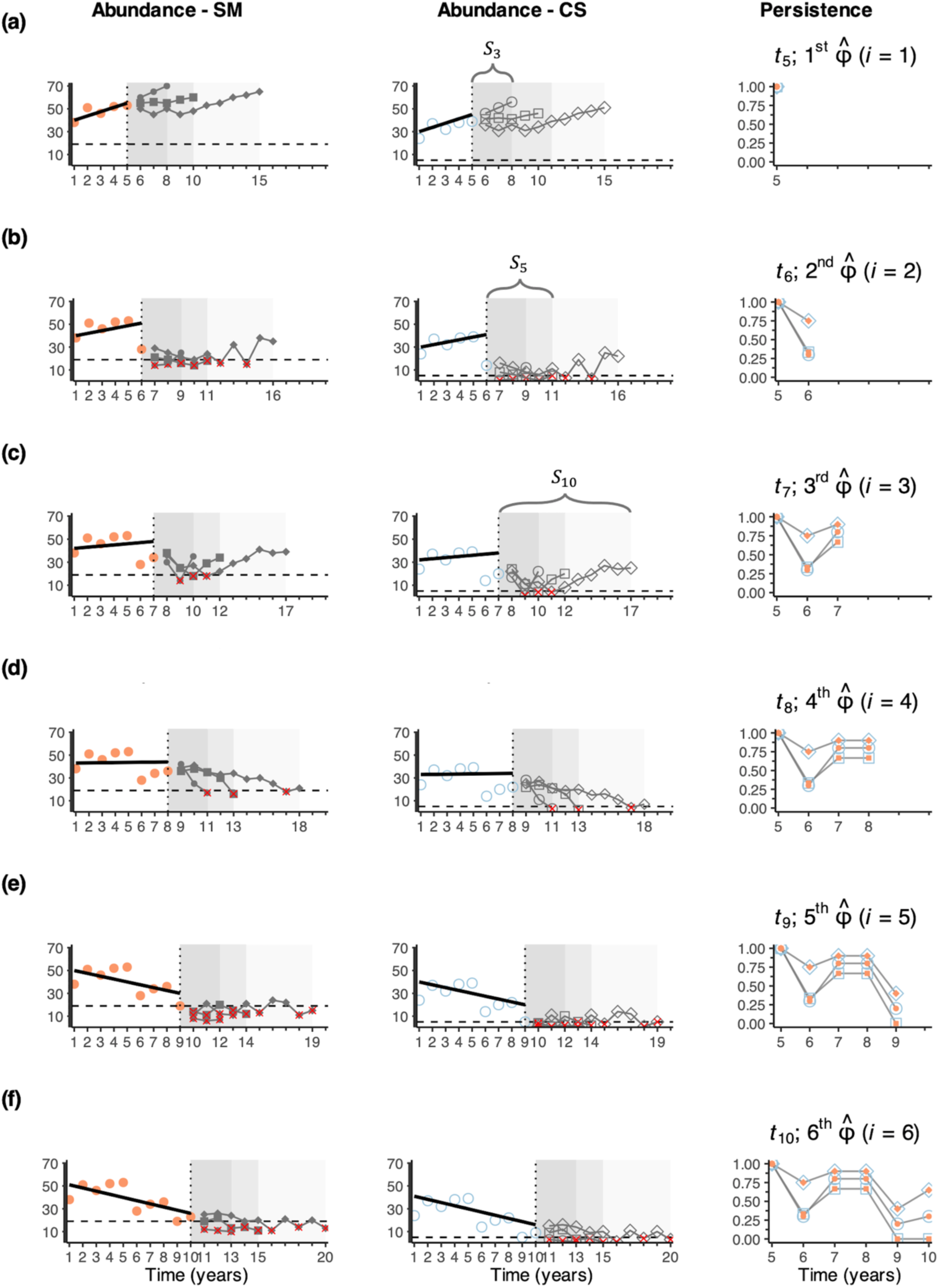
The risk-based viable population monitoring (VPM) framework for estimating local persistence probabilities over time. Observed abundance (points) informs a population model fitted from 𝑡_1_ to 𝑡_𝑖_ (black lines). Using estimated parameters (e.g., growth rate, environmental or observation noise), stochastic trajectories are simulated beyond the vertical dotted line (𝑡_𝑖_) within moving windows of 𝑆 years (gray ribbons, lines, and shapes). These simulations estimate the probability of counts falling below a quasi-extinction threshold (dashed line, red crosses); the complement is the local persistence probability (φ). The VPM iteratively updates these estimates with new data. Panels (**a-f**) show monitoring progress: abundance from standardized monitoring (SM; left), from community science data (CS; center), and persistence probability estimation φ (right). Probabilities are estimated using illustrative simulation window lengths of 𝑆_3_ (3 years, dark gray ribbon, circles), 𝑆_5_ (5 years, gray ribbon, squares), and 𝑆_10_ (10 years, light gray ribbon, diamonds) across each time with new monitoring data, 𝑖 = 1 in 𝑡_5_ (**a**) to 𝑖 = 6 in 𝑡_10_ (**f**). Simulated trajectories change between iterations, reflecting changes in persistence estimates before abundance drops below the threshold. Lower abundance estimation in CS matches the dynamics of the population in SM, resulting in similar persistence estimates (filled vs hollowed points). Extending simulations enhances the estimated robustness.

To assess the accuracy of the estimated trend in local persistence probability from applying the VPM framework to community science data, we compared our estimates using eBird data to those obtained from standardized monitoring of the endangered Everglade snail kite (*Rostrhamus sociabilis plumbeus*, snail kite hereafter; (Dreitz et al., 2002; Reichert et al., 2016)). This approach addresses major limitations in estimating population extinction risk from community science data because it can be rapidly implemented with limited computation capacity and identifies trends at multiple spatial and temporal scales, while separating observational and ecological processes. The Supplementary Information (hereafter SI) presents a detailed tutorial to allow its broad applicability.

## 2 | MATERIALS AND METHODS

Our workflow consists of five stages that we summarize here and expand in the following sections. First, we downloaded available data from eBird for our species of interest (Section 2.1), filtering the complete checklists (lists hereafter) by their sampling effort information to improve their ecological use during analysis (Johnston et al., 2021). Second, we assigned each list to a spatial grid of sampling units, extracted the maximum observed count reported in any list within the sampling unit during a specific time (e.g., weeks), and constructed the observed time series dataset from eBird (Section 2.1). Third, we applied the VPM framework (Staples et al., 2005) to estimate population viability over time, monitoring population risk with community science data (Section 2.3). This stage included the population dynamics estimation through state-space model methods (Section 2.2). Fourth, leveraging the overlapping standardized monitoring (Section 2.1), we compared estimates of persistence probability between eBird and standardized monitoring of snail kites in a subpopulation that recently expanded the species’ breeding range (Machado-Stredel et al., 2024; Poli et al., 2020). Finally, we systematically reduced the eBird dataset, reapplied the VPM framework, and compared these estimates with those from the standardized monitoring dataset. Descriptions of the modeling framework are provided below, and a detailed tutorial is available in Zenodo (Anonymous 2025) and as tutorial file (SI).

### 2.1 | Study system and time series construction

The snail kite is an endangered, wetland-dependent raptor that has been systematically monitored in Florida (Dreitz et al., 2002; Fletcher et al., 2024; Martin et al., 2007; Reichert et al., 2016). This species moves among wetlands, tracking spatially varying environmental resources that include fluctuating densities of apple snails (*Pomacea* sp.) as its main prey (Cattau et al., 2016; Machado-Stredel et al., 2024; Poli et al., 2022; Robertson et al., 2017). Systematic monitoring of snail kites includes airboat transects across all known breeding sites, nest surveys, and capture-mark-resight methods; see more details in Fletcher et al. (2024) and references therein. Such standardized monitoring data show a trend of decreases in growth rate in the southern populations and increases in the north (Reichert et al., 2016, 2021), including recent colonization of wetlands over 100 km north of its breeding range (Cattau et al., 2016, 2017; Fletcher et al., 2024; Machado-Stredel et al., 2024; Poli et al., 2020, 2022). These data are ideal to validate the accuracy and performance of our modeling approach because the standardized monitoring ran in parallel with eBird time series records, thus facilitating the comparison of estimates with the most robust monitoring data available (SI-Section 4).

We downloaded the snail kite eBird data from August 1972 (the first date available for the species in Florida) through November 2024 from https://ebird.org/data/download (SI-Section 3). To improve the ecological information included in these data for analysis, we conducted refinements through filtering that constrained the variation of sampling effort in eBird (Backstrom et al., 2024; Johnston et al., 2021; Kelling et al., 2019; Strimas-Mackey et al., 2023). We only included complete lists from "traveling" or "stationary" protocols between January 2018 and November 2024 (≥1000 lists per year, SI-Section 3.4, Figure SI-11), within a sampling effort of ≤5 hours, ≤5 km, ≤10 observers, assigning each list to a hexagonal grid of 100 km^2^ area – our spatial sampling units (Figure 2 and SI-Section 3.3).

**Figure 2.**
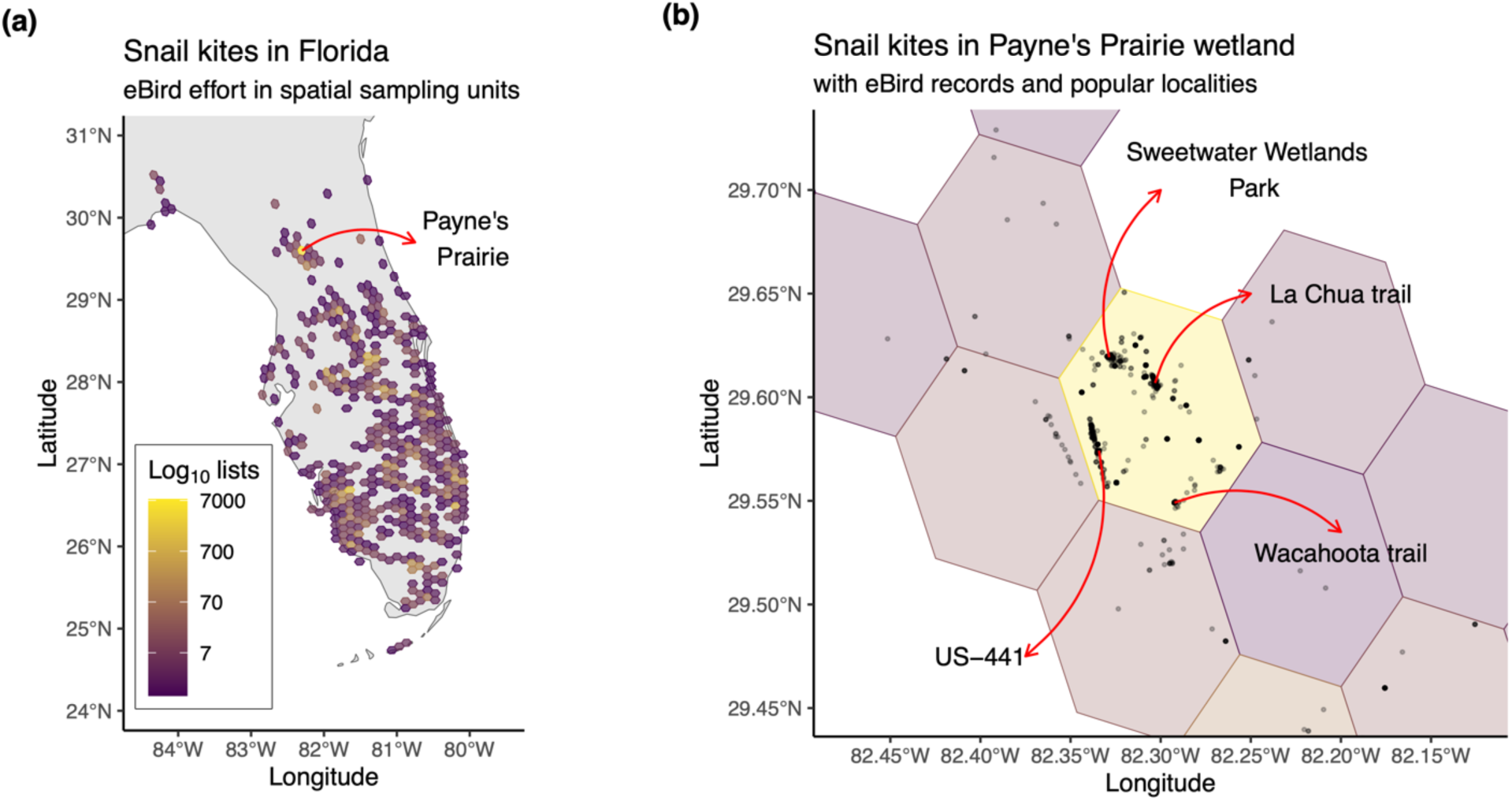
Map of spatial eBird sampling of snail kites in Florida emphasizing the hexagonal cell with the highest number of lists. **(a)** Each hexagonal cell corresponds to an area of ∼100 km^2^, and the bright color represents cells with a higher number of submitted lists to eBird (note log10 scale). **(b)** A single cell in northcentral Florida (∼Payne’s Prairie Preserve State Park, ∼85 km^2^) concentrated over 7,000 lists. eBird records of snail kites in Payne’s Prairie display some popular sites visited within this sampling unit. This sampling unit overlaps with ongoing standardized monitoring surveys.

Although several sampling units overlap with ongoing standardized monitoring surveys, we used data from a single sampling unit corresponding to the recently established population in north-central Florida: the Payne’s Prairie subpopulation in Alachua County (Figure 2). After Hurricane Irma flooded this ∼85 km^2^ wetland system in 2017, the snail kites established a nesting population in 2018 (Machado-Stredel et al., 2024; Poli et al., 2020) and have experienced an estimated steady increase in abundance (Poli et al., 2020), as inferred from eBird data from 2012-2022 (Fink, Auer, et al., 2023). This expansion motivated the inclusion of Payne’s Prairie in the standardized monitoring project of the species (Poli et al., 2020). Payne’s Prairie has 7,414 eBird lists reporting snail kites since 2018 (Figure 2). We took advantage of this overlap between high eBird observations and standardized monitoring datasets in Payne’s Prairie to compare local persistence estimates using these two data sources (see Section 2.3). We constructed the time series for the two datasets by extracting the maximum observed count reported in any list of eBird or any standardized monitoring survey per week in the sampling unit of 100 km^2^ that overlapped Payne’s Prairie. We assumed this weekly maximum count is a potential minimum number of individuals known to be alive during sampling (Poli et al., 2020).

This assumption implies homogeneous detection probability through time. Still, the state-space modeling approach addresses this issue by estimating the observation process from the two datasets separately. Moreover, to evaluate different structures of the weekly maximum counts time series, we randomly thinned the lists by sampling (list-level) and time (week-level) in our sensitivity analyses (SI-Section 7).

### 2.2 | State-Space population dynamics modelling

At the core of our modelling approach is the specification of a population dynamics model that represents the temporal fluctuation of the population of interest appropriately (represented by the straight black lines in Figure 1). Such a model should contain deterministic and stochastic components shaping the population dynamics to effectively capture and estimate population persistence probabilities (Ferguson & Ponciano, 2014). This model should also include estimation of the variability of the data due to observation error (Dennis et al., 2006; Dennis & Otten, 2000; Evans et al., 2023; Hostetler & Chandler, 2015; Ponciano et al., 2009). We chose a special case of the stochastic Gompertz state-space model (see SI for additional properties of this model (Dennis et al., 2006; Dennis & Ponciano, 2014)), which is the Exponential Growth State-Space (EGSS) model for density-independent dynamics (Humbert et al., 2009).

Let 𝑁_𝑡_ be the unobserved population abundance of the species of interest at time 𝑡. The deterministic skeleton of the ecological process is a discrete-time model using the difference equation 𝑁_𝑡_ = 𝑁_O_ exp(𝑎), where 𝑁_O_ is the initial population and 𝑎 is a positive constant representing maximum population growth rate (Humbert et al., 2009). The stochastic version including observation and environmental noise is a state-space model with unobserved and observed population abundance. The unobserved abundance is assumed to follow a stochastic process, 𝑁_𝑡_ = 𝑁_O_ exp(𝑎 + 𝐸_𝑡_), where 𝐸_𝑡_∼Normal(0, 𝜎^2^) and 𝜎^2^ is the variation of environmental noise. Let 𝑋(𝑡) be the unobserved log-abundance of the population at time 𝑡. We end with a continuous-time ordinary differential equation of a Brownian diffusion process (Humbert et al., 2009), 𝑑𝑋(𝑡_𝑖_) = (ln𝜆)𝑑𝑡_𝑖_ + 𝑑𝐵(𝑡_𝑖_), where ln𝜆 = 𝑎 is the expected change of log-abundance of the unobserved abundance (𝑋(𝑡_𝑖_)) and 𝑑𝐵(𝑡_𝑖_) is a random perturbation representing environmental variability, 𝑑𝐵(𝑡_𝑖_)∼Normal(0, 𝜎^2^𝑑𝑡_𝑖_), both occurring at one small-time unit of time 𝑡_𝑖_.

The fit of the EGSS model includes the aforementioned process, and the observation model given by 𝑌(𝑡_𝑖_) = 𝑋𝑡_𝑖_) + 𝐹_𝑖_. The observed log-abundance time series is 𝑌(𝑡_𝑖_), while 𝐹_𝑖_ is the observation error, 𝐹_𝑖_∼Normal(0, 𝜏^2^), with 𝜏^2^ being the variation of observation noise. This model allows sampling times to be discontinuous and not equally spaced (Dennis & Ponciano, 2014; Humbert et al., 2009). Thus, this density-independent model has four unknown parameters (SI-Section 2.2): ln𝜆 (the trend parameter or expected change of ln𝑁_𝑡_ in one small unit time 𝑡_𝑖_), 𝜎^2^ (variability of the process noise), 𝜏^2^ (variability of the observation noise), and 𝑥_O_(the initial log-abundance population). We compared the estimates of these parameters for the two datasets by visually inspecting the time series of the estimates and by computing the Root Mean Square Error (RMSE), where lower values indicate similarity between estimates from both datasets (SI-Section 6.3).

Although the state-space model formulates changes in abundance in discrete times of demographic dynamics (e.g., years), implementing the stochastic continuous models of diffusion process allowed us to use weekly observations as our time series data. This is possible because the discrete-time model has an exact continuous-time counterpart in the diffusion process. Specifically, the solution of the continuous-time model matches the solution of the difference equation model at discrete-time points (Dennis et al., 1991; Dennis & Ponciano, 2014; Humbert et al., 2009). Structural errors of applying a demographic year-based model to weeks have a lower effect on pattern inferences than parameter estimation errors (Taper et al., 2008). In fact, weekly estimation of local persistence from population dynamics models provides valuable insights for the management of the snail kite populations, given that wetland management (flood or drain regimes) and nest initiation rates during breeding season are typically summarized at such a temporal scale (Fletcher et al., 2021, 2024).

### 2.3 | Estimation of the local persistence probability (𝛗)

We applied a risk-based viable population monitoring framework (VPM, Staples et al. 2005; Figure 1). To estimate the unknown parameters in the state-space models, we computed restricted maximum likelihood because it improves identifiability for the multivariate normal distribution in the state-space population models (Dennis & Ponciano, 2014; Humbert et al., 2009). The risk-based VPM description recommended the simulation of population trajectories in moving windows of one, two, and five generations into the near future using annual time steps (Staples et al., 2005). Generation time estimates for snail kite range from 5-8 years (Bird et al., 2020; Cattau et al., 2017). Here, specifically, we compare trends using moving windows of 5, 10, and 25 years. Note that throughout the manuscript we also refer to the length of these moving simulation window sizes in weeks (𝑆_25O_, 𝑆_5OO_, or 𝑆_125O_, respectively), which was our temporal scale for the observation time series input and the local persistence probability estimation. Furthermore, although the specific population response to hydrology regime in the recently colonized wetland is still unknown, periodic droughts (2-10 years apart) influence snail kite population dynamics in historical locations (Beissinger, 1995); thus, these three moving window sizes can capture population responses to drought regimes at a temporal resolution that is meaningful for managers, as well as the potential rapid rates of population decline during assessment of species vulnerability to extinction (Bird et al., 2020).

Defining extinction risk during estimation of population viability is imperative for the VPM framework (Staples et al., 2005). Population extinction risk can be the probability of the population to decrease below a pre-established abundance threshold, known as quasi-extinction (Beissinger, 1995; Dennis, 2002; Dennis et al., 1991; Ginzburg et al., 1982; Staples et al., 2005; Trouillier et al., 2023). Expert opinion is often used to establish an appropriate quasi-extinction threshold. For example, a previous population viability analysis (PVA) of snail kites, at a time when population size fluctuated between 114 and 668 individuals during a twenty-year survey (uncorrected by sampling effort), used <25 individuals as a quasi-extinction threshold for the entire population in Florida (Beissinger, 1995). In contrast, the quasi-extinction threshold was set at one-half of the initial population estimates in examples within the original description of VPM (Staples et al., 2005). An appropriate quasi-extinction threshold should be low enough to estimate population risk accurately, yet high enough to avoid small population phenomena (e.g., Allee effect) or other extinction vortices that could increase the chances of extinction (Dennis et al., 1991; Dennis, 2002). Given that our counts came from two different datasets, we assumed that one-half of the observed mean counts for each dataset between January 2018 and November 2024 provides an appropriate balance for quasi-extinction; we did not run our estimates for our studied subpopulation with different values (but see SI-Section 8). Thus, we assumed that a near future population trajectory might be at extinction risk if, at some point, the predicted weekly high counts decline below a dataset-specific threshold of one-half of the observed mean; 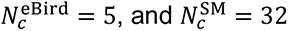. We did not use auxiliary information (i.e., same 𝑁*_c_* from literature or expert opinion, like <25 from Beissinger, 1995) because that would assume that the quasi-extinction threshold is fixed and the randomization process came from the data sampling (design-based), instead of treating the quasi-extinction threshold as a realization of a data-generating process (model-based; see Boyd et al., 2023; Williams & Brown, 2019).

To formalize our procedure of estimating persistence in each week 𝑖, we simulated 𝑀 = 50,000 trajectories in the three moving simulation windows (𝑆_25O_, 𝑆_5OO_, 𝑆_125O_). In each simulated trajectory 𝑚 = 1, 2, …, 𝑀, we defined the indicator function 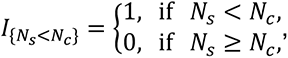, where 𝑁_𝑠_ is the simulated count for each week 𝑠 in 𝑆 after the week of persistence estimation 𝑖 within each window of simulation (𝑆_25O_, 𝑆_5OO_, 𝑆_125O_), while 𝑁_𝑐_ is the assumed threshold for quasi-extinction in each dataset. Then, for each simulated trajectory 𝑚_𝑖_ of length 𝑆, the estimated local persistence probability 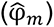 is given by 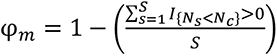. In other words, the probability of persistence at each time 𝑖 is calculated by the number of times the simulated counts fall below the critical threshold in the following moving window. The estimated probability of persistence E[φ_𝑖_] for the week 𝑖 are provided by the empirical mean, while the variability of the estimates was captured by its standard deviation and interquartile range values of the simulations. We iterated the process for all the following weeks 𝑖 of available high counts observations (𝑖 ≥ 1) in both datasets, repeating the model fitting, estimating parameters, simulating 𝑀_𝑖_ trajectories for the three moving windows of lengths 𝑆, and estimating φ_𝑖_ with its variability (SD, first, and third quartile). This iterated process was conducted for all weeks with observations between December 2019 (week 101, 𝑖 = 1) to November 2024 (week 360) for the eBird (258 weeks) and the standardized monitoring datasets (32 weeks).

The potential density-dependent and density-independent dynamics of the population can be contrasted in each time 𝑖 by fitting the continuous density-dependent dynamic model (Dennis & Ponciano, 2014), and evaluating the density-dependent parameter in the Gompertz state-space model (𝑏 ≈ 0, see SI-Section 2). We conducted this contrast initially (SI-Section 5.4), finding that density-dependent dynamic, suggested by preference of the Ornstein-Uhlenbeck diffusion process model over the Brownian diffusion process model (EGSS), was negligible for both datasets across the entire interval of persistence estimation (SI-Section 6.1, Figure SI-63).

To compare the trend of the probability of local persistence estimated from eBird (φ̂_eBird_) with that estimated from standardized monitoring data (φ̂_SM_), we followed a threefold approach. First, we visually inspected the time series of the estimates. Then, we performed linear regression and computed the associated Pearson correlation coefficient (ρ; see Figures SI- 74,80,85). We interpreted the fit of the linear models close to the 1:1 identity plot and higher values of the Pearson correlation between φ̂_eBird_ and φ̂_SM_ as indicative of similar trends of persistence probabilities estimated from the two datasets. Finally, we calculated the Root Mean Square Error (RMSE) of the absolute difference between φ̂_eBird_ and φ̂_SM_ (SI-Section 6.2). We interpreted lower RMSE values as indicative of, on average, close persistence estimates between the two time series datasets.

We conducted two separate sensitivity analyses at the temporal (weeks) and sampling (lists) levels (SI-Section 7). In both we randomly reduced 5% of the eBird data iteratively, from 95% to 5%, ending with a total of 38 eBird reduced datasets (19 temporal, 19 sampling) of weekly high counts data. Temporal thinning was applied to the eBird time series of weekly high counts described above. On the other hand, sampling thinning was applied to the original data of 7,414 complete lists, constructing different time series of weekly high counts. With these reduced datasets, we estimated φ_𝑖_ and its corresponding variability over time for the given simulation moving windows of two generations (𝑆_5OO_). As mentioned before, this window might reflect periodic droughts, every 2-10 years, affecting the population. In addition, this simulated window had the best compromise between high correlation and low RMSE between the two datasets (SI-Section 6.2). Similarly to the full eBird data comparison, we compared the relationship of φ̂_eBird_ from each eBird reduced data set with 𝛟̂_SM_through visual inspection of the resulting time series, performing linear regressions with the corresponding Pearson correlation coefficient, and calculating the RMSE. Again, we interpreted a fit of the linear model close to the 1:1 identity plot, higher values of Pearson coefficient, and lower RMSE values as validation indicators that the trend of φ̂_eBird_ is similar to φ̂_SM_.

All our analyses were conducted using the R software v4.3.3 (R Core Team, 2024). Specifically, we filtered the eBird data with the packages *auk* (Strimas-Mackey et al., 2018) and *tidyverse* (Wickham et al., 2019), managing dates with *lubridate* (De Valpine et al., 2017). We generated the hexagonal grid of spatiotemporal subsampling with the package *dggridR* (Barnes, 2023). We also adjusted the R scripts published by Humbert et al. (2009) and Dennis & Ponciano (2014), which require the package *MASS* (Venables & Ripley, 2002). In addition to figures using basic R or *ggplot2* (Wickham, 2016) within *tidyverse*, we used the packages *maps* (Becker et al., 2022) and *gridExtra* (Auguie, 2017) to load maps and generate composite figures, respectively, and *ggpubr* (Kassambara, 2023) to include some linear equations within the figures.

## 3 | RESULTS

### 3.1 | Predicted abundance and model parameters estimates

The predicted weekly high counts of snail kites in Payne’s Prairie, using independent EGSS models, closely matched the observed high counts in both eBird and standardized monitoring datasets (Figure 3). The predicted trajectory from standardized monitoring was highly correlated with the observed weekly high counts (linear model; intercept: −0.63 ± 1.08 SE; slope = 0.929 ± 0.030 SE, *p* < 0.001; Figures 3, SI-23). The EGSS model for eBird data predicted lower weekly high counts than standardized methods (linear model; intercept: 2.34 ± 0.79 SE; slope = 0.074 ± 0.022, *p* = 0.004; Figures 3). Nonetheless, the estimated parameters for environmental noise (𝜎e^2^ ≅ 0.03) and initial population size (weekly high count at 𝑡 = 1; 𝑒^𝑥̂0^ ≅ 1) were identical between the two datasets. The expected change in log-abundance per week showed a lower increase in eBird 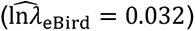 and a higher change in standardized monitoring weekly counts 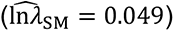. Unsurprisingly, observation noise was about an order of magnitude greater in the eBird data 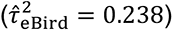 compared to the standardized monitoring data 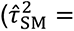0.027). We expected this strong difference due to the higher variation of the sampling effort in eBird compared to standardized surveys. This pattern stands across the 32 weeks of estimation with overlap between the datasets (SI-Section 6.3), with same estimation between datasets for initial population (𝑒^𝑥̂0^), close estimation for environmental noise (𝜎̂^2^) and trend parameters 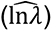, while the higher difference for observation noise estimates (𝜏̂^2^; Table 1, SI-Section 6.3).

**Figure 3.**
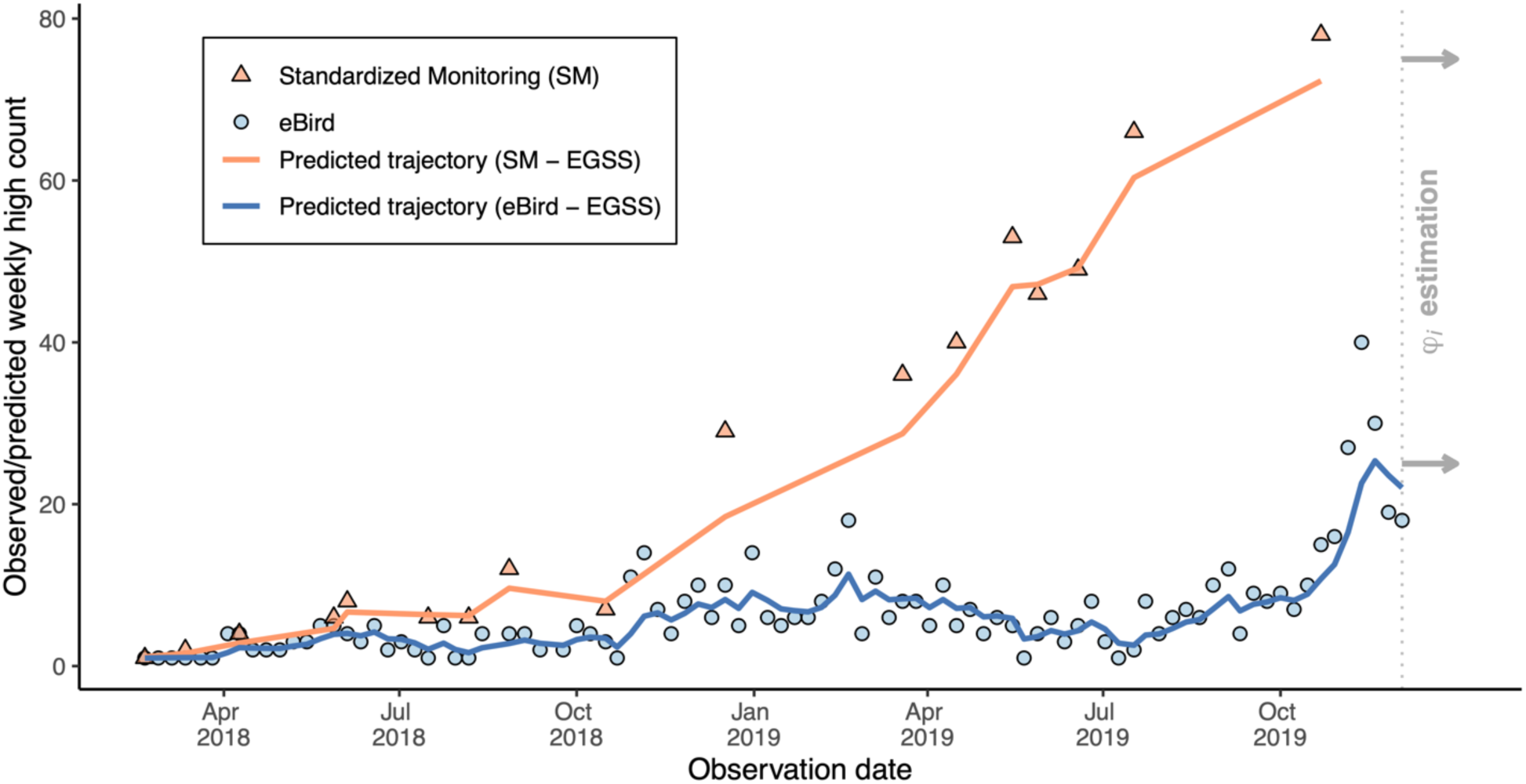
Weekly high counts observed (triangles and points) and predicted (solid lines) of snail kites in Payne’s Prairie from standardized monitoring (orange) and eBird (blue). Predicted values correspond to exponential growth state-space (EGSS) models fitted for the two datasets during the initial population dynamic in north-central Florida (from week 1, January 2018 to week 101, December 2019, which was the first week of local persistence estimation, dotted vertical line and gray arrows). Although eBird underpredicts the weekly counts, the estimated population parameters were very similar between the two datasets (SI-Section 6.3), with the main difference being higher observation noise in the models fitted with eBird data compared to the models fitted with standardized monitoring data (Table 1).

**Table 1.** Restricted maximum likelihood estimates for the parameters in the exponential growth state-space models used to estimate the local probability of persistence from two datasets.

| Parameter | Standardized monitoring | eBird | RMSE |
| --- | --- | --- | --- |
| Initial population ($e^{x_0}$) | ~1 | ~1 | 0 |
| Environmental noise ($\hat{\sigma}^2$) | 0.04<br>(0.02, 0.06) | 0.04<br>(0.03, 0.05) | 0.001 |
| Trend ($\widehat{\ln \lambda}$) | 0.02<br>(0.002, 0.049) | 0.01<br>(0.005, 0.032) | 0.009 |
| Observation noise ($\hat{\tau}^2$) | 0.02<br>($9.2 \times 10^{-7}$, 0.03) | 0.22<br>(0.19, 0.24) | 0.201 |

Note: Mean value estimated for 32 overlapping weeks between December 2019 and November 2024.
The ranges are shown in the parentheses. Time series showing each week's estimation are in the

### 3.2 | Estimated local persistence probability (𝛗̂)

Local persistence probability (φ̂) estimated from eBird data revealed a similar overall decreasing pattern to that estimated from the standardized monitoring data (Figure 4). The correlation between the two estimates was strongest when the population was projected one generation ahead (𝑆_25O_, Pearson ρ = 0.959; Figure 4a), slightly lower for two generations (𝑆_5OO_, ρ = 0.934; Figure 4b), and weakest for five generations (𝑆_125O_, ρ = 0.903; Figure 4c). However, the RMSE was lowest for 𝑆_125O_ (ρ = 0.005) compared to 𝑆_5OO_ (ρ = 0.039) or 𝑆_25O_ (ρ = 0.103), indicating closer agreement between the datasets at longer projection intervals (SI-Section 6.2). The decreasing trend through time in φ̂ was consistent among datasets and simulation windows during the monitoring of the population extinction risk, though overall lower φ̂ for shorter than longer windows (compare Figure 4a and 4c). This might be related to stochastic simulated trajectories length, with a higher proportion of simulated weeks ending below the quasi-extinction threshold, resulting in overall lower φ̂. The higher temporal resolution of eBird data allowed for the estimation of φ̂ when data from the standardized monitoring was lacking. This lack of data occurred because the prairie was too dry to conduct airboat surveys.

**Figure 4.**
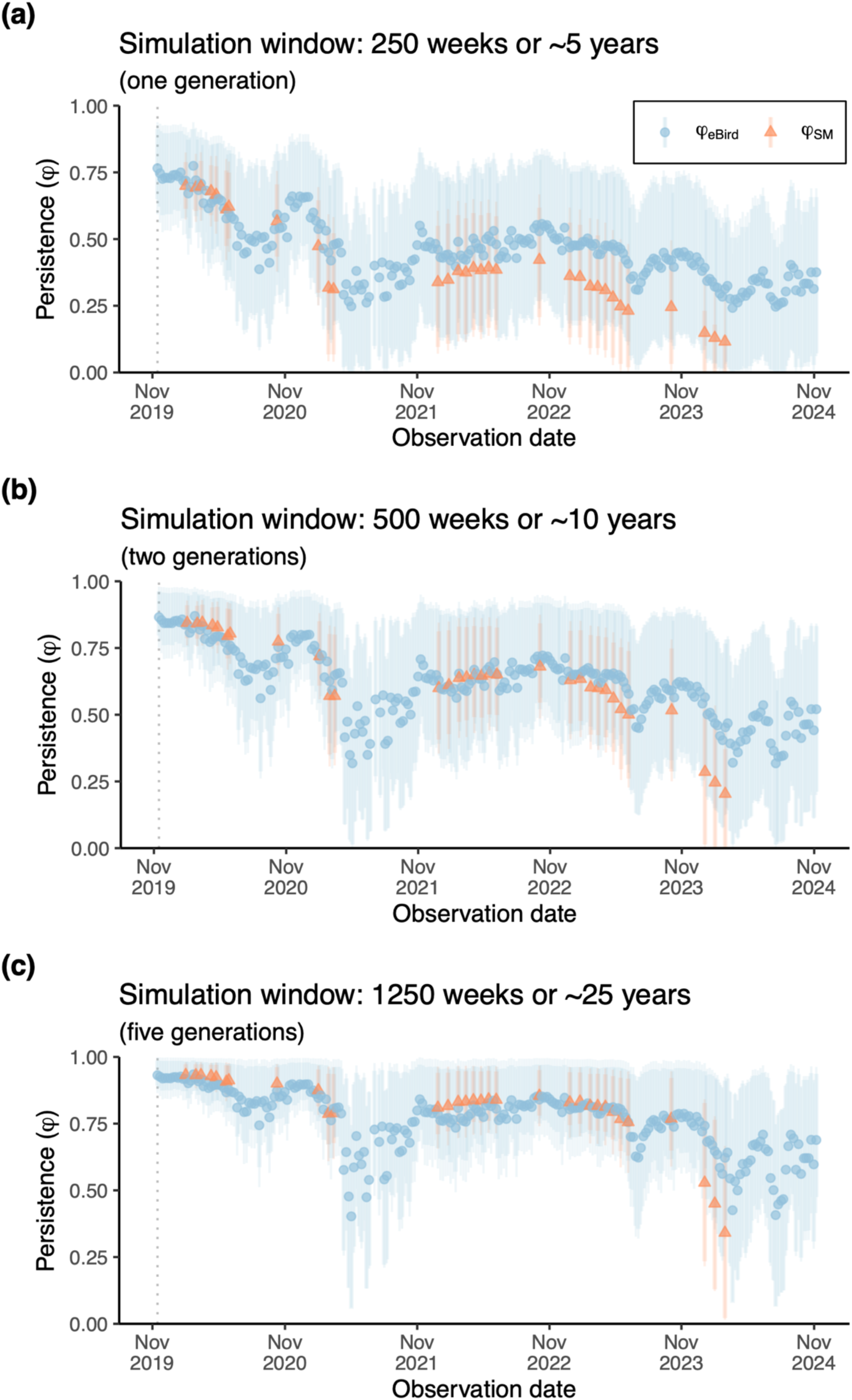
Estimated local persistence probability (φ̂) during five-years of monitoring (December 2019 to November 2024) after simulation trajectories for three moving windows (**a**: 𝑆_250_, **b**: 𝑆_500_, **c**: 𝑆_1250_). For each week of estimation, a density-independence model (EGSS) was fitted for two observation time series datasets, estimating parameters, simulating 50,000 trajectories, and estimating local persistence probability. Variation in φ̂ (points) is reflected by the ± one standard deviation (thin light line ranges) and the first and third quartiles (coarse line ranges). Higher temporal resolution is evident in eBird (*n* = 258) when compared with standardized monitored data (*n* = 32). See extended figures SI-69 to SI-85 (SI- Sections 6.4 to 6.6) with the observed high counts time series and the figure SI-63 (SI-Section 6.1) for the inclusion of potential density-dependent dynamics. The declining pattern between φ̂ from standardized monitoring and eBird datasets is highly correlated (ρ > 0.9), although the best compromise between high correlation and low RMSE was identified in a simulation window of two generations, 10 years (𝑆_500_).

### 3.3 | Sensitivity analyses to varying time series length and sampling

Overall, the correlation in the trend from the two data sources was consistently above ρ ≥ 0.90 with a temporal reduced datasets of 95-30% (about 47 to 18 weeks per year) and 20% (10 weeks per year) of the eBird high counts data (Figures 5, SI-88, SI-90). Similarly, the correlation in the trend was also above ρ ≥ 0.90 with a sampling reduced datasets of 95-50% (Figures 5, SI-94, SI-96). Moderate to high correlation, 0.65 ≤ ρ < 0.90, was detected with a temporal reduced dataset of 25% (13 weeks per year) and 15% (8 weeks per year), and a sampling reduced dataset of 45-25% (51-48 weeks or 670-370 lists per year). Few overlapping weeks between the datasets limited the correlation estimates in a temporal reduced datasets of 10-5%. A sampling reduction to the 20-5% of eBird (297 to 74 lists per year) still had a positive correlation (ρ ≥ 0.50; Figures 5, SI-94, SI-96). Remarkably, the declining trend of φ̂ was still visually evident even with 5% of the data retained, either at temporal or sampling level, with all reduced datasets having RMSE >0.18 (Figure 5, SI-Section 7). These results suggest that the framework is particularly robust and provides meaningful estimates of the probability of persistence even when available data is significantly reduced.

**Figure 5.**
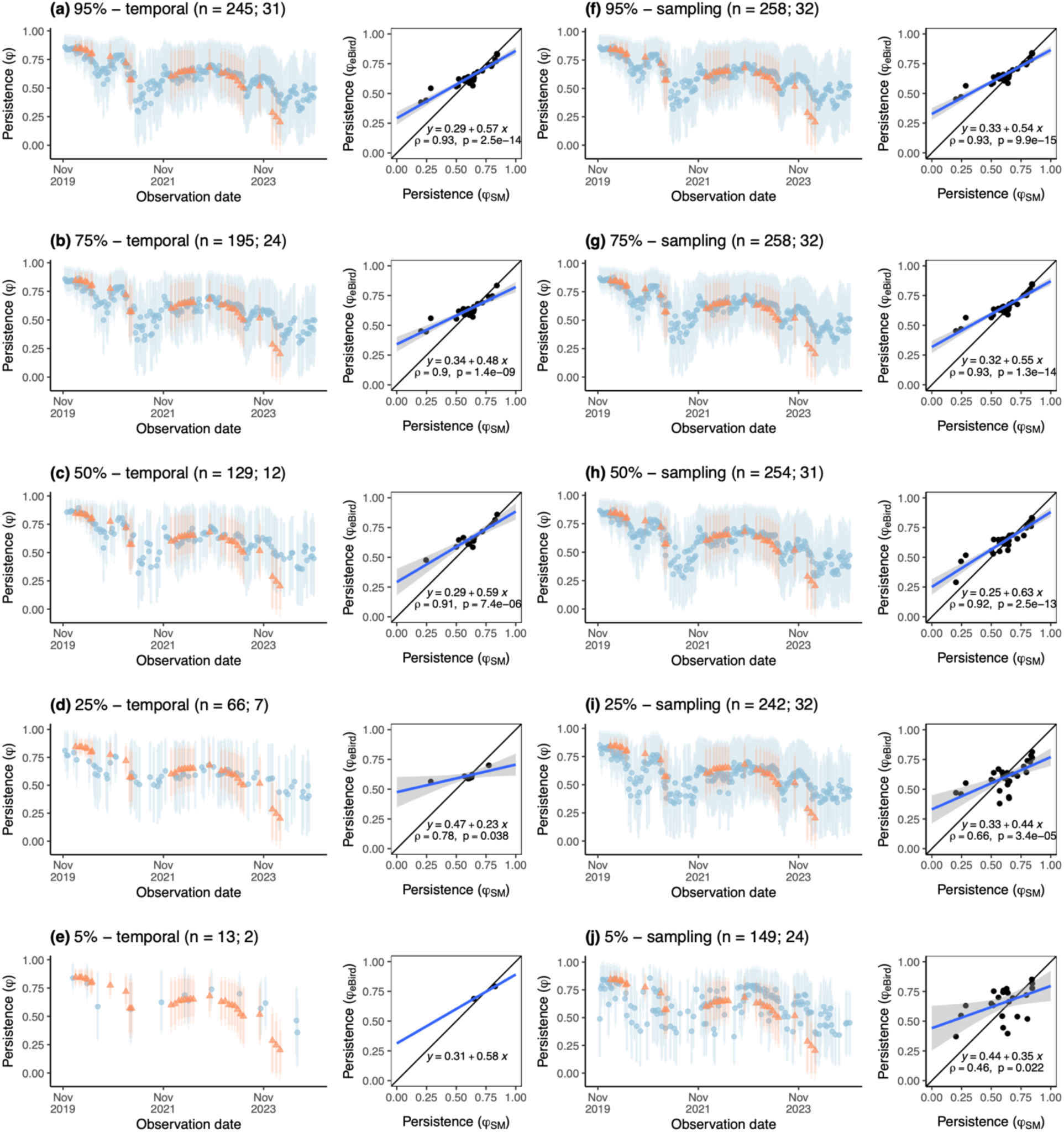
Data reduction as sensitivity analyses. Estimation of local persistence probability (φ̂) from standardized monitoring data (orange; φ̂_SM_) is replicated in each comparison with estimation of local persistence probability from eBird (blue; φ̂_eBird_) under temporal (left) and sampling (right) data reduction. The sequence of panels represents reduction from the complete eBird dataset (Figure 4c) to different percentages of randomly sampled data by sampling (**a-e**; see extended figures in SI-Section 7.1.1) and temporal thinning (**f-j**; see extended figures in SI-Section 7.1.2). The *n* in parentheses indicates the number of weeks from eBird reduced data and overlapping benchmark weeks. Trends of φ̂ are accompanied with the relationship of φ̂_eBird_ (ordinate) and φ̂_SM_ (abscissa). A linear model of this relationship is fitted and deployed with a blue line, including confidence intervals in gray ribbon (ρ below the equations shows Pearson correlation coefficient); the diagonal line is the line of identity (1:1).

Although model accuracy varies with decreasing data and some overestimation of φ̂_eBird_ compared to φ̂_SM_ suggests less conservative estimation, the trends are still similar even with as little as 5% of the original eBird weekly high counts (about three weeks or 74 lists per year).

## 4 | DISCUSSION

We show that a risk-based viable population monitoring (VPM) approach is a promising method for estimating local persistence probability over time using community science data. We compared local persistence estimates from eBird with those from standardized monitoring using an endangered wetland raptor as a case study. Both datasets revealed remarkably similar trends, even when eBird weekly observed counts were reduced from 258 to 13 weeks (around 5% of the original dataset) or from 7,714 to 385 lists across the 5 years of monitoring (Figures 4, 5). While persistence estimates using eBird were higher when compared to those estimated from standardized monitoring at lower values, our modeling approach effectively assessed population extinction risk over time.

The similarity in persistence estimates between the two datasets stems from applying continuous stochastic state-space models within the VPM framework, which offers flexibility in representing ecological processes. These models estimate population parameters using time series of uneven time intervals (Dennis & Ponciano, 2014; Humbert et al., 2009), as in our datasets of weekly counts (SI-Section 7), and assess the risk of declining below a quasi-extinction threshold for each period with data. This combined flexibility avoids reliance on arbitrary thresholds and directly quantifies weekly persistence probability from a model-based approach (Boyd et al., 2023; Williams & Brown, 2019). Defining population-size thresholds is a common issue in PVA applications, which has been solved using a percentage of either the carrying capacity (Ginzburg et al., 1982) or the initial population (Taper et al., 2008; Trouillier et al., 2023). We used one-half of the observed mean counts for each dataset, but other percentages of population size can be applied with our method (SI-Section 8; Dennis, 2002; Ferguson & Ponciano, 2014; Trouillier et al., 2023).

Despite underpredicting weekly counts compared to standardized monitoring (Figures 3, SI-23), eBird data captured population risk trends reliably (Table 1, Figure 4). Underprediction arises from spatial coverage bias (Royle et al., 2007) and observation noise during sampling (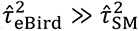; SI-Section 6.3; Poli et al. 2020). eBird observers are sampling only a portion of the population surveyed by standardized methods (Poli et al., 2020; SI-Section 4). The observers’ expertise in eBird is also directly related to detection probability (Fink et al., 2020; Johnston et al., 2018). Taken together, counts submitted to the platform likely do not represent all individuals available to be sampled in the population. These biases, however, are less critical for assessing population risk than for estimating absolute abundance because persistence probability estimation uses population process parameters to simulate future trajectories rather than analyze direct change in abundance prediction under confounding observation error. The similar trend, high correlation, and lower RMSE in risk predictions between datasets (Figures 4, 5) further supports the robustness of our approach (Williams & Brown, 2019).

Lower persistence probabilities in our case study do not necessarily equate to local extirpation. Snail kites in Florida exhibit spatially structured dynamics moving among wetlands in response to environmental variability (Martin et al., 2007; Poli et al., 2022; Reichert et al., 2016). This spatial structuring may explain why none of the local persistence probability estimates from systematic surveys identified density-dependent dynamics (SI-Figure SI-47). Density-dependence may act on broader spatiotemporal scales or be less relevant for an expanding population (Phillips, 2009), as is the case of snail kites in Payne’s Prairie – likely not reaching carrying capacity yet (Poli et al., 2020). Our findings for the eBird dataset align with density-independent dynamics observed in this population (Figure SI-63), matching the estimates from this community science platform with those from standardized monitoring data (Figures 4, 5).

Our approach can be extended to incorporate spatial structure, density dependence, biotic interactions, and environmental variables to capture nuanced dynamics in a population (Dennis & Otten, 2000; Ferguson et al., 2017; Ives et al., 2003).

Current methods using eBird data to estimate population trends focus on encounter rates during breeding seasons or unidirectional change in relative abundance (Fink et al., 2020; Fink, Johnston, et al., 2023; Horns et al., 2018; Neate-Clegg et al., 2020). In contrast, we derived weekly estimates of population persistence following a monitoring viability approach (Bakker & Doak, 2009; Staples et al., 2005), applying continuous state-space models to estimate parameters and simulate trajectories over temporal windows relevant to management (Fletcher et al., 2021, 2024; Staples et al., 2005). Our approach likely captures population dynamics that are more sensitive to environmental change. For instance, while the snail kite eBird status and trends data products suggest a population increase in north-central Florida (Fink, Auer, et al., 2023), our models reveal a decline in local persistence from the two overlapping datasets (Figure 4). This discrepancy likely reflects seasonal and post-colonization dynamics (Phillips, 2009; Yackulic et al., 2015), which seems overlooked by the discrete models used in the eBird status and trends products (Fink, Johnston, et al., 2023). Although, on average, eBird shows a rise in abundance (2012: 0 individuals, 2022: ∼14) in Payne’s Prairie, which is also evident in the standardized monitored counts (2012: 0, 2022: ∼100), our models provide insights into the nuanced dynamics of the population trend estimating a probabilistic measure of viability through iterative count-based PVAs (Bakker & Doak, 2009; Staples et al., 2005). The declining trend likely reflects snail kite response to environmental variables such as hydrology (Ferguson et al., 2017; Fletcher et al., 2021; Robertson et al., 2017), among other common population dynamics during range expansion shifts (Phillips, 2009; Yackulic et al., 2015). Focusing on local persistence probabilities derived from continuous state-space models, our approach complements existing methods by offering a risk-based framework for population monitoring using community science data. This can be used by researchers and managers to identify new sites or providing early warnings of population change in unmonitored subpopulations and prioritize further standardized monitoring (SI-Section 8; Anonymous 2025).

Integrated population models combining community science data with other demographic approaches improve abundance and temporal dynamics estimates (Fletcher et al., 2019; Stillman et al., 2023; Zhao et al., 2024). The problem arises when standardized monitoring is scarce or non-existent. Moreover, some advanced quantitative approaches, while useful (e.g., Fink, Johnston, et al., 2023; Johnston et al., 2025; Stillman et al., 2023), are computationally intensive and, hence, less accessible to a wider audience of ecologists that may lack the computational resources to apply them (Johnston et al., 2023). Our approach addresses these issues by providing an easy-to-use practical tool to assess persistence and population decline risks over time (i.e., monitoring extinction risk). Thus, it is potentially extendible where standardized monitoring is still absent but community science is available, enabling adaptive management during monitoring (Bakker & Doak, 2009; Staples et al., 2005). Our two sensitivity analyses (SI-Section 7) confirmed the robustness of our approach when applied to different scenarios like lower counts (by sampling thinning) or lower weeks with data (temporal thinning), fostering potential transferability to other species.

Our approach can track population trends and accurately estimate persistence probabilities, even if eBird counts are underpredicted. The method could be an excellent tool for near-term forecasting of population extinction risk at the local spatial scale, but it is not relevant for understanding the demographic mechanism driving the population viability. This means that although useful, this approach does not replace long-term standardized monitoring for population conservation; instead, it will provide a complementary tool. Standardized biodiversity monitoring programs are still exceptions rather than the rule globally, thus leverage on this readily available community science data is a great opportunity. eBird, particularly, includes over 10,000 bird species worldwide, a monthly release of data, and an increasing trend of observers’ participation. Our modeling framework provides a computationally efficient, fast, and easy-to-use capability (see the tutorial in the SI). This is especially useful in places where standardized monitoring needs reinforcement, and community science observations are already common and increasing.

## AUTHOR CONTRIBUTIONS

Orlando Acevedo-Charry, José Miguel Ponciano, and Miguel A. Acevedo conceived the ideas, Caroline L. Poli, Brian M. Jeffery, Robert J. Fletcher Jr., María Ángela Echeverry-Galvis, Bette A. Loiselle and Scott K. Robinson provided feedback to designing the study. Caroline L. Poli, Brian M. Jeffery, Scott K. Robinson, José Miguel Ponciano, Orlando Acevedo-Charry, Miguel A. Acevedo, and Robert J. Fletcher Jr. collected the data; Orlando Acevedo-Charry and José Miguel Ponciano analyzed the data, and interpretation was conducted by all authors; Orlando Acevedo-Charry, Miguel A. Acevedo, and José Miguel Ponciano led the writing of the manuscript. All authors contributed critically to the draft and gave final approval for publication.

### Statement of inclusion

Our study brings together authors from several different countries (Colombia, Guatemala, USA), including scientists based in various regions. All authors were engaged early on with the research and study design to ensure that the diverse perspectives they represent were considered from the onset. Whenever relevant, literature published by scientists from the underrepresented regions was cited. Unsuccessful efforts were made to consider relevant work published in a language different than English; we are planning to address this caveat in future research by submitting the application used in the present study in journals in Spanish language. We begin this effort including the Abstract in Spanish, Portuguese, and Chinese (simplified) for the online version of the article and plan to propose workshops in academic meetings and across institutions.

## DATA AVAILABILITY STATEMENT

eBird data is freely available (under request) on the website https://ebird.org/science/use-ebird-data. The R language source code with data for the eBird example and standardized benchmark, as well as a full tutorial describing the procedure to estimate persistence probability in the Supplementary Information is available in Zenodo https://doi.org/10.5281/zenodo.15475635 (Acevedo-Charry et al., 2025).

## ACKNOWLEDGEMENTS

We are in debt to the many volunteers reporting observations under good practices or reviewing eBird data to maintain quality of the platform; they are the underlying force that provides support to “the power of the people” by community, participatory, or citizen science data. Orlando Acevedo-Charry received departmental funding at the University of Florida, such as the College of Agricultural and Life Science (CALS) Dean’s Award, the Doris and Earl Lowe and Verna Lowe CALS Scholarship, and the School of Natural Resources and Environment Robin E. Nadeau Graduate Research Award. Snail kite monitoring was implemented under IACUC#202008334 and received funding from the US Army Corps of Engineers (W912HZ-20-2-0033), South Florida Water Management District (9500008729), Florida Fish and Wildlife Conservation Commission (13416), and the St John’s River Water Management District (34941). Talita Ribeiro and Min Zhao provided corrections to the Portuguese and Chinese abstracts, respectively.

Jackson Martin and Jay Kalogiros-Pepper locked in on providing comments to improve the manuscript. We also thank two anonymous reviewers whose comments improved the manuscript. We acknowledge that the work described in this article was carried out on the ancestral territory of the late Potano (Timucua) and Seminole (Alachua Seminole) people.

