## Supplementary material for "Monitoring population extinction risk with community science data": vpm_eBird-to-html.html

###### Authors…(removed for peer review in Journal of Applied Ecology)

###### 20 May 2025

### 1 Read me (begin here)

This file contains the code used to support the paper to . We hope this code serves as a practical tutorial and is applied for different users in an intuitive way. Our aim is to describe and test a quantitative approach that fits continuous state-space models iteratively to a time series of community science data (eBird), with the ultimate goal of estimating local persistence probability through time. We evaluated model accuracy by comparing estimates and trends using eBird data with those estimated from the endangered Everglade’s snail kite long-term standardized monitoring project. We also performed a sensitivity analysis to assess how robust the persistence estimates are to a reduction in the number of eBird observations available. We used the risk-based viable population monitoring (VPM) framework (Staples *et al.*, 2005), fitting continuous state-space population models under density-independent dynamics (Humbert *et al.*, 2009).

This population dynamic model can also be extended to density-dependent dynamics (Dennis & Ponciano, 2014; code provided). We focused on the recent (since 2018) expanded population of Everglade’s snail kite (*Rostrhamus sociabilis plumbeus*) in north-central Florida, US, a species with standardized monitoring efforts that provide benchmark population trends for comparison (see Section 2.1 in the main text).

Two continuous versions of the discrete-time equal sampling Gompertz State-Space model can be fitted with our quantitative approach:

1. The density-dependent Ornstein-Uhlenbeck State-Space model (OUSS)
2. The density-independent Exponential Growth State Space model (EGSS)

We present the risk-based viable population monitoring framework (see
Staples *et al.*, 2005),
estimating the probability of local persistence \(\phi\);
1-probability of crashing abundance below a specified threshold given a simulation window in near future.

The statistical properties for our proceedings are based on Dennis *et al.* (1991),
Dennis *et al.* (2006),
Dennis & Ponciano, (2014),
Humbert *et al.*, (2009),
and reference therein. We adjusted the functions and code published as
supplementary information in Humbert *et al.*, (2009)
and Dennis & Ponciano, (2014).
Thus, users should first run all the packages and functions in the next
section. Then, access eBird data and organize it as time series in spatiotemporal subsamples and estimate local persistence - monitoring population risk with community science data.

### 2 Packages, models, and functions

#### 2.1 Packages required

```
#R functions and datasets to support "Modern Applied Statistics with S", 
  #a book from W.N. Venables and B.D. Ripley
  library(MASS);
#Kernel Density Estimation
  library(kde1d)
#To conduct eBird data filtering and manipulation (see Strimas et al. 2018 and 2023)
  library(auk); 
#To data management and visualization - sevent packages in one
  library(tidyverse)
#To conduct Spatiotemporal Subsampling
  library(dggridR)
#Simple features to encode spatial vector data
  library(sf)
#load maps
  library(maps); 
#composite figure
  library(gridExtra); 
#equation in figure with panels (`ggplot2::ggplot()`; `facet_wrap()`)
  library(ggpubr);
```

#### 2.2 Special cases of the Gompertz state-space population dynamics model

We provide the rationale to use two special cases (continuous, diffusion process) models of the discrete-time Gompertz State Space model (Dennis *et al.*, 2006); the density-dependent version that we heretofore denotes as the GSS model and the density-independent model (EGSS) from Humbert *et al.* (2009).

Let \(N\_t\) be the latent unobserved abundance population at time \(t\), the ecological process in the GSS model. This abundance is defined as \(N\_t = N\_{t-1} ~e^{a+b\*\ln{N\_{t-1}} + E\_t}\)), where \(a\) and \(b\) are constants representing population growth rate and strength of density dependence, respectively, and \(E\_t\) is the environmental stochasticity or process noise, \(E\_t\sim \text{Normal}(0,\sigma^2)\) (Dennis *et al.*, 2006). On the logarithmic scale (\(X\_t = \ln{⁡N\_t}\)), the GSS becomes linear and follows an autoregressive model of order 1: \(X\_t = X\_{t-1} + a + b\*X\_{t-1} + E\_t\), which can be simplify as \(X\_t = a + c\*X\_{t-1} + E\_t\), where \(c = b+1\) is a constant that represents the strength of density dependence (Dennis *et al.*, 2006; Ponciano *et al.*, 2018; Reddingius, 1971 in Acta Biotheor, 20, 1-208). If \(b=0\) (\(c=1\)), the GSS reduces to the density-independent model fully treated in state-space model form by Humbert *et al.* (2009).

The discrete GSS population model has four unknown parameters: \(a\), \(c\), \(\sigma^2\), and \(\tau^2\) (Dennis *et al.*, 2006). The transition probability distribution of this logarithmic abundance model is normal, with mean and variance changing as a function of time. The parameter \(c\) represents the strength of density-dependence (Ponciano *et al.*, 2018). If the strength of density dependence (\(c\)) ranges \(-1<c<1\), the long-run probability distribution of log-abundance approaches a time-independent normal stationary distribution (\(X\_t→X\_∞\)), with mean \(\frac{a}{1-c}\) and variance \(\frac{\sigma^2}{1-c^2}\). Thus, instead of approaching a single population abundance value, or a deterministic carrying capacity, the density-dependent stochastic GSS model approaches a stationary distribution, a cloud of points around which the population fluctuates (Dennis & Taper, 1994; Wolda, 1989). The mean of this distribution, \(\frac{a}{1-c}\), represents a long-term expected population size (Dennis *et al.*, 2006). As mentioned above, the density-independent stochastic exponential growth model of Dennis *et al.* (1991) is attained when \(b=0\) (\(c=1\)); the state equation then becomes \(N\_t = N\_{t-1} ~e^{a + E\_t}\) (Dennis *et al.*, 1991). The state-space model formulation for these two models is completed with the specification of the sampling (observation) error model. Following Dennis *et al.* (2006), we assumed that the log-observation at time \(t\), \(Y\_t\), is a sample from the (stochastic) process model according to the equation \(Y\_t = X\_t + F\_t\); where the \(F\_t\) are independent and identically distributed normal random variables, i.e. \(F\_t \sim \text{Normal}(0,\tau^2)\). During transition growth dynamics, the EGSS model may more accurately represents population trends (but see Dennis & Ponciano, 2014 for details in non-stationary continuous GSS).

Fitting both state-space models when equally sampled population abundances are available is straightforward (Dennis *et al.*, 2006), yet unequally sampled populations over time tend to be the norm rather than the exception in ecology (Dennis et al., 2010). By exploiting the connection of the EGSS and GSS models to diffusion processes, here we show how to connect these models to unequally sampled data to later extend inferences to the dynamic of local persistence (as mathematical complement of local quasi-extinction risk). Indeed, the logarithmic transformation in the GSS discrete model opens the opportunity for estimating the infinitesimal mean and variance under diffusion processes for unequal sampling; Brownian motion diffusion in the EGSS model (Humbert *et al.*, 2009) and Ornstein-Uhlenbeck diffusion in the GSS model (Dennis & Ponciano, 2014). Specific statistical properties can be found in the following section with the specific functions per model.

#### 2.3 Functions

The following functions are required to manipulate data and fit the
models correctly.

##### 2.3.1 Miscelaneous functions

Function to convert time observation to hours since midnight in eBird
data

```
time_to_decimal <- function(x) {
  x <- hms(x, quiet = TRUE)
  hour(x) + minute(x) / 60 + second(x) / 3600
}
```

Multivariate normal random number generator - State Space models

```
randmvn <- function(n, mu.vec, cov.mat){

  # Save the length of the mean vector of the multivariate normal distribution to sample
  p         <- length(mu.vec);
  # The Cholesky decomposition 
    #(factorization of a real symmetric positive-definite sqr matriz)
  Tau       <- chol(cov.mat, pivot=TRUE);
  # generate normal deviates outside loop
  Zmat      <- matrix(rnorm(n=p*n,mean=0,sd=1),nrow=p,ncol=n);

  # empty matrix
  out       <- matrix(0,nrow=p,ncol=n);
  # iterate
  for(i in 1:n){
    Z       <- Zmat[,i];
    out[,i] <- t(Tau)%*%Z + mu.vec
  }

  return(out)
}
```

Function to generate equation of a simple linear model in the figures

```
  lm_eqn <- function(df, x, y){
    m <- lm(y ~ x, df);
    eq <- substitute(italic(y) == a + b %.% italic(x)*","~~italic(R)^2~"="~r2, 
         list(a = format(unname(coef(m)[1]), digits = 2),
              b = format(unname(coef(m)[2]), digits = 2),
             r2 = format(summary(m)$r.squared, digits = 3)))
    as.character(as.expression(eq));
}
```

##### 2.3.2 Functions for the Brownian diffusion EGSS model

The stochastic exponential growth model serve as a null hypothesis of density-dependence models (Dennis *et al.*, 2006). The exponential stochastic process for \(N\_t\) could be defined as \(N\_t = N\_0 \lambda^t + E\_t\), where \(λ^t\) expresses the finite rate of change (\(\lambda^t = e^{a(t)}\)); \(a\) being the instantaneous, intrinsic, or maximum per capita rate of change), \(N\_0\) is the initial population (\(N(0)\)), and \(E\_t\) is the environmental stochasticity or process noise at time \(t\), which follows a normal distribution with mean \(0\) and variance \(\sigma^2\) (\(E\_t\sim\text{Normal}(0,\sigma^2)\)). On the logarithmic scale (\(X\_t=\text{ln}(N\_t)\)), the discrete-time process can be defined as a continuous Brownian diffusion process with the stochastic differential equation:

\[
\begin{array}{ccc}
dX(t) &=& (\ln \lambda) dt+\beta dW(t) \\
&=& (a)dt+\beta dW(t)
\end{array}
\]

where \(X(t) = \ln{N(t)}\) or the log-normal population abundance at
time \(t\), \(a = \ln{\lambda}\) is a constant of
population growth rate (\(\ln{\lambda}-(\frac{\sigma^2}{2})\) as in the
***Eq13*** in Dennis *et al.* (1991);
named \(\mu = \ln{\lambda}\) in Humbert *et al.* (2009),
although this notation is confusing for the OUSS notation), and \(dW(t)\) is a random perturbation representing the environmental stochasticity or process
noise (Itô log-transformation of \(E\_t\) from GSS in Dennis *et al.* (2006)), with mean \(0\) and variance \(\sigma^2dt\), or the intensity of environmental noise scaled by \(\beta\), with \(\beta>0\) (Humbert *et al.*, 2009).

The EGSS model includes a component in sampling times \(t\_i\) not equally spaced. Thus, we denoted the latent realized abundance (e.g., weekly high counts in our case) as \(n(0), n(t\_1), n(t\_2), ..., n(t\_q)\). Let \(x(t) = \ln{n(t)}\) and let \(Y(t\_i)\) be a value of \(x(t)\) observed with error at time \(t\_i\). Then, our log-abundance observation model equation becomes \(Y(t\_i) = X(t\_i) + F\_i\), where \(F\_i\) follows a normal distribution with mean \(0\) and variance \(\tau^2\) (\(F\_i \sim \text{Normal}(0,\tau^2)\)). This state-space model has four unknown parameters: \(\ln \lambda = a\) (the trend parameter or population growth rate under density independence; note the notation differs from Humbert *et al.* (2009)), \(\sigma^2\) (variability of process noise), \(\tau^2\) (variability of observer noise), and \(x\_0\) (the initial log-abundance population).

The EGSS model has a multivariate normal log-likelihood function given
by (***EqA17*** in Humbert *et al.*, (2009)):

\[
\ln{L}(x\_0, a, \sigma^2, \tau^2) = -\frac{(q+1)}{2}\ln(2\pi)-\frac{1}{2}\ln(|\mathbf{V}|)-\frac{1}{2}(\mathbf{y}-\mathbf{m})'\mathbf{V}^{-1}(\mathbf{y}-\mathbf{m})
\]

For the numerical optimization, we will need three arguments:

1. A vector of time-series log-observed abundances `yt` (\(\mathbf{y}\))
2. A vector of observation times `tt`
3. A vector of initial parameters `fguess` (a first guess for the
   **four** parameters in EGSS), that could be roughly computing by
   provide the vector of log-abundance observations `yt` (\(\mathbf{y}\))
   and the vector of observation times `tt` with function
   `guess_eggs()`

```
guess_egss <- function(yt,tt){

  # Time-vector starting in 0.
  t.i       <- tt-tt[1];
  # Number of time-series transitions
  q         <- length(yt)-1;
  # length of time-series
  qp1       <- q+1;
  # time intervals (named as S.t in H_O and sometimes in DP_E)
  t.s       <- t.i[2:qp1]-t.i[1:q];

  #The Exponential Growth Observation Error (EGOE in H_O) initial values
  # mean of the observations as assumed to arise from stationary distribution
  Ybar      <- mean(yt);
  # mean of the time series
  Tbar      <- mean(t.i)
  # trend parameter for EGOE (a = ln(lambda) in H_O)
  a.egoe    <- sum((t.i-Tbar)*(yt-Ybar))/sum((t.i-Tbar)*(t.i-Tbar));
  # Initial population  of EGOE
  x0.egoe   <- Ybar-a.egoe*Tbar
  # sigma square for EGOE is 0 (assume no ecological process variation)
  ssq.egoe  <- 0
  # estimate of initial population observed under EGOE
  Yhat.egoe <- x0.egoe+a.egoe*t.i;
  # initial value for tau^2
  tsq.egoe  <- sum((yt-Yhat.egoe)*(yt-Yhat.egoe))/(q-1);

  #The Exponential Growth Process Noise (EGPN in H_O) initial values
  # Square root of time intervals (time trend?)
  Ttr       <- sqrt(t.s);
  # Observed trend?
  Ytr       <- (yt[2:qp1]- yt[1:q])/Ttr;
  # trend parameter for EGPN (mu = ln(lambda) in H_O)
  a.egpn    <- sum(Ttr*Ytr)/sum(Ttr*Ttr);
  # Trend of observed estimated
  Ytrhat    <- a.egpn*Ttr;
  # initial value for sigma^2
  ssq.egpn  <- sum((Ytr-Ytrhat)*(Ytr-Ytrhat))/(q-1);
  # tau square for EGPN is 0 (assume no observation variation)
  tsq.egpn  <- 0;
  # Initial population  of EGPN is the first observation
  x0.egpn   <- yt[1];

  #four parameters needed in EGSS and OUSS.NoSt
  a0    <- (a.egoe+a.egpn)/2;
  ssq0      <- ssq.egpn/2;
  tsq0      <- tsq.egoe/2;
  x0.out    <- (x0.egoe+x0.egpn)/2;

  return(c(a0, ssq0, tsq0, x0.out))
}
```

The numerical optimization for computing the Restricted Maximum Likelihood
Estimates for the multivariate normal distribution of parameters in `R` is:

```
negloglike_egss_reml <- function(fguess,yt,tt){

  sigmasq <- exp(fguess[1]); #in egss_reml, this have only the two parameters
  tausq   <- exp(fguess[2]);
  q       <- length(yt) - 1;
  qp1     <- q+1;

  ss      <- tt[2:qp1]-tt[1:q];
  wt      <- (yt[2:qp1]-yt[1:q])/ss;
  ut      <- wt[2:q]-wt[1:q-1];
  vx      <- matrix(0,qp1,qp1);
  for(i in 1:q){
    vx[(i+1):qp1,(i+1):qp1] <- matrix(1,(qp1-i),(qp1-i))*tt[i+1];
  }
  Sigma.mat<- sigmasq*vx;
  Itausq  <- matrix(rep(0,(qp1*qp1)), nrow=qp1, ncol=qp1);
  diag(Itausq)<- rep(tausq,qp1);
  V       <- Sigma.mat + Itausq;

  D1mat   <- cbind(-diag(1/ss),matrix(0,q,1))+cbind(matrix(0,q,1),diag(1/ss));
  D2mat   <- cbind(-diag(1,(q-1)),matrix(0,(q-1),1)) + cbind(matrix(0,(q-1),1),diag(1,(q-1)));
  V2      <- D2mat%*%D1mat%*%V%*%t(D1mat)%*%t(D2mat);

  ofn=((q-1)/2)*log(2*pi)+(0.5*log(det(V2))) + (0.5*(ut%*%ginv(V2)%*%ut));

  return(ofn)
}
```

To compute the EGSS-REMLs we use the function `egss_reml()`

```
egss_reml <- function(yt,tt,fguess){

  #Temporal vectors
  t.i     <- tt-tt[1];
  q       <- length(t.i)-1;
  qp1     <- q+1;
  t.s     <- t.i[2:qp1] - t.i[1:q];

  # initial guesses (sigmasq and tausq at log scale)
  guess.optim <- c(log(fguess[2:3]))
  # numerical optimization
  optim.out   <- optim(par=guess.optim,
                       fn=negloglike_egss_reml,
                       method="Nelder-Mead",
                       yt=yt,
                       tt=t.i)

  #extract parameters estimated by REML
  sigmasq <- exp(optim.out$par)[1]
  tausq <- exp(optim.out$par)[1]

  #to estimate trend parameter (a) and initial population (x0)
  vx      <- matrix(0,qp1,qp1);
  for(i in 1:q){
    vx[((i+1):qp1),((i+1):qp1)] <- matrix(1,(qp1-i),(qp1-i))*t.i[(i+1)];
  }
  Sigma.mat     <- sigmasq*vx;
  Itausq        <- matrix(rep(0,(qp1*qp1)),
                          nrow=qp1,
                          ncol=qp1);
  diag(Itausq)  <- rep(tausq,qp1);
  V             <- Sigma.mat + Itausq;
  D1mat=cbind(-diag(1/t.s),
              matrix(0,q,1))+cbind(matrix(0,q,1),
                                   diag(1/t.s));
  V1mat=D1mat%*%V%*%t(D1mat);
  W.t=(yt[2:qp1]-yt[1:q])/t.s;
  j1=matrix(1,q,1);
  V1inv=ginv(V1mat);

  #Trend parameter
  a.reml=(t(j1)%*%V1inv%*%W.t)/(t(j1)%*%V1inv%*%j1);

  j=matrix(1,qp1,1);
  Vinv=ginv(V);

  #initial population
  x0.reml=(t(j)%*%Vinv%*%(yt-as.numeric(a.reml)*t.i))/(t(j)%*%Vinv%*%j);

  #Extract REMLs and AIC
  remls      <- c(a.reml,exp(optim.out$par[1:2]),x0.reml)
  lnL.hat     <- - optim.out$value[1]
  AIC         <- -2*lnL.hat + 2*2 #where 2 = length(REMLs)...

  out         <- list(remls=remls,
                      lnL.hat = lnL.hat,
                      AIC=AIC)
  return(out)
}
```

With the EGSS-REML values, we can predict the trajectory of the latent ecological process with the function `egss_predict()`

```
egss_predict <- function(yt,tt,parms,plot.it="TRUE"){

  # Time-vector starting in 0.
  t.i     <- tt-tt[1];
  q       <- length(t.i)-1;
  qp1     <- q+1;
  t.s     <- t.i[2:qp1] - t.i[1:q];

  # parameters ()
  a.reml  <- parms[1];
  sigmasq  <- parms[2];
  tausq    <- parms[3];
  x0.reml <- parms[4];

  #Calculate estimated population size for EGSS model

  m=rep(1,qp1); # Will contain Kalman means for Kalman calculations.
  v=rep(1,qp1); # Will contain variances for Kalman calculations.

  m[1]=x0.reml; # Initial mean of Y(t).
  v[1]=tausq; # Initial variance of Y(t).

  for (ti in 1:q) # Loop to generate estimated population abundances
  { # using Kalman filter (see equations 6 & 7, # Dennis et al. (2006)).
    m[ti+1]=a.reml+(m[ti]+((v[ti]-tausq)/v[ti])*(yt[ti]-m[ti]));
    v[ti+1]=tausq*((v[ti]-tausq)/v[ti])+sigmasq+tausq;
  }

  # The following statement calculates exp{E[X(t) | Y(t), Y(t-1),...,Y(0)]};
  # see equation 54 in Dennis et al. (2006).

  Predict.EGSS.REML = exp(m+((v-tausq)/v)*(yt-m));

  if(plot.it=="TRUE"){
    #  Plot the data & model-fitted values
    #X11()
    plot(tt,exp(yt),xlab="Time",ylab="Population abundance",
         type="b",cex=1.5, lwd = 1.5, lty = 1,
         main="Predicted (--) and observed (-o-) abundances");
            # Population data are circles.
    points(tt,Predict.EGSS.REML, type="l", lwd=1, lty = 2);
  }

  return(list(cbind(Time = tt, Predict.EGSS.REML, Observed.y = exp(yt))))
}
```

And also simulate trajectories with the function `egss_sim()`

```
egss_sim <- function(nsims,tt,parms){

  # time and temporal scale
  t.i    <- tt-tt[1];
  q      <- length(t.i)-1;
  qp1    <- q+1;

  # parameters
  a      <- parms[1];
  sigmasq<- parms[2];
  tausq  <- parms[3];
  x0     <- parms[4];

  vx     <- matrix(0,qp1,qp1);
  for(i in 1:q){
    vx[((i+1):qp1),((i+1):qp1)] <- matrix(1,(qp1-i),(qp1-i))*t.i[(i+1)];
  }

  Sigma.mat<- sigmasq*vx;
  Itausq<- matrix(rep(0,(qp1*qp1)),
                          nrow=qp1,
                          ncol=qp1);
  diag(Itausq)  <- rep(tausq,qp1);
  V     <- Sigma.mat + Itausq;
  a.vec     <- matrix((x0+a*t.i),
                          nrow=qp1,
                          ncol=1);
  out   <- randmvn(n=nsims,
                   mu.vec=a.vec,
                   cov.mat=V);

  return(out)
}
```

And we can conduct a parametric bootstrap to estimate confidence intervals of the predicted trajectory

```
egss_pboot <- function(B=2,parms,yt, tt){
    
    t.i <- tt-tt[1]
    nparms    <- length(parms);
    preds.boot1<- matrix(0,nrow=B,ncol=length(t.i))

        boot.remles <- matrix(0,nrow=B,ncol=nparms+1); 
        all.sims  <- egss_sim(nsims=B,parms=parms,tt=tt);
        all.preds <- egss_predict(parms=parms, yt=yt,tt=tt,plot.it="FALSE")
        reml.preds<- all.preds[[1]][,2]

        for(b in 1:B ){
        
            bth.timeseries <- all.sims[,b];
            remles.out <- egss_reml(yt=bth.timeseries, tt=tt, fguess=parms);
            boot.remles[b,] <- c(remles.out$remls, remles.out$lnL.hat);
            all.bootpreds <- egss_predict(parms=remles.out$remls, yt=bth.timeseries,tt=tt,plot.it="FALSE");
            preds.boot1[b,] <- all.bootpreds[[1]][,2]
        } 
    
        CIs.mat <- apply(boot.remles,2,FUN=function(x){quantile(x,probs=c(0.025,0.975))});
        CIs.mat <- rbind(CIs.mat[1,1:4],parms,CIs.mat[2,1:4]);
        rownames(CIs.mat) <- c("2.5%","REMLE","97.5%");
        colnames(CIs.mat) <- c("a", "sigmasq","tausq", "x0");
        
        preds.CIs1 <- apply(preds.boot1,2,FUN=function(x){quantile(x,probs=c(0.025,0.975))});
        mean.boots <- apply(preds.boot1,2,FUN=function(x){quantile(x,probs=0.50)})
        preds.CIs1 <- t(rbind(tt,reml.preds-(mean.boots-preds.CIs1[1,]), reml.preds, reml.preds+(preds.CIs1[2,]-mean.boots)));
        colnames(preds.CIs1) <- c("Time","CI.lower","REMLE","CI.upper");

        boot.list <- list(boot.remles = boot.remles, CIs.mat = CIs.mat, preds.CIs1 = preds.CIs1);
        
        return(boot.list)
        
}
```

##### 2.3.3 Functions for the Ornstein-Uhlenbeck diffusion GSS - OUSS model

The key to generalize the GSS for unequal sampling intervals lies in the mathematical insight that the solution of the discrete-time GSS model matches the solution at discrete time points of a diffusion process, which is a continuous time stochastic process. Specifically, the solution of the discrete time Gompertz model in the log scale matches exactly the well-known Ornstein-Uhlenbeck (OU) Gaussian diffusion process (Dennis & Ponciano, 2014). Let \(N\_t\) represent the population abundance in the Gompertz diffusion with environmental stochasticity and no demographic stochasticity (Dennis & Ponciano, 2014; Ponciano, 2018). This diffusion process is a well-known continuous-time version of an autoregressive process of order 1, characterized by a joint multivariate normal distribution of values across time points. The process is defined by its infinitesimal mean, variance, and covariance parameters (Dennis & Ponciano, 2014; Ponciano, 2018).

The infinitesimal mean and variance of the process are given by:
\(m\_N (n)= \theta n[\ln \kappa - \ln ⁡n]\) and \(\sigma\_N^2 (n)= \beta^2 n^2\), respectively; here \(\theta\) represents the speed of equilibration, \(\kappa\) the equilibrium abundance, and \(\beta\) scales the random perturbation by environment stochasticity, \(dW\). The OU diffusion process is usually presented in its stochastic differential equation form \(dN\_t = \theta N\_t [\ln\kappa - ⁡\ln{N\_t})dt + \beta N\_t dW\_t\). A smooth transformation to \(N\_t\), given by \(X\_t=g(N\_t)\) (e.g. \(X\_t=\ln ⁡N\_t\)), is also a diffusion process whose infinitesimal mean is given by \(m\_X (x)=m\_N (n) g'(n) + 1/2 \sigma\_N^2 (n)g''(n)\) and infinitesimal variance by \(\sigma\_X^2 (x) = \sigma\_N^2 n [g' (n)]^2\), where \(n=g^{-1} (x)\). This result is the well-known Itô-transformation for diffusion processes widely used in stochastic population dynamics modeling. For \(g(n) = \ln⁡n\), the infinitesimal mean simplifies to \(m\_X (x)= \theta (\mu - x)\), where \(\mu = \ln\kappa - \frac{\beta^2}{2 \theta}\), and the infinitesimal variance becomes \(\sigma\_X^2 (x)= \beta^2\). Thus, the stochastic differential equation of the logarithmic process \(X\_t\) is \(dX\_t= \theta(\mu - X\_t)dt + \beta dW\_t\). If the process starts at an initial log-abundance \(X\_0=x\_0\), the expected value and variance at any time \(t\) are given by \(\text{E}[X\_t | X\_0 = x\_0] = \mu - (\mu - x\_0) e^{\theta - t}\) and \(\text{V}[X\_t|X\_0 = x\_0] = \frac{\beta^2}{2\theta} (1-e^{-2\theta})\) respectively (Dennis & Ponciano, 2014). Over time, the process converges to a stationary distribution with mean \(\mu\) and variance \(\frac{\beta^2}{2\theta}\).

The one-to-one relationships of the discrete equal sampling GSS model parameters to the continuous OUSS model parameters are (Dennis & Ponciano, 2014):

\[
\begin{array}{ccc}
a &=& \mu(1- e^{-\theta}) \\
c &=& e^{-\theta} \\
\sigma^2 &=& \frac{(1-e^{-2\theta})\beta^2}{2\theta} \\
\tau^2 &=& \tau^2 \\
\end{array}
\]

Thus, the OUSS model has four unknown parameters under stationary distribution: \(\mu\) (mean stationary log-abundance), \(\theta\_{density~dependent}\) (the trend parameter under density dependence, or rate to approach stationarity), \(\beta^2\) (variability of the process noise), and \(\tau^2\) (variability of sampling). The OUSS model also adds a component in sampling times \(t\_i\) not equally spaced \(Y(t\_i) = X(t\_i) + F\_i\), where the observation error keeps a normal distribution and the same unknown parameter (\(F\_i \sim \text{Normal}(0,\tau^2)\)), and the underlying unobserved population \(X(t\_i)\) follows a continuous-time version of the GSS model. With the strength of density dependence parameter (\(c\)) ranging between \(0\) and \(1\), the dynamic of the population is stationary.

The inverse relationship between the OUSS and the GSS model parameters are:

\[
\begin{array}{ccc}
\mu &=& \frac{a}{1-c} \\
\theta &=& -\ln{c} \\
\beta^2 &=& -\frac{2\sigma^2 \ln{c}}{1-c^2} \\
\tau^2 &=& \tau^2 \\
\end{array}
\]

The normal stationary probability distribution has mean \(\mu\) and variance \(\frac{\beta^2}{2\theta}\) (Dennis & Ponciano, 2014). If the initial log-abundance of the population does not meet this assumption (e.g., it is under transition growth), a nonstationary distribution could be modeled with a different maximum likelihood estimation approach (Dennis & Ponciano, 2014). The normal transition in nonstationary cases has a mean \(\mu - (\mu-x\_0) e^{-\theta t}\) and variance (\(\frac{\beta^2}{2\theta} (1-e^{-2\theta})\)), adding an extra parameter to estimate (\(x\_0\)). Given that a restricted maximum likelihood estimation is not available for nonstationary OUSS (Dennis & Ponciano, 2014), we modeled the dynamic of the populations for density-independent (including initial nonstationary distributions) as EGSS, while stationary distributions as OUSS.

The multivariate normal log-likelihood for the stationary OUSS model is given by (see ***Eq. 19*** Dennis & Ponciano, (2014)).

\[
\ln{L}(\mu, \theta, \beta^2, \tau^2) = -\frac{(q+1)}{2}\ln(2\pi)-\frac{1}{2}\ln(|\mathbf{V}|)-\frac{1}{2}(\mathbf{y}-\mathbf{m})'\mathbf{V}^{-1}(\mathbf{y}-\mathbf{m})
\]

where \(q\) is the number of time-series transitions (thus, \(q+1\) reflect
the length of the time-series, with the initial population estimation
\(y\_0\) as a realized value of the random variable \(Y(0)\)), \(\mathbf{V}\)
is the variance-covariance matrix (with diagonal computed from
\(\text{V}[Y(t\_i)] = \tau^2+\frac{\beta^2}{2\theta}\); ***Eq. 17*** in Dennis & Ponciano, (2014)),
\(\mathbf{y}\) is the data values (\(y\_0\), \(y\_1\), \(y\_2\), …, \(y\_q\)), and
\(\mathbf{m}\) is the vector of same \(\mu\) in all \(q+1\) times
(\(E[Y(t\_i)] = \mu]\); ***Eq. 16*** in Dennis & Ponciano, (2014)).

This function requires three arguments for the numerical optimization in
`R`:

1. A vector of time-series of log-observed abundances `yt`
   (\(\mathbf{y}\))
2. A vector of observation times `tt`
3. A vector of parameters `fguess` (a first guess for the **four**
   parameters), that could be roughly computing by provide the vector
   of log-abundance observations `yt` (\(\mathbf{y}\)) and the vector of
   observation times `tt` with the function `guess_ouss()`

```
guess_ouss <- function(yt,tt){

  # Time-vector starting in 0.
  t.i     <- tt-tt[1];
  # Number of time-series transitions
  q       <- length(yt)-1;
  # length of time-series
  qp1     <- q+1;
  # time intervals
  t.s     <- t.i[2:qp1]-t.i[1:q];
  # mean of the observations as assumed to arise from stationary distribution
  Ybar    <- mean(yt);
  # Variance of the observations
  Yvar    <- sum((yt-Ybar)*(yt-Ybar))/q;
  # Initial mu estimate (at stationary distribution)
  mu1     <- Ybar;

  # Kludge an initial value for theta based on mean of Y(t+s) given Y(t).
  th1     <- -mean(log(abs((yt[2:qp1]-mu1)/(yt[1:q]-mu1)))/t.s);
  # Moment estimate using stationary distribution
  bsq1    <- 2*th1*Yvar/(1+2*th1);
  # Observation error variance, assumed as first guess as betasq=tausq.
  tsq1    <- bsq1;

  # What to do if initial guesses is three 0's (or NAs)? Assume arbitrary values
  three0s <- sum(c(th1,bsq1,tsq1))

  if(three0s==0|is.na(three0s)){
    th1   <- 0.5;
    bsq1  <- 0.09;
    tsq1  <- 0.23;}

  out1    <- c(th1,bsq1,tsq1);

  # What to do if initial guesses are too little? Assume arbitrary values
  if(sum(out1<1e-7)>=1){
    out1  <- c(0.5,0.09,0.23)}

  out     <- c(mu1,out1);

  return(abs(out))
}
```

The numerical optimization for computing the parameters Restricted Maximum Likelihood
Estimate within the multivariate log-likelihood for the
stationary Ornstein-Uhlenbeck State-Space (OUSS) in `R` is:

```
negloglike_ouss_reml=function(yt,tt,fguess){
  # Constrains parameters theta, beta^2, and tau^2 > 0

  # speed of equilibration (Eq1 in DP_E)
  theta  <- exp(fguess[2]);
  # variability of process noise
  betasq <- exp(fguess[3]);
  # variability of sampling
  tausq  <- exp(fguess[4]);
  # number of time-series transitions
  q      <- length(yt) - 1;
  # length of time-series
  qp1    <- q+1;
  # Variance (Eq11 in DP_E)
  Var.inf<- betasq/(2*theta);
  # time intervals (not used here?)
  t.s    <- tt[2:qp1] - tt[1:q];
  # part of Eq18 in DP_E
  t.cols <- matrix(rep(tt,each=qp1),
                          nrow=qp1,
                          ncol=qp1,
                          byrow=FALSE);
  # (part of Eq18 in DP_E)
  t.rows <- t(t.cols);
  # (part of Eq18 in DP_E)
  abs.diffs     <- abs(t.rows-t.cols);

  # Covariance of the process (Eq18 in DP_E)
  Sigma.mat     <- Var.inf*exp(-theta*abs.diffs);
  # Create a matrix full of 0s of the length of time series
  Itausq <- matrix(0,qp1,qp1);
  # Repeat the observation error variance guess in the diagonal of the matrix
  diag(Itausq)  <- rep(tausq,qp1);
  # add Covariance with the matrix
  V      <- Sigma.mat+Itausq;
  # Create the differencing matrix **D**
  Dmat   <- cbind(-diag(1,q),matrix(0,q,1)) + cbind(matrix(0,q,1),diag(1,q));
  # Variance-covariance matrix **Phi** (Eq20 DP_E)
  Phi.mat<- Dmat%*%V%*%t(Dmat);
  # simple differencing of the observations (W_i? )
  wt     <- yt[2:qp1]-yt[1:q];

  # note the signs change because we want here the negative log-likelihood (Eq22*-1)
  neglogl<- (q/2)*log(2*pi) + (1/2)*log(det(Phi.mat)) + (1/2)*wt%*%ginv(Phi.mat)%*%wt;

  # What to do if the `neglogl` is not finite? assign a big number of 50000
  if(is.infinite(neglogl)==TRUE){
    return(50000)}else{
      return(neglogl)}
}
```

To compute the OUSS-REMLs we implement the function `ouss_reml()`

```
ouss_reml <- function(yt, tt, fguess){

  # Time-vector starting in 0.
  t.i           <- tt-tt[1];
  # Number of time-series transitions
  # length of time-series
  q             <- length(yt)-1;
  qp1           <- q+1;
  # time intervals
  t.s           <- t.i[2:qp1]-t.i[1:q];
  # initial guesses (all, but negloglike.OU.reml will use only fguess[2:4])
  guess.optim   <- c(fguess[1],
                     log(fguess[2:4]));
  # numerical optimization
  optim.out     <- optim(par = guess.optim,
                         fn=negloglike_ouss_reml,
                         method="Nelder-Mead",
                         yt=yt,
                         tt=t.i);
  # Restricted maximum likelihood estimates (REMLs) and lnL.hat
  remls        <- exp(optim.out$par);
  theta.reml   <- remls[2];
  betasq.reml  <- remls[3];
  tausq.reml   <- remls[4];

  lnL.hat       <- -optim.out$value[1];

  # Variance (Eq11 in DP_E)
  Var.inf       <- betasq.reml/(2*theta.reml)
  # creates an matrix full of 1 dim qp1 x qp1
  vx            <- matrix(1,qp1,qp1);
  # iterate to fill the matrix (couldn't find vx in DP_E!)
  for (t.i in 1:q){
    vx[(t.i+1):qp1,t.i]=exp(-theta.reml*cumsum(t.s[t.i:q]));
    vx[t.i,(t.i+1):qp1]=vx[(t.i+1):qp1,t.i];
  }
  # ?
  Sigma.mat     <- vx*Var.inf;
  # Create a matrix full of 0s of the length of time series
  Itausq        <- matrix(0,qp1,qp1);
  # Repeat the observation error variance reml in the diagonal of the matrix
  diag(Itausq)  <- rep(tausq.reml,qp1);
  # Variance-covariance matrix (V.hat) evaluated with remls to estimate mu.hat
  V.reml       <- Sigma.mat+Itausq;
  # column vector matrix of ones
  j             <- matrix(1,qp1,1);
  # Inverse matrix (part of Eq23 in DP_E)
  Vinv          <- ginv(V.reml);
  # REML of mu (mu.hat) with Eq23 in DP_E
  mu.reml      <- (t(j)%*%Vinv%*%yt)/(t(j)%*%Vinv%*%j);
  #AIC
  AIC           <- -2*lnL.hat + 2*4 #where 4 = length(mles)...

  #Results
  out           <- list(remls = c(mu.reml,
                                   theta.reml,
                                   betasq.reml,
                                   tausq.reml),
                        lnLhat = lnL.hat,
                        AIC = AIC)
  return(out)
}
```

With the OUSS-REML values, we can predict the trajectory with the
function `ouss_predict()`

```
ouss_predict <- function(yt,tt,parms, plot.it="TRUE"){

  t.i             <- tt-tt[1];
  q               <- length(t.i)-1;
  qp1             <- q+1;

  # parameters
  mu              <- parms[1];
  theta           <- parms[2];
  betasq          <- parms[3];
  tausq           <- parms[4];

  Var.inf         <- betasq/(2*theta);
  t.s             <- t.i[2:qp1] - t.i[1:q];
  t.cols          <- matrix(rep(t.i,each=qp1),nrow=qp1,ncol=qp1, byrow=FALSE);
  t.rows          <- t(t.cols);
  abs.diffs       <- abs(t.rows-t.cols);

  nmiss           <- t.s-1;
  long.nmiss      <- c(0,nmiss);
  Nmiss           <- sum(nmiss)

  long.t          <- t.i[1]:max(t.i)
  where.miss      <- which(is.na(match(x=long.t,table=t.i)),
                           arr.ind=TRUE)
  lt.cols         <- matrix(rep(long.t),
                            nrow=(qp1+Nmiss),
                            ncol=(qp1+Nmiss),
                            byrow=FALSE);
  lt.rows         <- t(lt.cols);
  labs.diffs      <- abs(lt.rows-lt.cols);

  Sigma.mat       <- Var.inf*exp(-theta*abs.diffs);
  Itausq          <- matrix(0,qp1,qp1);
  diag(Itausq)    <- rep(tausq,qp1);
  V               <- Sigma.mat+Itausq;

  long.V          <- Var.inf*exp(-theta*labs.diffs) + diag(rep(tausq,(qp1+Nmiss)))

  Predict.t       <- rep(0,qp1);
  Muvec           <- rep(mu,q);
  miss.predict    <- list()
  Muvec.miss      <- rep(mu,qp1);
  start.miss      <- 1
  stop.miss       <- 0
  for (tj in 1:qp1){
    Y.omitj       <- yt[-tj];    #  Omit observation at time tj.
    V.omitj       <- V[-tj,-tj];  #  Omit row tj and col tj from var-cov matrix.
    V12           <- V[tj,-tj];       #  Submatrix:  row tj without col tj.
    Predict.t[tj] <- mu+V12%*%ginv(V.omitj)%*%(Y.omitj-Muvec);  #  Graybill's 1976 Thm.

    if(long.nmiss[tj]==0){
      miss.predict[[tj]] <- Predict.t[tj]}else
        if(long.nmiss[tj]>0){

          start.miss <- stop.miss+1
          ntjmiss    <- long.nmiss[tj]
          mu.miss    <- rep(mu,ntjmiss);
          ind.tjmiss <- where.miss[start.miss:(start.miss+(ntjmiss-1))]
          stop.miss  <- stop.miss+ntjmiss

          longV12    <- long.V[ind.tjmiss,-where.miss]

          miss.predict[[tj]] <- c(mu.miss + longV12%*%ginv(V)%*%(yt-Muvec.miss),
                                  Predict.t[tj])
        }
  }

  Predict.t <- exp(Predict.t);
  LPredict.t <- exp(as.vector(unlist(miss.predict)))

  isinf <- sum(is.infinite(Predict.t))
  if(isinf>0){
    where.infs <- which(is.infinite(Predict.t)==TRUE, arr.ind=TRUE)
    Predict.t[where.infs] <- .Machine$double.xmax
  }

  isinf2 <- sum(is.infinite(LPredict.t))
  if(isinf2>0){
    where.infs <- which(is.infinite(LPredict.t)==TRUE, arr.ind=TRUE)
    LPredict.t[where.infs] <- .Machine$double.xmax
  }

  if(plot.it=="TRUE"){
    #  Plot the data & model-fitted values
    #X11()
    plot(tt,exp(yt),xlab="Time",ylab="Population abundance",type="b",cex=1.5,
         main="Predicted (--) and observed (-o-) abundances");
        # Population data are circles.
    par(lty="dashed"); #  Predicted abundances are dashed line.
    points(tt,Predict.t, type="l", lwd=1);
  }

  return(list(cbind(tt,Predict.t,exp(yt)), cbind(long.t,LPredict.t) ))
}
```

And also simulate trajectories with `ouss_sim()`

```
ouss_sim <- function(nsims,tt,parms){

  # Time-vector starting in 0.
  t.i       <- tt-tt[1];
  # Number of time-series transitions
  q         <- length(t.i)-1;
  # length of time-series
  qp1       <- q+1;

  # parameters
  mu        <- parms[1];
  theta     <- parms[2];
  betasq    <- parms[3];
  tausq     <- parms[4];

  Var.inf   <- betasq/(2*theta);
  t.s       <- t.i[2:qp1] - t.i[1:q];
  t.cols    <- matrix(rep(t.i,each=qp1),
                      nrow=qp1,
                      ncol=qp1,
                      byrow=FALSE);
  t.rows    <- t(t.cols);
  abs.diffs <- abs(t.rows-t.cols);
  V         <- Var.inf*exp(-theta*abs.diffs);
  diag(V)   <- diag(V) + rep(tausq,qp1);
  m.vec     <- rep(mu,qp1);
  out       <- randmvn(n=nsims,
                       mu.vec=m.vec,
                       cov.mat = V)
  return(out)
}
```

### 3 eBird data organization

Users should download the `ebd` file from
eBird. See the supplement to
Johnston *et al.* (2021) in
Strimas-Mackey *et al.* (2023), which is a key reference for the next section.

#### 3.1 Download eBird data

##### 3.1.1 Go to eBird and sign in

Go to eBird. You have to sign-in into
eBird:

##### 3.1.2 Request data

If you are in the home page, check you are signed-in and move down on the
page to “Request data”.

You have to submit an application to have access to the data. Once you
have access, click on “Basic dataset (EBD)” (it will show the window of access).

Then, you can select by species, region, and/or date. In our case, lets
download *Rostrhamus sociabilis* in Florida (US).

In the options, include the sampling event data always is a good recommendation. In snail kite for US it will save the sampling event, but for other Neotropical species tested in preliminary attempts, the sampling data was not included and the user should download the “Sampling event data” of 5.5 GB (comprised in `.tar` format). This step is highly recommended if you are interested in control for imperfect detection!!

After submitting the request, the link to download will arrive to the
email registered in your eBird account. You can save the `.txt` files
in a `data_raw` directory to be called during the refining process through filtering.

This file will have the detection and observer counts for our species.

#### 3.2 Pre-filtering

We can simplify the eBird data selecting only columns of our interest
(it will reduce the size of the dataset)

```
colsE <- c("observer_id", "sampling_event_identifier",
           "group identifier",
           "common_name", "scientific_name",
           "observation_count",
           "country", "state_code", "locality_id", "latitude", "longitude",
           "protocol_type", "all_species_reported",
           "observation_date",
           "time_observations_started",
           "duration_minutes", "effort_distance_km",
           "number_observers")
```

To conduct some filters, we will generate temporal files in our
computer. Here we generate only a single temporal file that can be
overwritten to assess different species.

```
f_ebd <- "data_tmp/ebd_Examples.txt" 
f_sed <- "data_tmp/sed_Examples.txt"
```

The construction of time-series of the individuals counted from
eBird will assume spatiotemporal subsampling, selecting a single value
with the high counts (assuming to be the minimum number of individuals detected) per week in spatial sampling units of ~ \(100 \text{ km}^2\). To construct a discrete global grid system, we can use the function `dgconstruct()` in the package `dgconstruct`; the argument `spacing` indicates the spacing between the center of adjacent cells (related with Characteristic Length Scale - CLS), in our case `spacing = 11` indicates a diameter of ~\(11\ km\), representing an area of \(95.98\ km^2\) (~ \(100\ km^2\)).

```
#specify seed for random number generation
dggs_pop <- dgconstruct(spacing = 11)
```

```
## Resolution: 12, Area (km^2): 95.9778454662114, Spacing (km): 9.67579208698331, CLS (km): 11.0545373458563
```

#### 3.3 Refinament data by filtering

The package `auk`, in combination with `tidyverse`, allows the filtering
of the eBird data (see Strimas-Mackey *et al.* 2023). Note that the
first function `auk_ebd()` includes the path of the eBird data downloaded and saved in your working directory (up to November 2024), and it is the initial creation of an `auk_ebd` object. Then, different functions serve to filter the data by the metadata of the checklists. We followed the next order below:

- by protocol (`auk_protocol()`; only traveling or stationary),
- by distance (`auk_distance()`; \(≤5\) km),
- by duration (`auk_duration()`; \(≤5\) hours, note the units are in minutes: \(300\)), and
- only complete lists (`auk_complete()`).

Then, the filters defined are converted to an AWK script with the function `auk_filter()`, generating a filtered eBird Reference Dataset (ERD), storing in the temporal files `f_ebd` with only the selected columns (defined in the pre-filtering). Finally, the function `read_ebd()` read the filtered file.

```
ebd_filt <- auk_ebd("data_raw/ebd_US-FL_snakit_smp_relNov-2024.txt") %>%
  auk_protocol(c("Traveling", "Stationary")) %>%
  auk_distance(distance = c(0,5)) %>%
  auk_duration(duration = c(0,300))%>%
  auk_complete() %>%
  auk_filter(f_ebd,overwrite=T, keep = colsE) %>%
  read_ebd()
```

Then, just for the sake of double checking and organization, we can remove
the observations without counts, add distance \(0\) to
stationary protocols, modify the time of observations started to decimal,
round hour sampling to an integer, extract year, month, week, and
day\_of\_year. Also, we can confirm and filter out by effort, such as
observers \(≤10\), distance \(≤5 \text{ km}\), duration \(≤5 \text{ hours}\),
and only records with counts included. These covariates could be further used to test their effects on population dynamic estimates (see Fink *et al.* 2023).

```
#Some effort extraction and confirmation
ebd_filt <- ebd_filt %>%
  mutate(
    # We don't want here count in 'X', to convert to NA we use `as.integer()`
    observation_count = as.integer(observation_count),
    # effort_distance_km to 0 for non-travelling counts
    effort_distance_km = if_else(protocol_type == "Stationary",
                                 0, effort_distance_km),
    # convert time to decimal hours since midnight
    time_observations_started = time_to_decimal(time_observations_started),
    hour_sampling = round(time_observations_started, 0),
    # split date into year, month, week, and day of year
    year = year(observation_date),
    month = month(observation_date),
    week = week(observation_date),
    day_of_year = yday(observation_date)) %>%
  filter(number_observers <= 10,         #Only list with less than 10 observers
         effort_distance_km <= 5,        #be sure of distance effort
         duration_minutes %in% (0:300),  #be sure of duration effort
         !is.na(observation_count))      #only records with counts reported
```

Now we can add a new variable that identify each `cell` from a grid of
hexagons (spatial sampling units), using the `longitude` and `latitude`
information of our `ebd_filt` dataset and the function
`dgGEO_to_SEQNUM()`. With the new variable, we can extract the maximum
count of individuals and number of checklists per week per cell.

```
SnailKite <- ebd_filt %>%
  mutate(cell = dgGEO_to_SEQNUM(dggs_pop, #id for cells
                                longitude, latitude)$seqnum) %>%
  group_by(cell, year, month, week) %>%
  mutate(max_count = max(observation_count, na.rm = T), 
         n_lists = n()) |>
  ungroup()
```

This file is saved as a backup

```
#and save the filter
saveRDS(SnailKite, "data_tmp/SnailKiteCellsID_filtered.rds")
```

#### 3.4 Time series of eBird weekly high-counts

We adjusted the time-series from the 1st week of 2018 (January) to the
last with data of 2024 (November). Since 2018, snail kites reached more than 1000 annual records.

```
SnailKite <- readRDS("data_tmp/SnailKiteCellsID_filtered.rds")

png('data_tmp/FigSI-1_HistogramTemporalBias.png',
    width = 10, height = 5, units = "in", res = 300) 

hist(SnailKite$year, 
     breaks = 50, 
     main = "Florida Snail kites, checklists per year", 
     xlab = "Year")
abline(h = 1000, v = 2017, col = "red")
dev.off()
```

```
## quartz_off_screen 
##                 2
```

In addition, the spatial cell with higher records overlaps with the
Payne’s Prairie State Park wetland in Alachua County, north central
Florida, where snail kites established in 2018. We
extracted the high count per week in the cell of Payne’s Prairie to illustrate our method.

But first, we filter by observation data greater than or equal to
`2018-01-01`, and generate a new variable called `Time.t`, to have the
accumulated id of weeks from 2018 to the end of our time series. We used
the function `case_when()` based on `year`.

```
snailkites.week <- SnailKite |>
  filter(observation_date >= "2018-01-01") |>
  mutate(Time.t = case_when(year == 2018 ~ week,
                            year > 2018 ~ week+(52*(year-2018)))) 
summary(snailkites.week$observation_date)
```

```
##         Min.      1st Qu.       Median         Mean      3rd Qu.         Max. 
## "2018-01-01" "2020-01-21" "2021-12-28" "2021-09-06" "2023-03-13" "2024-11-30"
```

```
summary(snailkites.week$Time.t)
```

```
##    Min. 1st Qu.  Median    Mean 3rd Qu.    Max. 
##     1.0   107.0   208.0   192.1   271.0   360.0
```

Then, we group by the `cell` and `Time.t`, summarizing the maximum
integer count per week, named `Observed.y` in our data set.

```
snailkites.week.counts <- snailkites.week |>
  group_by(cell, Time.t) |>
  summarise(Observed.y = round(max(max_count),0))
```

```
## `summarise()` has grouped output by 'cell'. You can override using the
## `.groups` argument.
```

```
head(snailkites.week.counts)
```

And we can save this outcome as a backup.

```
saveRDS(snailkites.week, "data_tmp/SnailKiteCellsWeek.rds")
saveRDS(snailkites.week.counts, "data_tmp/SnailKiteCellsCountsWeek.rds")
```

This file will serve to generate a map figure with the sampling effort
after filtering the eBird data following best practices for analysis (see Johnston *et al.*, 2021).

### 4 Snail Kite in Payne’s Prairie from eBird

We focused on the sampling unit with higher number of checklists, which
correspond to the Payne’s Prairie State Park wetland system in Alachua
County. For this locality, we also can access counts published from Poli *et al.* (2020),
and 2 areas under current monitoring: Payne’s Prairie and Payne’s
Prairie Central.

This wetland overlaps with an hexagonal cell that concentrate most records.

#### 4.1 Map of sampling units

To generate the map of spatiotemporal sampling from eBird, we can use
the package `sf`, using the database included in the function
`st_as_sf()`.

```
#A global map to make figures ###
world1 <- sf::st_as_sf(maps::map(database = 'world', plot = FALSE, fill = TRUE))
world1
```

We can summarize the filtered ebd-data set `SnailKite` by the number of
observations in each hexagonal cell. Below code will generate a tibble
with two variables and 444 observations, the count of observations per
each cell.

```
#Get the number of observations in each cell
CellObservationsSK   <- SnailKite %>% 
  group_by(cell) %>%
  summarise(count=n())
```

And get a grid cell boundaries for cells with the observations counts
with the function `dgcellstogrid()`, using the discrete global grid
system saved as `dggs_pop`. **CAUTION**, if you generate `dggs_pop` in a
different computer or session than `SnailKite`, there could be conflict
with the `cell` id.

```
gridSnailKite <- dgcellstogrid(dggs_pop,CellObservationsSK$cell)
```

This is an `sf` object with the `cell` named as `seqnum`. We can update
the grid cells’ properties to include the number of lists in each cell
and handling the spatial data with `st_wrap_dateline()`

```
gridSnailKite <- merge(gridSnailKite, CellObservationsSK, by.x="seqnum", by.y="cell")

# Handle cells that cross 180 degrees
wrapped_gridSnailKite = st_wrap_dateline(gridSnailKite,
                                         options = c("WRAPDATELINE=YES",
                                                     "DATELINEOFFSET=180"), 
                                         quiet = TRUE)

#save the wrapped grid
saveRDS(wrapped_gridSnailKite, "data_tmp/wrapped_gridSnailKite.rds")
```

Some aesthetics are defined for the log-scales and arrows

```
my_breaks = c(7, 70, 700, 7000)

arrow1 <- tibble(
  x1 = -82.3,
  x2 = -80.75,
  y1 = 29.6,
  y2 = 29.7
)
```

and the figure is generated with `ggplot()`

```
Fig2a <- ggplot() +
  geom_sf(data = world1)+
  geom_sf(data=wrapped_gridSnailKite,
          aes(color = count, 
              fill = count),
          alpha = 0.7) +
  #  geom_point(data = SnailKite, aes(x = longitude, y = latitude), size = 0.1)+
  scale_color_gradient(low="#440154", 
                       high="#FDE725",
                       trans = "log10",  
                       breaks = my_breaks,
                       labels = my_breaks)+
  scale_fill_gradient(low="#440154", 
                      high="#FDE725",
                      trans = "log10",  
                      breaks = my_breaks,
                      labels = my_breaks)+
  coord_sf(xlim = c(-84.5, -79.5), 
           ylim =  c(24.1, 30.9)) +
  labs(y = "Latitude",
       x = "Longitude",       
       tag = expression(bold("(a)")),
       title = "Snail kites in Florida",
       subtitle = "eBird effort in spatial sampling units",
       color = expression(Log["10"]~"lists"),
       fill = expression(Log["10"]~"lists")) +
  annotate("text", x = -80, y = 29.75, label = "Payne's \n Prairie") +
  geom_curve(data = arrow1, aes(x = x1, y = y1, xend = x2, yend = y2),
             arrow = arrow(length = unit(0.08, "inch")), size = 0.5,
             color = "red", curvature = -0.3) +
  theme_classic()+
  theme(legend.position = c(0.2,0.25),
        legend.direction = "vertical",
        legend.box.background = element_rect(colour = "black"))

#Zoom to Payne's Prairie
arrow2 <- tibble(
  x1 = c(-82.328576, -82.334182, -82.303112, -82.292372),
  x2 = c(-82.3, -82.375, -82.25, -82.2),
  y1 = c(29.619306, 29.574222, 29.606876, 29.549109),
  y2 = c(29.7, 29.475, 29.65, 29.535)
)

Fig2b <- ggplot() +
  geom_sf(data=wrapped_gridSnailKite,
          aes(color = count, 
              fill = count)) +
  geom_point(data = SnailKite, aes(x = longitude, y = latitude), 
             size = 0.5, alpha = 0.25)+
  scale_color_gradient(low="#440154", 
                       high="#FDE725",
                       trans = "log10",  
                       breaks = my_breaks,
                       labels = my_breaks)+
  scale_fill_gradient(low = alpha("#440154", 0.25), 
                      high = alpha("#FDE725", 0.25),
                      trans = "log10",  
                      breaks = my_breaks,
                      labels = my_breaks)+
  coord_sf(xlim = c(-82.475, -82.125), 
           ylim =  c(29.45, 29.725),
           expand = T) +
  labs(y = "Latitude",
       x = "Longitude",       
       tag = expression(bold("(b)")),
       title = "Snail kites in Payne's Prairie wetland",
       subtitle = "with eBird records and popular localities") +
  annotate("text", x = -82.25, y = 29.71, label = "Sweetwater Wetlands \n Park") +
  annotate("text", x = -82.4, y = 29.475, label = "US-441") +
  annotate("text", x = -82.2, y = 29.65, label = "La Chua trail") +
  annotate("text", x = -82.2, y = 29.525, label = "Wacahoota trail") +
  geom_curve(data = arrow2, aes(x = x1, y = y1, xend = x2, yend = y2),
             arrow = arrow(length = unit(0.08, "inch")), size = 0.5,
             color = "red", curvature = -0.3) +
  theme_classic()+
  theme(legend.position = "none")

Fig2 <- grid.arrange(Fig2a, Fig2b, ncol = 2, widths = c(1, 2))

ggsave("results/Fig2_SnailKitesMap.pdf", 
       plot = Fig2, dpi = 300, width = 10, height = 5, units = "in")

ggsave("results/Fig2_SnailKitesMap.png", 
       plot = Fig2, dpi = 300, width = 10, height = 5, units = "in")
```

#### 4.2 Standardized monitoring - a benchmark to compare eBird

Joining eBird and standardized monitoring (our benchmark) in data set
`snailkites.PP`.

First, load saved data from eBird, identifying the cell with more
checklists. Recall that different packages in `R` might include same name for different functions, that could generate conflict for replicability (e.g., `select()` in packages `MASS` and `dplyr`). To avoid confusion, users can add the name of the package (e.g., `dplyr::select()`)

```
#load saved data
SnailKite <- readRDS("data_tmp/SnailKiteCellsID_filtered.rds")
snailkites.week <- readRDS("data_tmp/SnailKiteCellsWeek.rds")
snailkites.week.counts <- readRDS("data_tmp/SnailKiteCellsCountsWeek.rds")

#Cell with more values
CellTop <- SnailKite |>
  group_by(cell) |>
  mutate(n_checklists = n()) |>
  ungroup() |>
  filter(n_checklists == max(n_checklists)) |>
  dplyr::select(cell) |>
  unique()
CellTop #ID of the cell with more records - Payne's Prairie
```

Call the data from Poli *et al.* (2020),
and organize in the same way that eBird data (the highest value per week since January 2018).

```
#The monitoring data of Poli et al. (2020)
Poli.etal <- data.frame(observation.date = c("2018-02-19",
                                             "2018-03-12",
                                             "2018-04-09",
                                             "2018-05-31",
                                             "2018-06-04",
                                             "2018-07-16",
                                             "2018-08-07",
                                             "2018-08-27",
                                             "2018-10-15",
                                             "2018-12-17"),
                        abundance.monitored = c(1,
                                                2,
                                                4,
                                                6,
                                                8,
                                                6,
                                                6,
                                                12,
                                                7,
                                                29))

Poli.etal <- Poli.etal |>
  mutate(observation.date = ymd(observation.date),
         year = year(observation.date),
         week = week(observation.date),
         Time.t = case_when(year == 2018 ~ week,
                            year > 2018 ~ week+(52*(year-2018)))) |>
  dplyr::select(!observation.date)
```

Call the data form standardized monitoring in the project Snail Kites
(since 2019). This data is also organized by the high count per week, as
the eBird data.

```
snail.kite.project.pp <- read_csv("data_raw/Snail kite surveys on PP and PPC 2018_2024.csv") |> 
  mutate(observation.date = mdy(date),
         year = year(observation.date),
         week = week(observation.date),
         Time.t = case_when(year == 2018 ~ week,
                            year > 2018 ~ week+(52*(year-2018)))) |>
  group_by(Time.t, year, week) |>
  summarise(abundance.monitored = max(count))
```

```
## Rows: 53 Columns: 4
## ── Column specification ────────────────────────────────────────────────────────
## Delimiter: ","
## chr (3): location, date, survey_num
## dbl (1): count
## 
## ℹ Use `spec()` to retrieve the full column specification for this data.
## ℹ Specify the column types or set `show_col_types = FALSE` to quiet this message.
## `summarise()` has grouped output by 'Time.t', 'year'. You can override using the `.groups` argument.
```

Combine with the published data to have a single Standardized Monitoring
abundance data set, named `snail.kite.project.pp`. Here is important to
be sure that the data set is organized by the variable `Time.t` (week
since `2018-01-01` id in our case), applying the function `arrange()`.

```
#including Poli et al.
snail.kite.project.pp <- snail.kite.project.pp |>
  full_join(Poli.etal) |>
  arrange(Time.t)
```

```
## Joining with `by = join_by(Time.t, year, week, abundance.monitored)`
```

Now we can unify the data sets in a single object. First, we filter the
`snailkites.week.counts` object by the id of the cell that contains
Payne’s Prairie (`CellTop$cell`). To avoid conflict with gaps in the time
series for `observation.date`, we generated an object of dates from the
minimum value of `snailkites.week$observation_date`, and join it to the
subset of Snail kites high count per week in Payne’s Praire
(`snailkites.paynesp`).

```
#Generate the time-series for the cell with more records
snailkites.paynesp <- snailkites.week.counts |>
  filter(cell == CellTop$cell)
head(snailkites.paynesp)
```

```
#To return dates from original data
datesPP <- snailkites.week |> 
  group_by(Time.t) |> 
  summarise(observation.date = min(observation_date))
head(datesPP)
```

```
tail(datesPP)
```

```
snailkites.paynesp <- snailkites.paynesp |>
  left_join(datesPP)
```

```
## Joining with `by = join_by(Time.t)`
```

```
head(snailkites.paynesp)
```

Finally, we add the standardized monitoring (sorted by week id `Time.t`).
**CAUTION** if you do not sort by time, some functions will crash and not
run correctly, also make sure that you have only one value per time-step
selected (in our case, weeks).

```
#Add standardized monitored
snailkites.PP <- snailkites.paynesp |>
  left_join(snail.kite.project.pp, by = "Time.t") |>
  arrange(Time.t)
snailkites.PP
```

Note that the `abundance.monitored` variable, from the standardized
monitored surveys, have many `NA` values when compared with the
`Observed.y`, which is the weekly high-counts in eBird.

We can save the backup.

```
#save backup
saveRDS(snailkites.PP, file = "data_tmp/snailkitesPP.rds")
saveRDS(datesPP, file = "data_tmp/datesTimeseriesPaynesPrairie.rds")
```

#### 4.3 Visual comparison of the two time series

And we can see both time-series graphically

```
datesPP <- readRDS("data_tmp/datesTimeseriesPaynesPrairie.rds")
snailkites.PP <- readRDS("data_tmp/snailkitesPP.rds")

#Figure
snailkites.PP |>
  pivot_longer(cols = !c(Time.t, cell, year, week, observation.date),
               names_to = "group",
               values_to = "Abundance") |> 
  drop_na(Abundance) |>
  ggplot(aes(x = observation.date, 
             y = Abundance, 
             fill = group, 
             shape = group))+
    geom_segment(aes(y = 0, 
                     yend = Abundance,
                     color = group), 
                 alpha = 0.5) +
    geom_point(color = "black", 
               alpha = 0.6)+
    geom_vline(xintercept = snailkites.PP$observation.date[91], 
             color = "gray", linetype = "dotted") +
    geom_hline(yintercept = c(5,32),
               linetype = "dashed",
               color = "red")+
    labs(x = "Observation date",
       y = "Observed weekly high counts",
       title = "Time series contrast",
       tag = "",
       fill = "",
       color = "",
       shape = "")+
  scale_shape_manual(values = c(24, 21),
                       labels = c("Standardized Monitored",
                                  "eBird observations")) +
    scale_color_manual(values = c("#fc8d5995", 
                                  "#91bfdb95"),
                       labels = c("Standardized Monitored",
                                  "eBird observations"))+
    scale_fill_manual(values = c("#fc8d5995", 
                                  "#91bfdb95"),
                       labels = c("Standardized Monitored",
                                  "eBird observations"))+
  theme_classic()+
    theme(legend.position = c(0.15, 0.8))
```

```
## Warning: A numeric `legend.position` argument in `theme()` was deprecated in ggplot2
## 3.5.0.
## ℹ Please use the `legend.position.inside` argument of `theme()` instead.
## This warning is displayed once every 8 hours.
## Call `lifecycle::last_lifecycle_warnings()` to see where this warning was
## generated.
```

Figure 4.1: Time series counts per week in two data sets

Initial dynamic of the population is depicted to the left of the vertical dotted gray line (weeks 1-101; from the first week of January 2018 to the first week of December 2019). Local persistence probability estimation (\(\hat{\varphi}\)) will correspond to the weeks to the right of the dotted gray line (\(n\_{eBird} = 258\), \(n\_{SM}=32\)). Red dashed lines represent two threshold values of quasi-extinction for each dataset (\(N\_{c}^{SM} = 32\), \(N\_{c}^{eBird} = 5\)), one half of the mean observed counts of the entire time series.

### 5 Viability Population Monitoring framework and local persistence estimation - \(\hat{\varphi}\)

In the next section, we provide a step by step example and the iterated
process for each week estimation of local persistence probability
(\(\hat{\varphi}\)).

#### 5.1 Fit population dynamics model to a first part of the time-series

Let’s fit an EGSS model for the first two years – weeks from 1 to 101, n = 92, matching Standardized monitoring data (17 weeks of overlap), including the published data (Poli *et al.*, 2020).

First, load the data saved and filter `Time.t` between `1:101`.

```
datesPP <- readRDS("data_tmp/datesTimeseriesPaynesPrairie.rds")
snailkites.PP <- readRDS("data_tmp/snailkitesPP.rds")

sk.pp.init <- snailkites.PP |>
  filter(Time.t %in% c(1:101))
```

To fit a first EGSS model for the eBird data observations, we have to
adjust the data as required by the functions (vectors of log-abundance and
time-steps, in order and with a single value of log-abundance).

```
#Define variables
yt1 = sk.pp.init |>
  ungroup() |>
  arrange(Time.t) |>
  drop_na(Observed.y) |>
  dplyr::select(Observed.y) |>
  log()

#log-abundance estimate as a vector
yt1 <- yt1$Observed.y
yt1
```

```
##  [1] 0.0000000 0.0000000 0.0000000 0.0000000 0.0000000 0.0000000 1.3862944
##  [8] 1.3862944 0.6931472 0.6931472 0.6931472 1.0986123 1.0986123 1.6094379
## [15] 1.6094379 1.3862944 1.0986123 1.6094379 0.6931472 1.0986123 0.6931472
## [22] 0.0000000 1.6094379 0.0000000 0.0000000 1.3862944 1.3862944 1.3862944
## [29] 0.6931472 0.6931472 1.6094379 1.3862944 1.0986123 0.0000000 2.3978953
## [36] 2.6390573 1.9459101 1.3862944 2.0794415 2.3025851 1.7917595 2.3025851
## [43] 1.6094379 2.6390573 1.7917595 1.6094379 1.7917595 1.7917595 2.0794415
## [50] 2.4849066 2.8903718 1.3862944 2.3978953 1.7917595 2.0794415 2.0794415
## [57] 1.6094379 2.3025851 1.6094379 1.9459101 1.3862944 1.7917595 1.6094379
## [64] 0.0000000 1.3862944 1.7917595 1.0986123 1.6094379 2.0794415 1.0986123
## [71] 0.0000000 0.6931472 2.0794415 1.3862944 1.7917595 1.9459101 1.7917595
## [78] 2.3025851 2.4849066 1.3862944 2.1972246 2.0794415 2.1972246 1.9459101
## [85] 2.3025851 2.7080502 2.7725887 3.2958369 3.6888795 3.4011974 2.9444390
## [92] 2.8903718
```

```
tt1 <- sk.pp.init |>
  ungroup() |>
  arrange(Time.t) |>
  drop_na(Observed.y) |>
  dplyr::select(Time.t)

#time vector (week since 2018-01)
tt1 <- tt1$Time.t
tt1
```

```
##  [1]   8   9  10  11  12  13  14  15  16  17  18  19  20  21  22  23  24  25  26
## [20]  27  28  29  30  31  32  33  35  36  37  39  40  41  42  43  44  45  46  47
## [39]  48  49  50  51  52  53  54  55  56  57  58  59  60  61  62  63  64  65  66
## [58]  67  68  69  70  71  72  73  74  75  76  77  78  79  80  81  82  83  84  85
## [77]  86  87  88  89  90  91  92  93  94  95  96  97  98  99 100 101
```

```
#Estimate REML parameters
sk.egss.parms <- egss_reml(yt = yt1,
                            tt = tt1,
                            fguess = guess_egss(yt = yt1,
                                                tt = tt1))
print(sk.egss.parms)
```

```
## $remls
## [1]  0.03243391  0.03863836  0.23818505 -0.01604207
## 
## $lnL.hat
## [1] -83.59513
## 
## $AIC
## [1] 171.1903
```

The function `egss_reml()` fit an EGSS model and compute the Restricted
Maximum Likelihood Estimates, which are stored in the first object of
the list (named here `sk.egss.parms`). The values correspond (in
order) to the trend parameter (\(\hat{a}\_{eBird}=0.0324\)), the
environmental noise (\(\hat{\sigma^2}\_{eBird}=0.0386\)), observation
error noise (\(\hat{\tau^2}\_{eBird}=0.2382\)), and initial population
(\(\hat{x\_0}\_{eBird}=-0.0160\); note that \(e^{x\_0}\approx1\)).

With the `egss_predict()` function, we can predict the trajectory for
the EGSS model.

```
sk.egss.predict.init <- egss_predict(yt = yt1,
                                tt = tt1,
                                parms = sk.egss.parms$remls,
                                plot.it = TRUE)
```

(#fig:predict initial dynamic trajectory with model fitted)EGSS fitted to eBird

```
head(sk.egss.predict.init[[1]])
```

```
##      Time Predict.EGSS.REML Observed.y
## [1,]    8         0.9840859          1
## [2,]    9         1.0142038          1
## [3,]   10         1.0363956          1
## [4,]   11         1.0501256          1
## [5,]   12         1.0579126          1
## [6,]   13         1.0622152          1
```

As we put the `plot.it` argument as `TRUE`, we have a figure of the
predicted and observed trajectory. The `head()`
of the object shows the vector time (`Time.t`, week since January 2018,
in our case), the predicted abundance (`Predict.EGSS.REML`), and the
observed vector (`Observed.y`, the eBird weekly high counts in our
case). For our example, we want to compare with standardized monitored
surveys, so it might be convenient to change the names of this output.

We can estimate the lower and upper 95% confidence intervals of the estimation with a parametric bootstrap

```
sk.egss.predict.init.CI <- egss_pboot(B = 100, 
                                  parms = sk.egss.parms$remls,
                                  yt = yt1,
                                  tt = tt1)

plot(x = sk.egss.predict.init.CI$preds.CIs1[,1],
     y = sk.egss.predict.init.CI$preds.CIs1[,3],
     type = "l",col = "black", 
     ylab = "Abundance", xlab = "Time", main = "eBird",
     ylim = c(0,45))
lines(x = sk.egss.predict.init.CI$preds.CIs1[,1],
     y = sk.egss.predict.init.CI$preds.CIs1[,2],
     lty = 2, col = "darkgray")
lines(x = sk.egss.predict.init.CI$preds.CIs1[,1],
     y = sk.egss.predict.init.CI$preds.CIs1[,4],
     lty = 2, col = "darkgray")
points(x = sk.egss.predict.init[[1]][,1],
       y = sk.egss.predict.init[[1]][,3],
       col = "blue")
```

```
#change names to combine
colnames(sk.egss.predict.init[[1]]) <- c("Time.t", "Estimated_eBird_EGSS", "eBird.Observed")
```

We can fit the EGSS for the standardized monitored in the same way

```
#Define variables
ytSM1 = sk.pp.init |>
  ungroup() |>
  arrange(Time.t) |>
  drop_na(abundance.monitored) |>
  dplyr::select(abundance.monitored) |>
  log()

#Standardized abundance monitored as a vector
ytSM1 <- ytSM1$abundance.monitored
ytSM1
```

```
##  [1] 0.0000000 0.6931472 1.3862944 1.7917595 2.0794415 1.7917595 1.7917595
##  [8] 2.4849066 1.9459101 3.3672958 3.5835189 3.6888795 3.9702919 3.8286414
## [15] 3.8918203 4.1896547 4.3567088
```

```
ttSM1 <- sk.pp.init |>
  ungroup() |>
  arrange(Time.t) |>
  drop_na(abundance.monitored) |>
  dplyr::select(Time.t)

#corresponding time vector
ttSM1 <- ttSM1$Time.t
ttSM1
```

```
##  [1]  8 11 15 22 23 29 32 35 42 51 64 68 72 74 77 81 95
```

```
skSM.egss.parms <- egss_reml(yt = ytSM1,
                            tt = ttSM1,
                            fguess = guess_egss(yt = ytSM1,
                                                tt = ttSM1))
skSM.egss.parms
```

```
## $remls
## [1] 0.04893746 0.02954371 0.02677264 0.13109801
## 
## $lnL.hat
## [1] 11.8404
## 
## $AIC
## [1] -19.6808
```

Again, the values correspond (in order) to the trend parameter
(\(\hat{a}\_{SM}=0.0489\)), the environmental noise
(\(\hat{\sigma}\_{SM}^2=0.0295\)), observation error noise
(\(\hat{\tau}\_{SM}^2=0.0268\)), and initial population
(\(\hat{x\_0}=0.1311\)).

It is interesting that \(\hat{\sigma}\_{SM}^2\) is very similar to \(\hat{\sigma}\_{eBird}^2\) (\(\sim 0.03\)), as well as \(e^{x\_0 = 0.1311}\_{SM}\approx1\) and \(e^{x\_0 = -0.0160}\_{eBird}\approx1\). Similarly, the expected change in log-abundance per week describe lower change in eBird (\(\hat{a}\_{eBird}=0.0324\)) and higher changes in standardized monitoring weekly counts (\(\hat{a}\_{SM}=0.0489\)). In contrast, the observation error is about an order of magnitude bigger in eBird data with respect to the standardized monitoring (\(\hat{\tau}\_{SM}^2=0.0268\) \(<<\) \(\hat{\tau}\_{eBird}^2=0.2382\)).

Let’s predict the trajectory and change the column names for the
figure.

```
skSM.egss.predict.init <- egss_predict(yt = ytSM1,
                                tt = ttSM1,
                                parms = skSM.egss.parms$remls,
                                plot.it = T)
```

(#fig:predict initial dynamic trajectory SK project)EGSS fitted to SM

```
#change names to combine
colnames(skSM.egss.predict.init[[1]]) <- c("Time.t", "Estimated_SKProj_EGSS","abundance.monitored")
```

We can also compute the CI with parametric bootstrap

```
skSM.egss.predict.init.CI <- egss_pboot(B = 100, 
                                        parms = skSM.egss.parms$remls,
                                        yt = ytSM1, 
                                        tt = ttSM1)

plot(x = skSM.egss.predict.init.CI$preds.CIs1[,1],
     y = skSM.egss.predict.init.CI$preds.CIs1[,3],
     type = "l",col = "black", 
     ylab = "Abundance", xlab = "Time", main = "Standardized monitoring",
     ylim = c(0,170))
lines(x = skSM.egss.predict.init.CI$preds.CIs1[,1],
     y = skSM.egss.predict.init.CI$preds.CIs1[,2],
     lty = 2, col = "darkgray")
lines(x = skSM.egss.predict.init.CI$preds.CIs1[,1],
     y = skSM.egss.predict.init.CI$preds.CIs1[,4],
     lty = 2, col = "darkgray")
points(x = skSM.egss.predict.init[[1]][,1],
       y = skSM.egss.predict.init[[1]][,3],
       col = "blue")
```

To make the figure 3 of the main text, we combine the predicted trajectories in a single data frame to use `tidyverse` and `ggplot`.

```
sk.Estimated <- data.frame(sk.egss.predict.init) |>
  left_join(data.frame(skSM.egss.predict.init)) |>
  left_join(datesPP) #this recover the observation date per week
```

```
## Joining with `by = join_by(Time.t)`
## Joining with `by = join_by(Time.t)`
```

```
sk.Estimated_figA <- sk.Estimated |>
  pivot_longer(cols = !c(Time.t,observation.date),
               names_to = "group",
               values_to = "Abundance") |>
  drop_na(Abundance)


FigS3a <-
  ggplot(data = sk.Estimated_figA, 
         aes(x = observation.date, 
             y = Abundance))+
  geom_vline(xintercept = snailkites.PP$observation.date[92], 
             color = "gray", linetype = "dotted") +
  geom_point(data = sk.Estimated_figA |>
                filter(group %in% c("abundance.monitored",
                                      "eBird.Observed")), 
             aes(fill = factor(group,
                           levels = c("abundance.monitored",
                                      "eBird.Observed")),
                 shape = factor(group,
                           levels = c("abundance.monitored",
                                      "eBird.Observed"))),
               size = 2)+
     scale_fill_manual(values = c("#fc8d5999","#91bfdb99"),
                       labels = c("Standardized Monitoring (SM)", 
                                  "eBird"))+
    scale_shape_manual(values = c(24,21),
                       labels = c("Standardized Monitoring (SM)", 
                                  "eBird"))+
    geom_line(data = sk.Estimated_figA |>
                filter(group %in% c("Estimated_SKProj_EGSS",
                                  "Estimated_eBird_EGSS")),
              aes(color = factor(group,
                           levels = c("Estimated_SKProj_EGSS",
                                      "Estimated_eBird_EGSS"))),
              linetype = "solid", size = 1) +
   scale_color_manual(values = c("#FE996B", "#4575b4"),
                       labels = c("Predicted trajectory (SM - EGSS)", 
                                  "Predicted trajectory (eBird - EGSS)"))+
    labs(x = "Observation date",
       y = "Observed/predicted weekly high count",
#       tag = expression(bold("(a)")),
       fill = "",
       color = "",
       shape = "",
       linetype = "",
       alpha = "")+
  scale_x_date(breaks = seq(as.Date("2018-01-01"), 
                            max(sk.Estimated$observation.date), 
                            by = "3 months"), date_labels="%b \n%Y")+
   theme_classic()+
    geom_segment(aes(x = snailkites.PP$observation.date[92],
                     xend = snailkites.PP$observation.date[96],
                     y = 75,
                     yend = 75),
                 arrow = arrow(length = unit(0.2, "cm")),
                 color = "darkgray", size = 1)+
    geom_segment(aes(x = snailkites.PP$observation.date[92],
                     xend = snailkites.PP$observation.date[96],
                     y = 25,
                     yend = 25),
                 arrow = arrow(length = unit(0.2, "cm")),
                 color = "darkgray", size = 1)+
  annotate(geom = "text", label = expression(bold(varphi[italic(i)]~" estimation")),
           color = "darkgray",
           x = snailkites.PP$observation.date[93], 
           y = 50, angle = 90, vjust = 1)+
    theme(legend.position = c(0.25, 0.8), 
          legend.title = element_blank(),
        legend.spacing.y = unit(-0.5, "cm"),
        legend.background = element_blank(),
        legend.box.background = element_rect(colour = "black"))
```

```
## Warning: Using `size` aesthetic for lines was deprecated in ggplot2 3.4.0.
## ℹ Please use `linewidth` instead.
## This warning is displayed once every 8 hours.
## Call `lifecycle::last_lifecycle_warnings()` to see where this warning was
## generated.
```

```
FigS3a
```

```
## Warning in geom_segment(aes(x = snailkites.PP$observation.date[92], xend = snailkites.PP$observation.date[96], : All aesthetics have length 1, but the data has 218 rows.
## ℹ Please consider using `annotate()` or provide this layer with data containing
##   a single row.
```

```
## Warning in geom_segment(aes(x = snailkites.PP$observation.date[92], xend = snailkites.PP$observation.date[96], : All aesthetics have length 1, but the data has 218 rows.
## ℹ Please consider using `annotate()` or provide this layer with data containing
##   a single row.
```

```
## Warning in is.na(x): is.na() applied to non-(list or vector) of type
## 'expression'
```

(#fig:Figure 3a)Figure 3a - initial dynamic of the population

How much is the prediction out of the observed data in SM? Lets generate Figure 3b (we can draw the equation in the figure with our function `lm_eqn()`, removed for now).

```
#Predicted EGSS from eBird ~ Observed counts SM
lm(Estimated_eBird_EGSS~abundance.monitored, data = sk.Estimated)
```

```
## 
## Call:
## lm(formula = Estimated_eBird_EGSS ~ abundance.monitored, data = sk.Estimated)
## 
## Coefficients:
##         (Intercept)  abundance.monitored  
##             2.34423              0.07418
```

```
#Predicted EGSS from Standardized Monitored ~ Observed counts SM
lm(Estimated_SKProj_EGSS~abundance.monitored, data = sk.Estimated)
```

```
## 
## Call:
## lm(formula = Estimated_SKProj_EGSS ~ abundance.monitored, data = sk.Estimated)
## 
## Coefficients:
##         (Intercept)  abundance.monitored  
##             -0.6304               0.9286
```

```
FigS3b <- ggplot(sk.Estimated)+
  geom_abline(slope = 1)+
  geom_smooth(aes(x=abundance.monitored, 
                  y=Estimated_SKProj_EGSS),
              method = "lm", color = "#FE996B99", 
              se = F, fill = "#fc8d5905", linetype = "dashed")+
  geom_line(aes(x=abundance.monitored, y=Estimated_SKProj_EGSS), 
             color = "#FE996B", linetype = "solid", size = 1)+
#  geom_point(aes(x=abundance.monitored, y=Estimated_SKProj_EGSS), 
#             color = "black", fill = "#FE996B", shape = 24, size = 2)+
# geom_text(x = 25, y = 50, 
#            label = lm_eqn(df = sk.Estimated,
#                           x = sk.Estimated$abundance.monitored, 
#                           y = sk.Estimated$Estimated_SKProj_EGSS), 
#            parse = TRUE, color = "#fc8d59")+
  geom_smooth(aes(x=abundance.monitored, y=Estimated_eBird_EGSS),
              method = "lm", color = "#4575b499", 
              se = F, fill = "#91bfdb05", linetype = "dashed")+
  geom_line(aes(x=abundance.monitored, y=Estimated_eBird_EGSS), 
             color = "#4575b4", linetype = "solid", size = 1)+
#  geom_point(aes(x=abundance.monitored, y=Estimated_eBird_EGSS), 
#             color = "black", fill = "#4575b4", shape = 24, size = 2)+
#  geom_text(x = 50, y = 0, 
#            label = lm_eqn(df = sk.Estimated,
#                           x = sk.Estimated$abundance.monitored, 
#                           y = sk.Estimated$Estimated_eBird_EGSS), 
#            parse = TRUE, color = "#91bfdb")+
  scale_y_continuous(limits = c(0,80))+
  labs(x = "\nObserved weekly high count (SM)",
       y = "Predicted trajectory of weekly high count",
       tag = expression(bold("(b)")))+
  coord_fixed()+
  theme_classic()
FigS3b
```

```
## `geom_smooth()` using formula = 'y ~ x'
```

```
## Warning: Removed 75 rows containing non-finite outside the scale range
## (`stat_smooth()`).
```

```
## `geom_smooth()` using formula = 'y ~ x'
```

```
## Warning: Removed 75 rows containing non-finite outside the scale range
## (`stat_smooth()`).
```

```
## Warning: Removed 75 rows containing missing values or values outside the scale range
## (`geom_line()`).
## Removed 75 rows containing missing values or values outside the scale range
## (`geom_line()`).
```

(#fig:Figure 3b-1)Figure 3b - predicted vs observed trajectories

```
#and combine in the figure
FigS3 <- grid.arrange(FigS3a, FigS3b, ncol = 2, widths = c(1.5, 1))
```

```
## Warning in geom_segment(aes(x = snailkites.PP$observation.date[92], xend = snailkites.PP$observation.date[96], : All aesthetics have length 1, but the data has 218 rows.
## ℹ Please consider using `annotate()` or provide this layer with data containing
##   a single row.
```

```
## Warning in geom_segment(aes(x = snailkites.PP$observation.date[92], xend = snailkites.PP$observation.date[96], : All aesthetics have length 1, but the data has 218 rows.
## ℹ Please consider using `annotate()` or provide this layer with data containing
##   a single row.
```

```
## Warning in is.na(x): is.na() applied to non-(list or vector) of type
## 'expression'
```

```
## `geom_smooth()` using formula = 'y ~ x'
```

```
## Warning: Removed 75 rows containing non-finite outside the scale range
## (`stat_smooth()`).
```

```
## `geom_smooth()` using formula = 'y ~ x'
```

```
## Warning: Removed 75 rows containing non-finite outside the scale range
## (`stat_smooth()`).
```

```
## Warning: Removed 75 rows containing missing values or values outside the scale range
## (`geom_line()`).
## Removed 75 rows containing missing values or values outside the scale range
## (`geom_line()`).
```

(#fig:Figure 3b-2)Figure 3b - predicted vs observed trajectories

It looks not good in the `Rmd` file, but it is saved with good proportions

```
ggsave("results/Figure3ab_EstimatedTrajectory2initdays.pdf", 
       plot = FigS3a, dpi = 300, width = 8, height = 4.2, units = "in")
```

```
## Warning in geom_segment(aes(x = snailkites.PP$observation.date[92], xend = snailkites.PP$observation.date[96], : All aesthetics have length 1, but the data has 218 rows.
## ℹ Please consider using `annotate()` or provide this layer with data containing
##   a single row.
```

```
## Warning in geom_segment(aes(x = snailkites.PP$observation.date[92], xend = snailkites.PP$observation.date[96], : All aesthetics have length 1, but the data has 218 rows.
## ℹ Please consider using `annotate()` or provide this layer with data containing
##   a single row.
```

```
## Warning in is.na(x): is.na() applied to non-(list or vector) of type
## 'expression'
```

```
ggsave("results/Figure3ab_EstimatedTrajectory2initdays.png", 
       plot = FigS3a, dpi = 300, width = 8, height = 4.2, units = "in")
```

```
## Warning in geom_segment(aes(x = snailkites.PP$observation.date[92], xend = snailkites.PP$observation.date[96], : All aesthetics have length 1, but the data has 218 rows.
## ℹ Please consider using `annotate()` or provide this layer with data containing
##   a single row.
```

```
## Warning in geom_segment(aes(x = snailkites.PP$observation.date[92], xend = snailkites.PP$observation.date[96], : All aesthetics have length 1, but the data has 218 rows.
## ℹ Please consider using `annotate()` or provide this layer with data containing
##   a single row.
```

```
## Warning in is.na(x): is.na() applied to non-(list or vector) of type
## 'expression'
```

#### 5.2 Three timeframe (~5, ~10, and ~25 years) of first estimation \(\hat{\varphi}\)

With the estimated model parameters and the data during establishment of the population (up to week 101 with observations, \(i=1\) is week 101), we estimated the probability that the population declines to values below a critical
threshold (\(N\_{critical}\), as the critical number of individuals to
assume quasi-extinction), in three different periods, given timeframes, or moving simulation windows (~5 years or \(250\) weeks, ~10 years or \(500\) weeks, and ~25 years or \(1250\) weeks).

Let assume \(N\_{critical~eBird} = \frac{1}{2} (e^{\bar{y}}) = 5.47\). The probability
of persistence in each trajectory \(m\) (\(\varphi\_m\)) will be the mathematical complement of the number of time steps within the three different periods (250, 500, 1250) that the population was lower than the threshold. The resulting probability \(\varphi\_m\) is recorded for each trajectory \(m\) (\(M = 50,000\)), estimating the expected value (\(\hat{\varphi}\)), while the standard deviation (\(\sqrt{Var(\hat{\varphi})}\)), first (25%) and third (75%) quartiles as variability of the estimation per each week with data.

```
#variables:
#Log observations for the entire dataset
yt = snailkites.PP |>
  ungroup() |>
  arrange(Time.t) |>
  drop_na(Observed.y) |>
  dplyr::select(Observed.y) |>
  log()

#Vector form
yt <- yt$Observed.y
yt
```

```
##   [1] 0.0000000 0.0000000 0.0000000 0.0000000 0.0000000 0.0000000 1.3862944
##   [8] 1.3862944 0.6931472 0.6931472 0.6931472 1.0986123 1.0986123 1.6094379
##  [15] 1.6094379 1.3862944 1.0986123 1.6094379 0.6931472 1.0986123 0.6931472
##  [22] 0.0000000 1.6094379 0.0000000 0.0000000 1.3862944 1.3862944 1.3862944
##  [29] 0.6931472 0.6931472 1.6094379 1.3862944 1.0986123 0.0000000 2.3978953
##  [36] 2.6390573 1.9459101 1.3862944 2.0794415 2.3025851 1.7917595 2.3025851
##  [43] 1.6094379 2.6390573 1.7917595 1.6094379 1.7917595 1.7917595 2.0794415
##  [50] 2.4849066 2.8903718 1.3862944 2.3978953 1.7917595 2.0794415 2.0794415
##  [57] 1.6094379 2.3025851 1.6094379 1.9459101 1.3862944 1.7917595 1.6094379
##  [64] 0.0000000 1.3862944 1.7917595 1.0986123 1.6094379 2.0794415 1.0986123
##  [71] 0.0000000 0.6931472 2.0794415 1.3862944 1.7917595 1.9459101 1.7917595
##  [78] 2.3025851 2.4849066 1.3862944 2.1972246 2.0794415 2.1972246 1.9459101
##  [85] 2.3025851 2.7080502 2.7725887 3.2958369 3.6888795 3.4011974 2.9444390
##  [92] 2.8903718 2.6390573 2.4849066 2.7080502 2.7080502 2.7080502 2.9444390
##  [99] 2.9444390 2.7725887 3.1780538 2.4849066 2.7725887 2.0794415 2.7080502
## [106] 4.0943446 2.3978953 2.3025851 2.8903718 1.7917595 2.7725887 1.7917595
## [113] 2.3025851 2.3978953 2.3978953 2.3978953 2.3025851 2.0794415 1.7917595
## [120] 2.1972246 1.9459101 1.9459101 1.0986123 2.0794415 1.6094379 1.0986123
## [127] 1.3862944 1.7917595 1.6094379 1.6094379 1.7917595 0.6931472 2.0794415
## [134] 1.7917595 1.3862944 1.0986123 1.7917595 2.1972246 2.6390573 1.7917595
## [141] 2.0794415 2.4849066 1.6094379 3.5553481 3.2188758 2.4849066 2.9957323
## [148] 3.3672958 3.0910425 3.2580965 3.2580965 2.0794415 2.7725887 2.3025851
## [155] 2.0794415 2.4849066 1.3862944 2.4849066 1.3862944 1.3862944 2.0794415
## [162] 1.9459101 1.9459101 1.6094379 0.0000000 1.0986123 0.0000000 0.0000000
## [169] 1.0986123 1.6094379 0.6931472 0.6931472 1.0986123 0.0000000 0.6931472
## [176] 1.7917595 1.0986123 1.7917595 0.0000000 0.6931472 2.0794415 1.0986123
## [183] 1.6094379 1.6094379 0.6931472 1.0986123 1.7917595 1.3862944 1.0986123
## [190] 2.0794415 1.7917595 2.4849066 3.4011974 2.5649494 1.9459101 2.1972246
## [197] 2.4849066 2.0794415 2.4849066 2.0794415 1.7917595 1.9459101 2.0794415
## [204] 2.0794415 1.7917595 2.0794415 1.6094379 2.9957323 1.7917595 1.9459101
## [211] 1.9459101 2.3025851 2.3978953 2.9957323 2.4849066 2.3025851 2.3978953
## [218] 2.0794415 3.1354942 2.9957323 2.6390573 3.4011974 2.3025851 1.7917595
## [225] 2.3025851 2.4849066 2.0794415 3.2188758 3.4965076 2.3025851 1.9459101
## [232] 3.4011974 2.9444390 2.7725887 2.9957323 3.0445224 2.7080502 3.9120230
## [239] 3.7612001 3.2188758 3.9318256 3.5553481 3.7376696 3.4011974 3.2188758
## [246] 3.0910425 3.0445224 3.2188758 2.3978953 4.3174881 2.7080502 2.7080502
## [253] 2.9957323 2.9444390 2.9957323 2.9957323 3.2188758 3.2188758 3.4011974
## [260] 2.9957323 2.9957323 3.1354942 2.4849066 2.7080502 3.0445224 3.2188758
## [267] 3.0910425 2.9957323 3.2188758 2.7725887 3.2188758 2.9957323 2.7080502
## [274] 2.4849066 2.3025851 1.6094379 1.9459101 1.0986123 1.3862944 1.7917595
## [281] 2.1972246 2.1972246 2.3978953 2.7725887 2.7725887 2.6390573 2.3025851
## [288] 2.3025851 3.4011974 2.6390573 2.7080502 2.7080502 2.3025851 3.4011974
## [295] 3.1780538 2.8332133 3.2188758 2.8332133 2.9957323 2.4849066 2.4849066
## [302] 3.2188758 2.6390573 2.0794415 2.7080502 1.6094379 2.0794415 1.7917595
## [309] 1.6094379 1.6094379 1.0986123 1.0986123 1.7917595 1.7917595 0.6931472
## [316] 0.0000000 1.3862944 0.6931472 1.3862944 1.6094379 1.0986123 1.6094379
## [323] 1.6094379 2.3025851 2.0794415 1.9459101 2.3025851 2.6390573 1.6094379
## [330] 1.7917595 0.6931472 0.6931472 0.0000000 0.6931472 0.6931472 0.6931472
## [337] 1.7917595 2.6390573 3.0910425 2.4849066 0.6931472 2.8332133 0.6931472
## [344] 2.0794415 1.7917595 1.7917595 2.8332133 1.0986123 2.9957323
```

```
tt <- snailkites.PP |>
  ungroup() |>
  arrange(Time.t) |>
  drop_na(Observed.y) |>
  dplyr::select(Time.t)

#Time vector of the time series
tt <- tt$Time.t
tt
```

```
##   [1]   8   9  10  11  12  13  14  15  16  17  18  19  20  21  22  23  24  25
##  [19]  26  27  28  29  30  31  32  33  35  36  37  39  40  41  42  43  44  45
##  [37]  46  47  48  49  50  51  52  53  54  55  56  57  58  59  60  61  62  63
##  [55]  64  65  66  67  68  69  70  71  72  73  74  75  76  77  78  79  80  81
##  [73]  82  83  84  85  86  87  88  89  90  91  92  93  94  95  96  97  98  99
##  [91] 100 101 102 103 104 105 106 107 108 109 110 111 112 113 114 115 116 117
## [109] 118 119 120 121 122 123 124 125 126 127 128 129 130 131 132 133 134 135
## [127] 136 137 138 139 140 141 142 143 144 145 146 147 148 149 150 151 152 153
## [145] 154 155 156 157 158 159 160 161 162 163 164 165 166 167 168 169 170 171
## [163] 172 173 174 175 176 177 178 179 180 181 182 183 184 186 188 189 190 191
## [181] 192 193 194 195 196 197 198 199 200 201 202 203 204 205 206 207 208 209
## [199] 210 211 212 213 214 215 216 217 218 219 220 221 222 223 224 225 226 227
## [217] 228 229 230 231 232 233 234 235 236 237 238 239 240 241 242 243 244 245
## [235] 246 247 248 249 250 251 252 253 254 255 256 257 258 259 260 261 262 263
## [253] 264 265 266 267 268 269 270 271 272 273 274 275 276 277 278 279 280 281
## [271] 282 283 284 285 286 287 288 289 290 291 292 293 294 295 296 297 298 299
## [289] 300 301 302 303 304 305 306 307 308 309 310 311 312 313 314 315 316 317
## [307] 318 319 320 321 322 323 324 325 326 327 328 329 330 331 332 333 334 335
## [325] 336 337 338 339 340 341 342 343 344 345 346 347 348 349 350 351 352 353
## [343] 354 355 356 357 358 359 360
```

```
#the establishment (matching SM data) is going up to the last position
last.tt <- 92

#Define N_critical for eBird
N.critical = round(1/2*mean(exp(yt)),0)

#number of simulations
ntraj = 50000

#last observed time (week 101 - December 2019)
l = last(tt[1:last.tt])

#Simulation length
len.sim <- c(251, 501, 1251)

#which correspond to three timeframes
timeframes <- c("250 weeks or ~5 years", 
                "500 weeks or ~10 years",
                "1250 weeks or ~25 years")
```

```
#eval=FALSE

#save 𝜑 (SD) 
phi_results <- list()

# Set the base filename
base_filename <- "supporting/FigS2_Step2_1stTrajectories_10_5"

#time of starting process
StartTime <- Sys.time()

for (i in seq_along(len.sim)){
  #to save figures in png and call them after
  png_filename <- paste0(base_filename, "_", (len.sim[i]-1), ".png")
  png(png_filename, width = 10, height = 5, units = "in", res = 300)
  
  #plot the abundance data and model estimation
  plot(tt[1:last.tt], 
     exp(yt[1:last.tt]), 
     type="b", lwd=2, cex.lab=1.25, col = "#4575b4", pch = 1,
     xlim=c(min(tt), (max(tt)*3)),
     ylim=c(0,250), 
     ylab="Observed/predicted weekly high counts", 
     xlab="Time (week since January 2018)");
  
  points(x = sk.Estimated$Time.t,
     y = sk.Estimated$Estimated_eBird_EGSS,
     type = "b", col = "#91bfdb", pch = 2)

  thres.times <- as.numeric(0:(len.sim[i]-1))
  len <- max(thres.times)+1
  
  sim.mat.eBird <- egss_sim(ntraj,
                          tt = thres.times,
                          parms = sk.egss.parms$remls)
  phi <- rep(0, ntraj) 
  last.points <- rep(0, ntraj)

  for(n in 1:ntraj){
    Pop.sim <- exp(c(yt[last.tt], sim.mat.eBird[-1,n]));
    last.points[n] <- Pop.sim[len] 
 
    #How many times each trajectory go below the threshold?
    below.threshold <- sum(Pop.sim < N.critical) 
    phi[n] <- 1-(below.threshold/len)
    
        if( phi[n]>=0.5){
      lines(l+thres.times, Pop.sim, col="#d3d3d308", lty = "solid")
    }else{lines(l+thres.times, Pop.sim, col="#FF000008", lty = "solid")
    }
  }
    #add critical value line
  abline(h=N.critical, lty=2, lwd=1);
     
  #Expected value of probability of local persistence, and SD
  phi_mean <- mean(phi)
  phi_SD <- sqrt(var(phi))
  phi_1st <- as.numeric(quantile(phi, probs = 0.25))
  phi_3rd <- as.numeric(quantile(phi, probs = 0.75))
  
  phi_results[[timeframes[i]]] <- cbind(phi_mean,phi_SD, phi_1st, phi_3rd)
  
  #Create a kernel density estimate (KDE) of the `last.points` results (library(ke31d))
  kde.sims <- kde1d(x=last.points);
  
  #add the distribution to the plot
  #above the threshold
  lines(x = (kde.sims$values*(ntraj/5) + 
               l + last(thres.times)),
      y = kde.sims$grid_points,
      col="#d3d3d398", lwd=2, lty=1)

    lines(x = (kde.sims$values[kde.sims$grid_points <= N.critical]*(ntraj/5) + 
                 l + last(thres.times)),
      y = kde.sims$grid_points[kde.sims$grid_points <= N.critical],
      col="#FF000098", lwd=2, lty=1)
    
    title(main=paste0("𝜑 = ", 
                  signif(phi_mean,2),
                  " (1st Q: ",
                  signif(phi_1st,2),
                  ", 3rd Q: ",
                  signif(phi_3rd,2),
                  ") ; 1st EGSS through week ", tt[last.tt],
                  ", projected for ~", 
                  signif((len.sim[i]-1)/52, 1), 
                  " year(s)"), 
      cex=1.5)
    dev.off();
    
    print(cbind(tt[last.tt], phi_1st, phi_mean, phi_3rd, len.sim[i]-1))
}

#time process finished
EndTime <- Sys.time()
#difference of time (time lasted in the process)
timelast <- EndTime-StartTime
timelast
  
#Results for the first week in three timeframes
phi_results
```

We can see graphically the trajectories and the difference of \(\varphi\) estimated for the week 101. The figures show the observed weekly high counts (blue circles), the fitted model (light blue triangles), 50,000 trajectories (lines; those with
\(\varphi<0.5\) in reddish, else in gray), \(N\_{critical}\) as horizontal dashed
line, and the kernel density estimates of the last point of the
trajectories (reddish below \(N\_{critical}\), else gray).

#### 5.3 Add the next observations, repeat model fit, projection, and \(\hat{\varphi}\)

We can add the next week of observations (week 102; second week of December 2019) to the time-series, re-estimate model parameters as well as the probability of crashing below the \(N\_{critical}\) during the three timeframes (250, 500, 1250 weeks). Here, users can try to fit the OUSS model the following weeks, and decided by the density dependence parameter \(\hat{\theta}\) to fit the EGSS model instead. We provide below the example, but the main results focused on EGSS, which is the expected dynamic in this expanded population.

```
#The next week: 101+1
last.tt <- which(tt==102)

yt[1:last.tt] #It was already converted to the log-abundance
```

```
##  [1] 0.0000000 0.0000000 0.0000000 0.0000000 0.0000000 0.0000000 1.3862944
##  [8] 1.3862944 0.6931472 0.6931472 0.6931472 1.0986123 1.0986123 1.6094379
## [15] 1.6094379 1.3862944 1.0986123 1.6094379 0.6931472 1.0986123 0.6931472
## [22] 0.0000000 1.6094379 0.0000000 0.0000000 1.3862944 1.3862944 1.3862944
## [29] 0.6931472 0.6931472 1.6094379 1.3862944 1.0986123 0.0000000 2.3978953
## [36] 2.6390573 1.9459101 1.3862944 2.0794415 2.3025851 1.7917595 2.3025851
## [43] 1.6094379 2.6390573 1.7917595 1.6094379 1.7917595 1.7917595 2.0794415
## [50] 2.4849066 2.8903718 1.3862944 2.3978953 1.7917595 2.0794415 2.0794415
## [57] 1.6094379 2.3025851 1.6094379 1.9459101 1.3862944 1.7917595 1.6094379
## [64] 0.0000000 1.3862944 1.7917595 1.0986123 1.6094379 2.0794415 1.0986123
## [71] 0.0000000 0.6931472 2.0794415 1.3862944 1.7917595 1.9459101 1.7917595
## [78] 2.3025851 2.4849066 1.3862944 2.1972246 2.0794415 2.1972246 1.9459101
## [85] 2.3025851 2.7080502 2.7725887 3.2958369 3.6888795 3.4011974 2.9444390
## [92] 2.8903718 2.6390573
```

```
tt[1:last.tt]
```

```
##  [1]   8   9  10  11  12  13  14  15  16  17  18  19  20  21  22  23  24  25  26
## [20]  27  28  29  30  31  32  33  35  36  37  39  40  41  42  43  44  45  46  47
## [39]  48  49  50  51  52  53  54  55  56  57  58  59  60  61  62  63  64  65  66
## [58]  67  68  69  70  71  72  73  74  75  76  77  78  79  80  81  82  83  84  85
## [77]  86  87  88  89  90  91  92  93  94  95  96  97  98  99 100 101 102
```

```
#fit the OUSS
OUSS.partial <- ouss_reml(yt = yt[1:last.tt],
                           tt = tt[1:last.tt],
                           fguess = guess_ouss(yt = yt[1:last.tt],
                                               tt = tt[1:last.tt]))

OUSS.partial$remls
```

```
## [1] 1.492068e+00 4.163928e-07 4.356977e-02 2.339288e-01
```

The function `ouss_reml()` fit the OUSS model and estimate parameters. In order, the values correspond to the mean stationary distribution log-abundance (\(\hat{\mu}\_{eBird}=1.4921\)); the trend, speed of equilibration, rate to approach stationary distribution, or density dependence parameter (\(\hat{\theta}\_{eBird}=4.163928\*10^{-7}\)); the environmental noise (\(\hat{\sigma}^2\_{SM}=0.0436\)); and the observation error noise (\(\hat{\tau}^2\_{SM}=0.2339\)). Note that the density dependence parameter \(\hat{\theta}<0.025\), which suggests density-independence dynamic and the use of the EGSS model instead.

```
model <- if(OUSS.partial$remls[2] < 0.025){
  "EGSS"}else{
    "OUSS"}

model
```

```
## [1] "EGSS"
```

```
#Lets fit EGSS model partial for second observation (week 102)
EGSS.partial <- egss_reml(yt = yt[1:last.tt], 
                         tt = tt[1:last.tt],
                         fguess = guess_egss(yt = yt[1:last.tt],
                                             tt = tt[1:last.tt]))
EGSS.partial
```

```
## $remls
## [1]  0.02981099  0.03824735  0.23768876 -0.01442101
## 
## $lnL.hat
## [1] -84.33227
## 
## $AIC
## [1] 172.6645
```

```
sk.egss.predict1 <- egss_predict(yt = yt[1:last.tt],
                                         tt = tt[1:last.tt],
                                         parms = EGSS.partial$remls,
                                 plot.it = F)

head(sk.egss.predict1[[1]])
```

```
##      Time Predict.EGSS.REML Observed.y
## [1,]    8         0.9856825          1
## [2,]    9         1.0133450          1
## [3,]   10         1.0336965          1
## [4,]   11         1.0462829          1
## [5,]   12         1.0534193          1
## [6,]   13         1.0573602          1
```

Remember that `Time` is the week in our case, and `Observed.y` the
weekly high count in eBird. Let’s change column names to combine with
the `datesPP`.

```
colnames(sk.egss.predict1[[1]]) <- c("Time.t", "Estimated_eBird_EGSS","eBird.Observed")

sk.Estimated.1 <- data.frame(sk.egss.predict1) |>
  left_join(datesPP)
```

```
## Joining with `by = join_by(Time.t)`
```

And we can run the simulations with the new model updated, saving the
\(\hat{\varphi}\) values and their standard deviation for three timeframes.

```
#eval=FALSE
l = last(tt[1:last.tt])

#save 𝜑 (SD) 
phi_results <- list()

# Set the base filename
base_filename <- "supporting/FigS3_Step3_2ndTrajectories_10_5"

StartTime <- Sys.time()

for (i in seq_along(len.sim)){
  
  #to save figures in png and call them after
  png_filename <- paste0(base_filename, "_", len.sim[i]-1, ".png")
  png(png_filename, width = 10, height = 5, units = "in", res = 300)
  
  #plot the abundance data and model estimation
  plot(tt[1:last.tt], 
     exp(yt[1:last.tt]), 
     type="b", lwd=2, cex.lab=1.25, col = "#4575b4", pch = 1,
     xlim=c(min(tt), (max(tt)*3)),
     ylim=c(0,250), 
     ylab="Observed/predicted weekly high counts", 
     xlab="Time (week since January 2018)");
  
  points(x = sk.Estimated.1$Time.t,
     y = sk.Estimated.1$Estimated_eBird_EGSS,
     type = "b", col = "#91bfdb", pch = 2)

  thres.times <- as.numeric(0:(len.sim[i]-1))
  len <- max(thres.times)+1
  
  sim.mat.eBird <- egss_sim(ntraj,
                          tt = thres.times,
                          parms = EGSS.partial$remls)
  phi <- rep(0, ntraj) 
  last.points <- rep(0, ntraj)

  for(n in 1:ntraj){
    Pop.sim <- exp(c(yt[last.tt], sim.mat.eBird[-1,n]));
    last.points[n] <- Pop.sim[len] 
 
    #How many times each trajectory go below the threshold?
    below.threshold <- sum(Pop.sim < N.critical) 
    phi[n] <- 1-(below.threshold/len)
    
        if(phi[n]>=0.5){
      lines(l+thres.times, Pop.sim, col="#d3d3d308", lty = "solid")
    }else{lines(l+thres.times, Pop.sim, col="#FF000008", lty = "solid")
    }
  }
    #add critical value line
  abline(h=N.critical, lty=2, lwd=1);
  
  #Expected value of probability of local persistence, and SD
  phi_mean <- mean(phi)
  phi_SD <- sqrt(var(phi))
  phi_1st <- as.numeric(quantile(phi, probs = 0.25))
  phi_3rd <- as.numeric(quantile(phi, probs = 0.75))
  
  phi_results[[timeframes[i]]] <- cbind(phi_mean,phi_SD, phi_1st, phi_3rd)
     
  #Create a kernel density estimate (KDE) of the `last.points` results (library(ke31d))
  kde.sims <- kde1d(x=last.points);
  
    #add the distribution to the plot
  #above the threshold
  lines(x = (kde.sims$values*(ntraj/5) + 
               l + 
               last(thres.times)),
      y = kde.sims$grid_points,
      col="#d3d3d398", lwd=2, lty=1)

    lines(x = (kde.sims$values[kde.sims$grid_points <= N.critical] * 
                 ntraj + l + last(thres.times)),
      y = kde.sims$grid_points[kde.sims$grid_points <= N.critical],
      col="#FF000098", lwd=2, lty=1)
    
    title(main=paste0("𝜑 = ", 
                  signif(phi_mean,2),
                  " (1st Q: ",
                  signif(phi_1st,2),
                  ", 3rd Q: ",
                  signif(phi_3rd,2),
                  ") ; 2nd EGSS through week ", tt[last.tt],
                  ", projected for ~", 
                  signif((len.sim[i]-1)/52, 1), 
                  " year(s)"), 
      cex=1.5)
    
    dev.off()
    print(cbind(tt[last.tt], phi_1st, phi_mean, phi_3rd, len.sim[i]-1))
}
#time process finished
EndTime <- Sys.time()
#difference of time (time lasted in the process)
timelast <- EndTime-StartTime
timelast

phi_results
```

And we can see the new trajectories projected in three timeframes.

#### 5.4 Iterate the process

##### 5.4.1 Iterate the process with eBird contrasting OUSS-EGSS (some figures as examples)

Let’s iterate the process for some weeks that share data in eBird
and Standardized Monitored. This is going to change the ending position;
instead of manually change 101 to 102, we use a vector of the “ending
positions” and iterate from a sequence of length of that vector.

```
#eval=FALSE
#It last an hour
# Log observations for the entire dataset
yt = snailkites.PP |>
  ungroup() |>
  arrange(Time.t) |>
  drop_na(Observed.y) |>
  dplyr::select(Observed.y) |>
  log()

# Vector form
yt <- yt$Observed.y
yt

tt <- snailkites.PP |>
  ungroup() |>
  arrange(Time.t) |>
  drop_na(Observed.y) |>
  dplyr::select(Time.t)

# Time vector of the time series
tt <- tt$Time.t
tt

# Define N_critical for eBird
N.critical = round((1/2)*mean(exp(yt)), 0)

# List of end positions to be used
which(tt == 101) #week 101 - around 2019-12-10 - is position 92 in tt
which(tt == 211) #week 211 - around 2022-01-15 - is position 200 in tt
which(tt == 316) #week 316 - around 2024-01-22 - is position 305 in tt

end_positions <- c(92,200,305) 

#save 𝜑 (SD) 
phi_results <- vector("list", length = length(end_positions))

#save model used
modelSS <- vector("list", length = length(end_positions))

#save parameters
parms <- vector("list", length = length(end_positions))

# Set the base filename for figures
base_filename <- "supporting/FigS4_Step4_IterateTraj_10_5"

# Start process timing
StartTime <- Sys.time()

for (j in seq_along(end_positions)) {
  
  last.tt <- end_positions[j]
  l <- tt[last.tt]

  for (i in seq_along(len.sim)){
      # Set up plot for all three timeframes
  png_filename <- paste0(base_filename, 
                         "_endpos_", l, 
                         "_timeframe_",(len.sim[i]-1),
                         ".png")
  png(png_filename, width = 10, height = 5, units = "in", res = 300)

    plot(tt[1:last.tt],
         exp(yt[1:last.tt]),
         type="b", lwd=2, cex.lab=1.25, col = "#4575b4", pch = 1,
         xlim=c(min(tt), (max(tt)*3)),
         ylim=c(0,250), 
         ylab="Observed/predicted weekly high counts", 
         xlab="Time (week since January 2018)");
    
    OUSS.partial <- ouss_reml(yt = yt[1:last.tt],
                               tt = tt[1:last.tt],
                               fguess = guess_ouss(yt = yt[1:last.tt],
                                                   tt = tt[1:last.tt]))
    
    model <- if(OUSS.partial$remls[2] < 0.025){
      "EGSS"
    }else{
      "OUSS"
    }
    modelSS[[j]][[timeframes[i]]] <- model
    
    if(model == "OUSS"){
      
      parms[[j]][[timeframes[i]]] <- OUSS.partial$remls
      
      parcial.predict <- ouss_predict(yt = yt[1:last.tt],
                                      tt = tt[1:last.tt],
                                      parms = OUSS.partial$remls,
                                      plot.it = F)
      
      points(x = (parcial.predict)[[1]][,1],
             y = (parcial.predict)[[1]][,2],
             type = "b", col = "#91bfdb", pch = 2)
      
      thres.times <- as.numeric(0:(len.sim[i]-1))
      len <- max(thres.times) + 1
      
      sim.mat.eBird <- ouss_sim(ntraj, 
                                tt = thres.times, 
                                parms = OUSS.partial$remls)
      
      phi <- rep(0, ntraj)
      last.points <- rep(0, ntraj)
      
      for(n in 1:ntraj){
        Pop.sim <- exp(c(yt[last.tt], sim.mat.eBird[-1,n]));
        last.points[n] <- Pop.sim[len] 
      
      # How many times each trajectory go below the threshold?
        below.threshold <- sum(Pop.sim < N.critical)
        phi[n] <- 1-(below.threshold/len)
        
        if(phi[n] >= 0.5){
          lines(l + thres.times, Pop.sim, col="#d3d3d308", lty = "solid")
          } else {
            lines(l + thres.times, Pop.sim, col="#FF000008", lty = "solid")
            }
    }
      # Add critical value line
      abline(h=N.critical, lty=2, lwd=1);
      
      #Expected value of probability of local persistence, and SD
      phi_mean <- mean(phi)
      phi_SD <- sqrt(var(phi))
      phi_1st <- as.numeric(quantile(phi, probs = 0.25))
      phi_3rd <- as.numeric(quantile(phi, probs = 0.75))
      
      phi_results[[timeframes[i]]] <- cbind(phi_mean,phi_SD, phi_1st, phi_3rd)

      # Create a kernel density estimate (KDE) of the `last.points` results (library(ke31d))
      kde.sims <- kde1d(x=last.points);
      
      # Add the distribution to the plot
      lines(x = (kde.sims$values*(ntraj/5) + l + last(thres.times)),
            y = kde.sims$grid_points,
            col="#d3d3d398", lwd=2, lty=1)
      
      lines(x = (kde.sims$values[kde.sims$grid_points <= N.critical]*(ntraj/5) +
                   l + last(thres.times)),
            y = kde.sims$grid_points[kde.sims$grid_points <= N.critical],
            col="#FF000098", lwd=2, lty=1)
      
      title(main=paste0("𝜑 = ", 
                      signif(phi_mean,2),
                      " (1st Q: ",
                      signif(phi_1st,2),
                      ", 3rd Q: ",
                      signif(phi_3rd,2),
                      "); ", model, "; through week ", tt[last.tt],
                      ", projected for ~", 
                      signif((len.sim[i]-1)/52, 1), 
                      " year(s)"), 
          cex=1.5)
    }
    
    else{
      
      EGSS.partial <- egss_reml(yt = yt[1:last.tt],
                                 tt = tt[1:last.tt],
                                 fguess = guess_egss(yt = yt[1:last.tt],
                                                     tt = tt[1:last.tt]));
      
      parms[[j]][[timeframes[i]]] <- EGSS.partial$remls
      
      parcial.predict <- egss_predict(yt = yt[1:last.tt],
                                      tt = tt[1:last.tt],
                                      parms = EGSS.partial$remls,
                                      plot.it = F)
      
      points(x = (parcial.predict)[[1]][,1],
             y = (parcial.predict)[[1]][,2],
             type = "b", col = "#91bfdb", pch = 2)
      
      thres.times <- as.numeric(0:(len.sim[i]-1))
      len <- max(thres.times) + 1
      
      sim.mat.eBird <- egss_sim(ntraj, 
                                tt = thres.times, 
                                parms = EGSS.partial$remls)
      
      phi <- rep(0, ntraj)
      last.points <- rep(0, ntraj)
      
      for(n in 1:ntraj){
        Pop.sim <- exp(c(yt[last.tt], sim.mat.eBird[-1,n]));
        last.points[n] <- Pop.sim[len] 
      
      # How many times each trajectory go below the threshold?
        below.threshold <- sum(Pop.sim < N.critical)
        phi[n] <- 1-(below.threshold/len)
        
        if(phi[n] >= 0.5){
          lines(l+thres.times, Pop.sim, col="#d3d3d308", lty = "solid")
          } else {
            lines(l+thres.times, Pop.sim, col="#FF000008", lty = "solid")
            }
    }
      # Add critical value line
      abline(h=N.critical, lty=2, lwd=1);
      
      #Expected value of probability of local persistence, and SD
      phi_mean <- mean(phi)
      phi_SD <- sqrt(var(phi))
      phi_1st <- as.numeric(quantile(phi, probs = 0.25))
      phi_3rd <- as.numeric(quantile(phi, probs = 0.75))
      
      phi_results[[timeframes[i]]] <- cbind(phi_mean,phi_SD, phi_1st, phi_3rd)
      
      # Create a kernel density estimate (KDE) of the `last.points` results (library(ke31d))
      kde.sims <- kde1d(x=last.points);
      
      # Add the distribution to the plot
      lines(x = (kde.sims$values*(ntraj/5) + l + last(thres.times)),
            y = kde.sims$grid_points,
            col="#d3d3d398", lwd=2, lty=1)
      
      lines(x = (kde.sims$values[kde.sims$grid_points <= N.critical]*(ntraj/5) + 
                   l + last(thres.times)),
            y = kde.sims$grid_points[kde.sims$grid_points <= N.critical],
            col="#FF000098", lwd=2, lty=1)
      
      title(main=paste0("𝜑 = ", 
                      signif(phi_mean,2),
                      " (1st Q: ",
                      signif(phi_1st,2),
                      ", 3rd Q: ",
                      signif(phi_3rd,2),
                      "); ", model, "; through week ", tt[last.tt],
                      ", projected for ~", 
                      signif((len.sim[i]-1)/52, 1), 
                      " year(s)"), 
          cex=1.5)
    }
    
    dev.off()
  
  }
  print(cbind(l, model, phi_1st, phi_mean, phi_3rd, phi_SD, len.sim[i]-1))
}

#time process finished
EndTime <- Sys.time()
#difference of time (time lasted in the process)
timelast <- EndTime-StartTime
timelast
```

###### 5.4.1.1 Trajectories in timeframe of ~5 years.

Let’s see the examples.

How is changing in 5 moments \(\hat{\varphi}\_{eBird\ \sim 5\ years~simulated}\)?

Again, it is not expected for the population to be at stationary distribution, unless we adjust a new initial population of the time series (see Dennis & Ponciano for more details on the OUSS model).

###### 5.4.1.2 Trajectories in timeframe of ~10 years.

How is changing in 5 moments \(\hat{\varphi}\_{eBird\ \sim 10~years~simulated}\)?

The pattern selected on week 211 keeps selecting OUSS, likely misidentifying density-dependence from the eBird data.

###### 5.4.1.3 Trajectories in timeframe of ~25 years.

How is changing in three moments \(\hat{\varphi}\_{eBird\ \sim 25\ years~simulated}\)?

Note that for the week 211 (January 2022) the population model fitted is the density-dependence model (OUSS). However, this might be do to the fact that lower and less variable weekly high counts are similar to the initial population (1), misidentifying density dependence dynamic in this expanding population. We include this example to show the users how to apply the contrast for established populations (see also Section 8 - Other populations of snail kites).

##### 5.4.2 Iterate each week to extract \(\hat{\varphi}\) and SD from eBird contrasting OUSS-EGSS (no figures)

```
#Change at the end to `{r eval=FALSE}`; it might take long time (~9 hrs)

#the data
snailkites.PP <- readRDS("data_tmp/snailkitesPP.rds")

#Log observations for the entire data set
yt = snailkites.PP |>
  ungroup() |>
  arrange(Time.t) |>
  drop_na(Observed.y) |>
  dplyr::select(Observed.y) |>
  log()

#Vector form
yt <- yt$Observed.y
yt

# Define N_critical for eBird
N.critical = round((1/2)*mean(exp(yt)),0)

tt <- snailkites.PP |>
  ungroup() |>
  arrange(Time.t) |>
  drop_na(Observed.y) |>
  dplyr::select(Time.t)

#Time vector of the time series
tt <- tt$Time.t
tt

#the first two years length
last.tt <- 87 

#Number of trajectories and bootstraps (CIs)
ntraj = 50000

# End positions to modeling (steps)
end_positions <- which(tt > last.tt)

#check weeks ID selected as `end_positions`
tt[end_positions]

#save 𝜑 (SD) 
phi_results <- vector("list", length = length(end_positions))

#save model used
modelSS <- vector("list", length = length(end_positions))

# Start process timing
StartTime <- Sys.time()

for (j in seq_along(end_positions)) {
  
  last.tt <- end_positions[j]
  l <- tt[last.tt]

  for (i in seq_along(len.sim)){
    
    OUSS.partial <- ouss_reml(yt = yt[1:last.tt],
                               tt = tt[1:last.tt],
                               fguess = guess_ouss(yt = yt[1:last.tt],
                                                   tt = tt[1:last.tt]))
    
    model <- if(OUSS.partial$remls[2] < 0.025){
      "EGSS"
    }else{
      "OUSS"
    }
    modelSS[[j]][[timeframes[i]]] <- model
    
    if(model == "OUSS"){
      
      thres.times <- as.numeric(0:(len.sim[i]-1))
      len <- max(thres.times) + 1
      
      sim.mat.eBird <- ouss_sim(ntraj, 
                                tt = thres.times, 
                                parms = OUSS.partial$remls)
      
      phi <- rep(0, ntraj)
      last.points <- rep(0, ntraj)
      
      for(n in 1:ntraj){
        Pop.sim <- exp(c(yt[last.tt], sim.mat.eBird[-1,n]));
        last.points[n] <- Pop.sim[len] 
      
      # How many times each trajectory go below the threshold?
        below.threshold <- sum(Pop.sim < N.critical)
        phi[n] <- 1-(below.threshold/len)
      }
      
      #Expected value of probability of local persistence, and SD
      phi_mean <- mean(phi)
      phi_SD <- sqrt(var(phi))
      phi_1st <- as.numeric(quantile(phi, probs = 0.25))
      phi_3rd <- as.numeric(quantile(phi, probs = 0.75))
  
      phi_results[[j]][[timeframes[i]]] <- cbind(phi_1st, phi_mean, phi_3rd, phi_SD)
      
    }
    else{
      
      EGSS.partial <- egss_reml(yt = yt[1:last.tt],
                                 tt = tt[1:last.tt],
                                 fguess = guess_egss(yt = yt[1:last.tt],
                                                     tt = tt[1:last.tt]));
      
      thres.times <- as.numeric(0:(len.sim[i]-1))
      len <- max(thres.times) + 1
      
      sim.mat.eBird <- egss_sim(ntraj, 
                                tt = thres.times, 
                                parms = EGSS.partial$remls)
      
      phi <- rep(0, ntraj)
      last.points <- rep(0, ntraj)
      
      for(n in 1:ntraj){
        Pop.sim <- exp(c(yt[last.tt], sim.mat.eBird[-1,n]));
        last.points[n] <- Pop.sim[len] 
      
      # How many times each trajectory go below the threshold?
        below.threshold <- sum(Pop.sim < N.critical)
        phi[n] <- 1-(below.threshold/len)
      }
      
      #Expected value of probability of local persistence, and SD
      phi_mean <- mean(phi)
      phi_SD <- sqrt(var(phi))
      phi_1st <- as.numeric(quantile(phi, probs = 0.25))
      phi_3rd <- as.numeric(quantile(phi, probs = 0.75))
  
      phi_results[[j]][[timeframes[i]]] <- cbind(phi_1st, phi_mean, phi_3rd, phi_SD)
      
    }
  }
  print(cbind(l, model, 
              phi_1st, phi_mean,phi_3rd,
              phi_SD, 
              len.sim[i]-1))
}

#time process finished
EndTime <- Sys.time()
#difference of time (time lasted in the process)
timelast <- EndTime-StartTime
timelast
```

...

This is a longer process due to our time step scale of weeks and simulation window of 25 years (\(\sim3\) days; currently is `{r eval=FALSE}`), so we save the `phi_results` (with `phi_1st`, `phi_hat`, `phi_3rd`, and `phi_SD`), and `modelSS` vectors as data frames to make a plot.

```
#creating data frames for phi_mean, phi_SD, and modelSS
phi_eBird_df <- do.call(rbind, lapply(seq_along(phi_results), function(j) {
  do.call(rbind, lapply(seq_along(phi_results[[j]]), function(i) {
    data.frame(
      Time.t = tt[end_positions[j]],
      Timeframe = timeframes[i],
      phi.1st = phi_results[[j]][[i]][1],
      phi.hat = phi_results[[j]][[i]][2],
      phi.3rd = phi_results[[j]][[i]][3],
      SD.phi = phi_results[[j]][[i]][4],
      Model = modelSS[[j]][[i]]
    )
  }))
}))

# Convert to factors for better plotting
phi_eBird_df$Timeframe <- factor(phi_eBird_df$Timeframe, levels = timeframes)
head(phi_eBird_df, n = 12)

saveRDS(phi_eBird_df, "results/phi_hat_SD_eBird_OUSS_EGSS.rds")
```

##### 5.4.3 Iterate each week to extract \(\hat{\varphi}\) and variation from eBird only EGSS (no figures)

```
#Change at the end to `{r eval=FALSE}`; it might take long time

#the data
snailkites.PP <- readRDS("data_tmp/snailkitesPP.rds")

#Log observations for the entire data set
yt = snailkites.PP |>
  ungroup() |>
  arrange(Time.t) |>
  drop_na(Observed.y) |>
  dplyr::select(Observed.y) |>
  log()

#Vector form
yt <- yt$Observed.y
yt

# Define N_critical for eBird
N.critical = round((1/2)*mean(exp(yt)),0)

tt <- snailkites.PP |>
  ungroup() |>
  arrange(Time.t) |>
  drop_na(Observed.y) |>
  dplyr::select(Time.t)

#Time vector of the time series
tt <- tt$Time.t
tt

#the first two years length
last.tt <- which(tt == 101) 

#Simulation length (1, 2, and 5 generations)
len.sim <- c(251, 501, 1251)

#which correspond to three timeframes
timeframes <- c("250 weeks or ~5 years", 
                "500 weeks or ~10 years",
                "1250 weeks or ~25 years")

#Number of trajectories and bootstraps (CIs)
ntraj = 50000

# End positions to modeling (steps)
end_positions <- which(tt >= 101)

#check weeks ID selected as `end_positions`
tt[end_positions]

#save 𝜑 (SD) 
phi_results <- vector("list", length = length(end_positions))

#save model used
modelSS <- vector("list", length = length(end_positions))

#save model parameters
parms <- vector("list", length = length(end_positions))

# Start process timing
StartTime <- Sys.time()

for (j in seq_along(end_positions)) {
  
  last.tt <- end_positions[j]
  l <- tt[last.tt]

  for (i in seq_along(len.sim)){
    
    model <- "EGSS"
    
    modelSS[[j]][[timeframes[i]]] <- model
    
      EGSS.partial <- egss_reml(yt = yt[1:last.tt],
                                 tt = tt[1:last.tt],
                                 fguess = guess_egss(yt = yt[1:last.tt],
                                                     tt = tt[1:last.tt]));
      
      parms[[j]][[timeframes[i]]] <- EGSS.partial$remls
      
      thres.times <- as.numeric(0:(len.sim[i]-1))
      len <- max(thres.times) + 1
      
      sim.mat.eBird <- egss_sim(ntraj, 
                                tt = thres.times, 
                                parms = EGSS.partial$remls)
      
      phi <- rep(0, ntraj)
      last.points <- rep(0, ntraj)
      
      for(n in 1:ntraj){
        Pop.sim <- exp(c(yt[last.tt], sim.mat.eBird[-1,n]));
        last.points[n] <- Pop.sim[len] 
      
      # How many times each trajectory go below the threshold?
        below.threshold <- sum(Pop.sim < N.critical)
        phi[n] <- 1-(below.threshold/len)
      }
      
      #Expected value of probability of local persistence, and SD
      phi_mean <- mean(phi)
      phi_SD <- sqrt(var(phi))
      phi_1st <- as.numeric(quantile(phi, probs = 0.25))
      phi_3rd <- as.numeric(quantile(phi, probs = 0.75))
  
      phi_results[[j]][[timeframes[i]]] <- cbind(phi_1st, phi_mean, phi_3rd, phi_SD)
      
    
  }
  print(cbind(l, model, 
              phi_1st,
              phi_mean,
              phi_3rd,
              phi_SD, 
              len.sim[i]-1))
}

#time process finished
EndTime <- Sys.time()
#difference of time (time lasted in the process)
timelast <- EndTime-StartTime
timelast
```

...

This is a longer process (\(\sim3\) days; currently is `{r eval=FALSE}`), so we saved the `phi_results` (with `phi_1st`, `phi_hat`, `phi_3rd`, and `phi_SD`), and `modelSS` vectors as data frames to make a plot. We also saved the four parameters in each week `parms` list.

```
#creating data frames for phi_mean, phi_SD, modelSS, and parms
phi_eBird_df <- do.call(rbind, lapply(seq_along(phi_results), function(j) {
  do.call(rbind, lapply(seq_along(phi_results[[j]]), function(i) {
    data.frame(
      Time.t = tt[end_positions[j]],
      Timeframe = timeframes[i],
      phi.1st = phi_results[[j]][[i]][1],
      phi.hat = phi_results[[j]][[i]][2],
      phi.3rd = phi_results[[j]][[i]][3],
      SD.phi = phi_results[[j]][[i]][4],
      Model = modelSS[[j]][[i]],
      ln.lambda = parms[[j]][[i]][1],
      sigma.sqr = parms[[j]][[i]][2],
      tau.sqr = parms[[j]][[i]][3],
      x0 = parms[[j]][[i]][4]
    )
  }))
}))

# Convert to factors for better plotting
phi_eBird_df$Timeframe <- factor(phi_eBird_df$Timeframe, levels = timeframes)
head(phi_eBird_df, n = 9)
tail(phi_eBird_df, n = 9)

saveRDS(phi_eBird_df, "results/phi_hat_SD_eBird_model_parameters.rds")
```

##### 5.4.4 Iterate the process with Standardized Monitoring data (figure examples)

Let iterates for the periods that share data in eBird and Standardized
Monitored (data available in standardized monitored). Here we use the comparison of density-dependence vs density-independence dynamics in each iteration.

```
#Standardized monitored data
#Log observations for the entire dataset
ytSM = snailkites.PP |>
  ungroup() |>
  arrange(Time.t) |>
  drop_na(abundance.monitored) |>
  dplyr::select(abundance.monitored) |>
  log()

#Vector form
ytSM <- ytSM$abundance.monitored
ytSM
```

```
##  [1] 0.0000000 0.6931472 1.3862944 1.7917595 2.0794415 1.7917595 1.7917595
##  [8] 2.4849066 1.9459101 3.3672958 3.5835189 3.6888795 3.9702919 3.8286414
## [15] 3.8918203 4.1896547 4.3567088 4.7095302 4.8040210 5.0172798 4.8040210
## [22] 4.7184989 4.1431347 4.4886364 4.4067192 3.9702919 2.4849066 2.7725887
## [29] 3.7135721 3.9120230 4.3438054 4.3307333 4.5849675 4.5108595 4.7273878
## [36] 4.6821312 5.5093883 4.8903491 5.0751738 4.6443909 4.7449321 4.6634391
## [43] 4.2626799 3.8501476 3.7376696 4.2195077 1.7917595 1.3862944 0.6931472
```

```
#N crticial
N.criticalSM <- round((1/2)*mean(exp(ytSM)),0)

ttSM <- snailkites.PP |>
  ungroup() |>
  arrange(Time.t) |>
  drop_na(abundance.monitored) |>
  dplyr::select(Time.t)

#Time vector of the time series
ttSM <- ttSM$Time.t
ttSM
```

```
##  [1]   8  11  15  22  23  29  32  35  42  51  64  68  72  74  77  81  95 112 116
## [20] 118 122 124 128 129 148 164 168 170 211 215 219 222 225 228 231 234 251 263
## [39] 267 271 274 277 280 283 286 303 316 320 324
```

And run the model

```
#This will take ~4 hours, consider change `{r eval=FALSE}`

ntraj =50000

# End positions to modeling (steps)
end_positions <- which(ttSM >= 101)

#actual weeks
ttSM[end_positions]

#save 𝜑 (SD) 
phi_results <- vector("list", length = length(end_positions))

#save model used
modelSS <- vector("list", length = length(end_positions))

#save model parameters
parms <- vector("list", length = length(end_positions))

# Set the base filename for figures
base_filename <- "supporting/FigS5_IterateTrajSM_10_5" 

# Start process timing
StartTime <- Sys.time()

for (j in seq_along(end_positions)) {
  
  last.tt <- end_positions[j]
  l <- ttSM[last.tt]

  for (i in seq_along(len.sim)){
      # Set up plot for all three timeframes
  png_filename <- paste0(base_filename, 
                         "_endpos_",l, 
                         "_timeframe_",(len.sim[i]-1),
                         ".png")
  png(png_filename, width = 10, height = 5, units = "in", res = 300)

    
    plot(ttSM[1:last.tt],
         exp(ytSM[1:last.tt]),
         type="b", lwd=2, cex.lab=1.25, col = "#d73027", pch = 1,
         xlim=c(min(tt), (max(tt)*3)),
         ylim=c(0,250), 
         ylab="Observed/predicted weekly high counts", 
         xlab="Time (week since January 2018)");
    
    OUSS.partial <- ouss_reml(yt = ytSM[1:last.tt],
                               tt = ttSM[1:last.tt],
                               fguess = guess_ouss(yt = ytSM[1:last.tt],
                                                   tt = ttSM[1:last.tt]))
    
    model <- if(OUSS.partial$remls[2] < 0.025){
      "EGSS"
    }else{
      "OUSS"
    }
    modelSS[[j]][[timeframes[i]]] <- model
    
    if(model == "OUSS"){
      
      parms[[j]][[timeframes[i]]] <- OUSS.partial$remls
      
      parcial.predict <- ouss_predict(yt = ytSM[1:last.tt],
                                      tt = ttSM[1:last.tt],
                                      parms = OUSS.partial$remls,
                                      plot.it = F)
      
      points(x = (parcial.predict)[[1]][,1],
             y = (parcial.predict)[[1]][,2],
             type = "b", col = "#fc8d59", pch = 2)
      
      thres.times <- as.numeric(0:(len.sim[i]-1))
      len <- max(thres.times) + 1
      
      sim.mat.eBird <- ouss_sim(ntraj, 
                                tt = thres.times, 
                                parms = OUSS.partial$remls)
      
      phi <- rep(0, ntraj)
      last.points <- rep(0, ntraj)
      
      for(n in 1:ntraj){
        Pop.sim <- exp(c(ytSM[last.tt], sim.mat.eBird[-1,n]));
        last.points[n] <- Pop.sim[len] 
      
      # How many times each trajectory go below the threshold?
        below.threshold <- sum(Pop.sim < N.criticalSM)
        phi[n] <- 1-(below.threshold/len)
        
        if(phi[n] >= 0.5){
          lines(l+thres.times, Pop.sim, col="#d3d3d308", lty = "solid")
          } else {
            lines(l+thres.times, Pop.sim, col="#FF000008", lty = "solid")
            }
    }
      # Add critical value line
      abline(h=N.criticalSM, lty=2, lwd=1);
      
      #Expected value of probability of local persistence, and SD
      phi_mean <- mean(phi)
      phi_SD <- sqrt(var(phi))
      phi_1st <- as.numeric(quantile(phi, probs = 0.25))
      phi_3rd <- as.numeric(quantile(phi, probs = 0.75))
  
      phi_results[[j]][[timeframes[i]]] <- cbind(phi_1st, phi_mean, phi_3rd, phi_SD)

      # Create a kernel density estimate (KDE) of the `last.points` results (library(ke31d))
      kde.sims <- kde1d(x=last.points);
      
      # Add the distribution to the plot
      lines(x = (kde.sims$values*(ntraj/5) + l + last(thres.times)),
            y = kde.sims$grid_points,
            col="#d3d3d398", lwd=2, lty=1)
      
      lines(x = (kde.sims$values[kde.sims$grid_points <= N.criticalSM]*(ntraj/5) + 
                   l + last(thres.times)),
            y = kde.sims$grid_points[kde.sims$grid_points <= N.criticalSM],
            col="#FF000098", lwd=2, lty=1)
      
      title(main=paste0("𝜑 = ", 
                      signif(phi_mean,2),
                     " (1st Q: ",
                      signif(phi_1st,2),
                      ", 3rd Q: ",
                      signif(phi_3rd,2),
                      "); ", model, "; through week ", tt[last.tt],
                       ", projected for ~", 
                      signif((len.sim[i]-1)/52, 2), 
                      " year(s)"), 
          cex=1.5)
    }
    
    else{
      
      EGSS.partial <- egss_reml(yt = ytSM[1:last.tt],
                                 tt = ttSM[1:last.tt],
                                 fguess = guess_egss(yt = ytSM[1:last.tt],
                                                          tt = ttSM[1:last.tt]));
      
      parms[[j]][[timeframes[i]]] <- EGSS.partial$remls
      
      parcial.predict <- egss_predict(yt = ytSM[1:last.tt],
                                      tt = ttSM[1:last.tt],
                                      parms = EGSS.partial$remls,
                                      plot.it = F)
      
      points(x = (parcial.predict)[[1]][,1],
             y = (parcial.predict)[[1]][,2],
             type = "b", col = "#fc8d59", pch = 2)
      
      thres.times <- as.numeric(0:(len.sim[i]-1))
      len <- max(thres.times) + 1
      
      sim.mat.eBird <- egss_sim(ntraj, 
                                tt = thres.times, 
                                parms = EGSS.partial$remls)
      
      phi <- rep(0, ntraj)
      last.points <- rep(0, ntraj)
      
      for(n in 1:ntraj){
        Pop.sim <- exp(c(ytSM[last.tt], sim.mat.eBird[-1,n]));
        last.points[n] <- Pop.sim[len] 
      
      # How many times each trajectory go below the threshold?
        below.threshold <- sum(Pop.sim < N.criticalSM)
        phi[n] <- 1-(below.threshold/len)
        
        if(phi[n] >= 0.5){
          lines(l+thres.times, Pop.sim, col="#d3d3d308", lty = "solid")
          } else {
            lines(l+thres.times, Pop.sim, col="#FF000008", lty = "solid")
            }
    }
      # Add critical value line
      abline(h=N.criticalSM, lty=2, lwd=1);
      
      #Expected value of probability of local persistence, and SD
      phi_mean <- mean(phi)
      phi_SD <- sqrt(var(phi))
      phi_1st <- as.numeric(quantile(phi, probs = 0.25))
      phi_3rd <- as.numeric(quantile(phi, probs = 0.75))
  
      phi_results[[j]][[timeframes[i]]] <- cbind(phi_1st, phi_mean, phi_3rd, phi_SD)
  
      # Create a kernel density estimate (KDE) of the `last.points` results (library(ke31d))
      kde.sims <- kde1d(x=last.points);
      
      # Add the distribution to the plot
      lines(x = (kde.sims$values*(ntraj/5) + l + last(thres.times)),
            y = kde.sims$grid_points,
            col="#d3d3d398", lwd=2, lty=1)
      
      lines(x = (kde.sims$values[kde.sims$grid_points <= N.criticalSM]*(ntraj/5) + 
                   l + last(thres.times)),
            y = kde.sims$grid_points[kde.sims$grid_points <= N.criticalSM],
            col="#FF000098", lwd=2, lty=1)
      
      title(main=paste0("𝜑 = ", 
                      signif(phi_mean,2),
                     " (1st Q: ",
                      signif(phi_1st,2),
                      ", 3rd Q: ",
                      signif(phi_3rd,2),
                      "); ", model, "; through week ", tt[last.tt],
                       ", projected for ~", 
                      signif((len.sim[i]-1)/52, 2), 
                      " year(s)"), 
          cex=1.5)
    }
    
    dev.off()
  
  }
  print(cbind(l, model, 
              phi_1st,
              phi_mean,
              phi_3rd,
              phi_SD, 
              len.sim[i]-1))
}
#time process finished
EndTime <- Sys.time()
#difference of time (time lasted in the process)
timelast <- EndTime-StartTime
timelast
```

...

<br:

Now we save the `phi_results` and `modelSS` vectors as data
frames to make a plot.

```
#creating data frames for phi_mean, phi_SD, modelSS, and parms
phi_SM_df <- do.call(rbind, lapply(seq_along(phi_results), function(j) {
  do.call(rbind, lapply(seq_along(phi_results[[j]]), function(i) {
    data.frame(
      Time.t = ttSM[end_positions[j]],
      Timeframe = timeframes[i],
      phi.1stSM = phi_results[[j]][[i]][1],
      phi.hatSM = phi_results[[j]][[i]][2],
      phi.3rdSM = phi_results[[j]][[i]][3],
      SD.phiSM = phi_results[[j]][[i]][4],
      Model = modelSS[[j]][[i]],
      ln.lambda = parms[[j]][[i]][1],
      sigma.sqr = parms[[j]][[i]][2],
      tau.sqr = parms[[j]][[i]][3],
      x0 = parms[[j]][[i]][4]
    )
  }))
}))

# Convert to factors for better plotting
phi_SM_df$Timeframe <- factor(phi_SM_df$Timeframe, levels = timeframes)
head(phi_SM_df, n = 12)

saveRDS(phi_SM_df, "results/phi_hat_SD_SM_model_parameters.rds")
```

###### 5.4.4.1 Trajectories in timeframe of ~5 years

Let’s see the examples.

How is changing in 2 moments \(\hat{\varphi}\_{SM\ \sim five~years~simulated}\)?

###### 5.4.4.2 Trajectories in timeframe of ~10 years.

How is changing in 2 moments \(\hat{\varphi}\_{SM\ \sim ten~years~simulated}\)?

###### 5.4.4.3 Trajectories in timeframe of ~25 years.

How is changing in 2 moments \(\hat{\varphi}\_{SM\ \sim 25~years~simulated}\)?

### 6 Results - \(\hat{\varphi}\)

Call the data and establish transformation of second Y-axis (supplementary figure to include time series counts and \(\varphi\)).

```
#Load data saved
  #dates
  datesPP <- readRDS("data_tmp/datesTimeseriesPaynesPrairie.rds")
  #data organized
  snailkites.PP <- readRDS("data_tmp/snailkitesPP.rds")
  #local persistence from eBird - OUSS or EGSS
  phi.eBird.OUSS.EGSS <- readRDS("results/phi_hat_SD_eBird_OUSS_EGSS.rds") 
  #local persistence from eBird - EGSS
  phi.eBird <- readRDS("results/phi_hat_SD_eBird_model_parameters.rds") 
  #local persistence from Standardized Monitored
  phi.SM <-readRDS("results/phi_hat_SD_SM_model_parameters.rds")
  
  N.critical <- 5 # 1/2*empirical K = 1/2 mean(exp(yt))
  N.criticalSM <- 32 # 1/2*empirical K = 1/2 mean(exp(ytSM))

#coefficient to convert second Y-axis
coeff <- 100
coeffSM <- 250
```

We can compare the overlap of \(\varphi\) estimation. First we combine the `phi.eBird` and `phi.SM` results.

```
head(phi.eBird)
```

```
head(phi.eBird.OUSS.EGSS)
```

```
head(phi.SM)
```

```
#combine phi dataframes
phi_combined <- phi.eBird |>
  bind_rows(.id = "dataset", phi.eBird.OUSS.EGSS) |> 
  mutate(dataset = case_when(dataset == 1~"eBird.EGSS",
                             dataset != 1~"eBird.OUSS.EGSS")) |>
  bind_rows(.id = "original.dataset", phi.SM) |> 
  mutate(original.dataset = case_when(original.dataset == 1~"eBird",
                             original.dataset != 1~"SM"),
         dataset = case_when(original.dataset == "SM"~"SM",
                                   original.dataset != "SM"~dataset))

head(phi_combined)
```

Then, we add the dates and filter by the three Simulation windows (Timeframes)

#### 6.1 \(\hat{\varphi}\) Temporal trend - Figure 4 and extended figures

```
#add dates while filtering Timeframe
phi_data_plot <- datesPP |> 
  left_join(phi_combined |> 
              filter(Timeframe == "250 weeks or ~5 years")) |>
  ungroup() |> 
  pivot_longer(cols = c(phi.1st, phi.1stSM),
               names_to = "group.1st",
               values_to = "phi_1st") |> 
  drop_na(phi_1st) |>
  pivot_longer(cols = c(phi.hat, phi.hatSM),
               names_to = "group",
               values_to = "phi_hat") |> 
  drop_na(phi_hat) |>
  pivot_longer(cols = c(phi.3rd, phi.3rdSM),
               names_to = "group.3rd",
               values_to = "phi_3rd") |> 
  drop_na(phi_3rd) |> 
  pivot_longer(cols = c(SD.phi, SD.phiSM),
               names_to = "group.SD",
               values_to = "phi_SD") |> 
  drop_na(phi_SD)
```

```
## Joining with `by = join_by(Time.t)`
```

```
phi_data_plot
```

```
FigOUSS_A <- ggplot(data = phi_data_plot,
       aes(x = observation.date, 
           y = phi_hat, 
           color = original.dataset)) +
    geom_vline(xintercept = snailkites.PP$observation.date[92], 
             color = "gray", linetype = "dotted") +
    facet_wrap(~dataset, ncol = 3) +
    geom_linerange(aes(ymin = phi_hat-phi_SD,
                       ymax = phi_hat+phi_SD),
                   alpha = 0.15) +
    geom_linerange(aes(ymin = phi_1st,
                       ymax = phi_3rd),
                   alpha = 0.15) +
    geom_point(aes(shape = Model),
                   alpha = 0.65) +
    scale_y_continuous(expand = c(0,0))+
    scale_color_manual(values = c("#91bfdb95",
                                "#fc8d5995"),
                     labels = c(expression(italic(varphi)[eBird]),
                                expression(italic(varphi)[SM])))+
    scale_x_date(limits = c(snailkites.PP$observation.date[90],
                          max(snailkites.PP$observation.date)),
               breaks = seq(snailkites.PP$observation.date[90], 
                            max(snailkites.PP$observation.date), 
                            by = "12 months"), date_labels="%b\n%Y")+
  labs(x = "Observation date",
       y = expression(italic(varphi)),
       tag = expression(bold("(a)")),
       title = "Simulation window: 250 weeks or ~5 years",
       subtitle = "(one generation)")  +
    theme_classic() +
    theme(legend.title = element_blank(),
        legend.position = c(0.9, 0.8),
        legend.spacing.y = unit(-0.5, "cm"),
        legend.background = element_blank(),
        legend.box.background = element_rect(colour = "black"))+
  guides(color = guide_legend(ncol=2),
         shape = guide_legend(ncol=2))+
    coord_cartesian(ylim = c(0,1))
FigOUSS_A
```

```
## Warning: Removed 11 rows containing missing values or values outside the scale range
## (`geom_segment()`).
## Removed 11 rows containing missing values or values outside the scale range
## (`geom_segment()`).
```

```
## Warning: Removed 11 rows containing missing values or values outside the scale range
## (`geom_point()`).
```

```
#Only EGSS comparison

#Figure overlap phi

Figure4a <- ggplot(data = phi_data_plot |>
         filter(dataset != "eBird.OUSS.EGSS"),
       aes(x = observation.date, 
           y = phi_hat, 
           fill = original.dataset, 
           color = original.dataset, 
           shape = original.dataset)) +
    geom_vline(xintercept = snailkites.PP$observation.date[92], 
             color = "gray", linetype = "dotted") +
    geom_linerange(aes(ymin = phi_hat-phi_SD,
                       ymax = phi_hat+phi_SD),
                   alpha = 0.15) +
    geom_linerange(aes(ymin = phi_1st,
                       ymax = phi_3rd),
                   alpha = 0.15, size = 1) +
    geom_point(alpha = 0.65) +
    scale_y_continuous(expand = c(0,0))+
    scale_shape_manual(values = c(21,24),
                     labels = c(expression(italic(varphi)[eBird]),
                                expression(italic(varphi)[SM]))) +
    scale_fill_manual(values = c("#91bfdb95",
                                "#fc8d5995"),
                     labels = c(expression(italic(varphi)[eBird]),
                                expression(italic(varphi)[SM]))) +
    scale_color_manual(values = c("#91bfdb95",
                                "#fc8d5995"),
                     labels = c(expression(italic(varphi)[eBird]),
                                expression(italic(varphi)[SM]))) +
    scale_x_date(limits = c(snailkites.PP$observation.date[90],
                          max(snailkites.PP$observation.date)),
               breaks = seq(snailkites.PP$observation.date[90], 
                            max(snailkites.PP$observation.date), 
                            by = "12 months"), date_labels="%b\n%Y")+
  labs(x = "Observation date",
       y = expression(paste("Persistence (",italic(varphi),")")),
       tag = expression(bold("(a)")),
       title = "Simulation window: 250 weeks or ~5 years",
       subtitle = "(one generation)")  +
    theme_classic() +
    theme(legend.title = element_blank(),
        legend.position = c(0.85, 0.9),
        legend.spacing.y = unit(-0.5, "cm"),
        legend.background = element_blank(),
        legend.box.background = element_rect(colour = "black"))+
  guides(color = guide_legend(ncol=2),
         shape = guide_legend(ncol=2))+
    coord_cartesian(ylim = c(0,1))

Figure4a
```

```
#add dates while filtering Timeframe
phi_data_plot <- datesPP |> 
  left_join(phi_combined |> 
              filter(Timeframe == "500 weeks or ~10 years")) |>
  ungroup() |> 
  pivot_longer(cols = c(phi.1st, phi.1stSM),
               names_to = "group.1st",
               values_to = "phi_1st") |> 
  drop_na(phi_1st) |>
  pivot_longer(cols = c(phi.hat, phi.hatSM),
               names_to = "group",
               values_to = "phi_hat") |> 
  drop_na(phi_hat) |>
  pivot_longer(cols = c(phi.3rd, phi.3rdSM),
               names_to = "group.3rd",
               values_to = "phi_3rd") |> 
  drop_na(phi_3rd) |> 
  pivot_longer(cols = c(SD.phi, SD.phiSM),
               names_to = "group.SD",
               values_to = "phi_SD") |> 
  drop_na(phi_SD)
```

```
## Joining with `by = join_by(Time.t)`
```

```
phi_data_plot
```

```
FigOUSS_B <- ggplot(data = phi_data_plot,
       aes(x = observation.date, 
           y = phi_hat, 
           color = original.dataset)) +
    geom_vline(xintercept = snailkites.PP$observation.date[92], 
             color = "gray", linetype = "dotted") +
    facet_wrap(~dataset, ncol = 3) +
    geom_linerange(aes(ymin = phi_hat-phi_SD,
                       ymax = phi_hat+phi_SD),
                   alpha = 0.15) +
    geom_linerange(aes(ymin = phi_1st,
                       ymax = phi_3rd),
                   alpha = 0.15) +
    geom_point(aes(shape = Model),
                   alpha = 0.65) +
    scale_y_continuous(expand = c(0,0))+
    scale_color_manual(values = c("#91bfdb95",
                                "#fc8d5995"),
                     labels = c(expression(italic(phi)[eBird]),
                                expression(italic(phi)[SM])))+
    scale_x_date(limits = c(snailkites.PP$observation.date[90],
                          max(snailkites.PP$observation.date)),
               breaks = seq(snailkites.PP$observation.date[90], 
                            max(snailkites.PP$observation.date), 
                            by = "12 months"), date_labels="%b\n%Y")+
  labs(x = "Observation date",
       y = expression(italic(varphi)),
       tag = expression(bold("(b)")),
       title = "Simulation window: 500 weeks or ~10 years",
       subtitle = "(two generations)")  +
    theme_classic() +
    theme(legend.title = element_blank(),
        legend.position = "none",
        legend.spacing.y = unit(-0.5, "cm"),
        legend.background = element_blank(),
        legend.box.background = element_rect(colour = "black"))+
  guides(color = guide_legend(ncol=2),
         shape = guide_legend(ncol=2))+
    coord_cartesian(ylim = c(0,1))

FigOUSS_B
```

```
## Warning: Removed 11 rows containing missing values or values outside the scale range
## (`geom_segment()`).
## Removed 11 rows containing missing values or values outside the scale range
## (`geom_segment()`).
```

```
## Warning: Removed 11 rows containing missing values or values outside the scale range
## (`geom_point()`).
```

```
#Only EGSS comparison

#Figure overlap phi

Figure4b <- ggplot(data = phi_data_plot |>
         filter(dataset != "eBird.OUSS.EGSS"),
       aes(x = observation.date, 
           y = phi_hat, 
           color = original.dataset, 
           fill = original.dataset, 
           shape = original.dataset)) +
    geom_vline(xintercept = snailkites.PP$observation.date[92], 
             color = "gray", linetype = "dotted") +
    geom_linerange(aes(ymin = phi_hat-phi_SD,
                       ymax = phi_hat+phi_SD),
                   alpha = 0.15) +
    geom_linerange(aes(ymin = phi_1st,
                       ymax = phi_3rd),
                   alpha = 0.15, size = 1) +
    geom_point(alpha = 0.65) +
    scale_y_continuous(expand = c(0,0))+
    scale_shape_manual(values = c(21,24),
                     labels = c(expression(italic(phi)[eBird]),
                                expression(italic(phi)[SM])))+
    scale_color_manual(values = c("#91bfdb95",
                                "#fc8d5995"),
                     labels = c(expression(italic(phi)[eBird]),
                                expression(italic(phi)[SM])))+
    scale_fill_manual(values = c("#91bfdb95",
                                "#fc8d5995"),
                     labels = c(expression(italic(phi)[eBird]),
                                expression(italic(phi)[SM])))+
    scale_x_date(limits = c(snailkites.PP$observation.date[90],
                          max(snailkites.PP$observation.date)),
               breaks = seq(snailkites.PP$observation.date[90], 
                            max(snailkites.PP$observation.date), 
                            by = "12 months"), date_labels="%b\n%Y")+
  labs(x = "Observation date",
       y = expression(paste("Persistence (",italic(varphi),")")),
       tag = expression(bold("(b)")),
       title = "Simulation window: 500 weeks or ~10 years",
       subtitle = "(two generations)")  +
    theme_classic() +
    theme(legend.title = element_blank(),
        legend.position = "none",
        legend.spacing.y = unit(-0.5, "cm"),
        legend.background = element_blank(),
        legend.box.background = element_rect(colour = "black"))+
  guides(color = guide_legend(ncol=2),
         shape = guide_legend(ncol=2))+
    coord_cartesian(ylim = c(0,1))

Figure4b
```

```
#add dates while filtering Timeframe
phi_data_plot <- datesPP |> 
  left_join(phi_combined |> 
              filter(Timeframe == "1250 weeks or ~25 years")) |>
  ungroup() |> 
  pivot_longer(cols = c(phi.1st, phi.1stSM),
               names_to = "group.1st",
               values_to = "phi_1st") |> 
  drop_na(phi_1st) |>
  pivot_longer(cols = c(phi.hat, phi.hatSM),
               names_to = "group",
               values_to = "phi_hat") |> 
  drop_na(phi_hat) |>
  pivot_longer(cols = c(phi.3rd, phi.3rdSM),
               names_to = "group.3rd",
               values_to = "phi_3rd") |> 
  drop_na(phi_3rd) |> 
  pivot_longer(cols = c(SD.phi, SD.phiSM),
               names_to = "group.SD",
               values_to = "phi_SD") |> 
  drop_na(phi_SD)
```

```
## Joining with `by = join_by(Time.t)`
```

```
phi_data_plot
```

```
FigOUSS_C <- ggplot(data = phi_data_plot,
       aes(x = observation.date, 
           y = phi_hat, 
           color = original.dataset)) +
    geom_vline(xintercept = snailkites.PP$observation.date[92], 
             color = "gray", linetype = "dotted") +
    facet_wrap(~dataset, ncol = 3) +
    geom_linerange(aes(ymin = phi_hat-phi_SD,
                       ymax = phi_hat+phi_SD),
                   alpha = 0.15) +
    geom_linerange(aes(ymin = phi_1st,
                       ymax = phi_3rd),
                   alpha = 0.15) +
    geom_point(aes(shape = Model),
                   alpha = 0.65) +
    scale_y_continuous(expand = c(0,0))+
    scale_color_manual(values = c("#91bfdb95",
                                "#fc8d5995"),
                     labels = c(expression(italic(phi)[eBird]),
                                expression(italic(phi)[SM])))+
    scale_x_date(limits = c(snailkites.PP$observation.date[90],
                          max(snailkites.PP$observation.date)),
               breaks = seq(snailkites.PP$observation.date[90], 
                            max(snailkites.PP$observation.date), 
                            by = "12 months"), date_labels="%b\n%Y")+
  labs(x = "Observation date",
       y = expression(italic(varphi)),
       tag = expression(bold("(c)")),
       title = "Simulation window: 1250 weeks or ~25 years",
       subtitle = "(five generations)")  +
    theme_classic() +
    theme(legend.title = element_blank(),
        legend.position = "none",
        legend.spacing.y = unit(-0.5, "cm"),
        legend.background = element_blank(),
        legend.box.background = element_rect(colour = "black"))+
  guides(color = guide_legend(ncol=2),
         shape = guide_legend(ncol=2))+
    coord_cartesian(ylim = c(0,1))

FigOUSS_C
```

```
## Warning: Removed 11 rows containing missing values or values outside the scale range
## (`geom_segment()`).
## Removed 11 rows containing missing values or values outside the scale range
## (`geom_segment()`).
```

```
## Warning: Removed 11 rows containing missing values or values outside the scale range
## (`geom_point()`).
```

```
#Only EGSS comparison

#Figure overlap phi

Figure4c <- ggplot(data = phi_data_plot |>
         filter(dataset != "eBird.OUSS.EGSS"),
       aes(x = observation.date, 
           y = phi_hat, 
           color = original.dataset, 
           fill = original.dataset, 
           shape = original.dataset)) +
    geom_vline(xintercept = snailkites.PP$observation.date[92], 
             color = "gray", linetype = "dotted") +
    geom_linerange(aes(ymin = phi_hat-phi_SD,
                       ymax = phi_hat+phi_SD),
                   alpha = 0.15) +
    geom_linerange(aes(ymin = phi_1st,
                       ymax = phi_3rd),
                   alpha = 0.15, size = 1) +
    geom_point(alpha = 0.65) +
    scale_y_continuous(expand = c(0,0))+
    scale_shape_manual(values = c(21,24),
                     labels = c(expression(italic(phi)[eBird]),
                                expression(italic(phi)[SM])))+
    scale_fill_manual(values = c("#91bfdb95",
                                "#fc8d5995"),
                     labels = c(expression(italic(phi)[eBird]),
                                expression(italic(phi)[SM])))+
    scale_color_manual(values = c("#91bfdb95",
                                "#fc8d5995"),
                     labels = c(expression(italic(phi)[eBird]),
                                expression(italic(phi)[SM])))+
    scale_x_date(limits = c(snailkites.PP$observation.date[90],
                          max(snailkites.PP$observation.date)),
               breaks = seq(snailkites.PP$observation.date[90], 
                            max(snailkites.PP$observation.date), 
                            by = "12 months"), date_labels="%b\n%Y")+
  labs(x = "Observation date",
       y = expression(paste("Persistence (",italic(varphi),")")),
       tag = expression(bold("(c)")),
       title = "Simulation window: 1250 weeks or ~25 years",
       subtitle = "(five generations)")  +
    theme_classic() +
    theme(legend.title = element_blank(),
        legend.position = "none",
        legend.spacing.y = unit(-0.5, "cm"),
        legend.background = element_blank(),
        legend.box.background = element_rect(colour = "black"))+
  guides(color = guide_legend(ncol=2),
         shape = guide_legend(ncol=2))+
    coord_cartesian(ylim = c(0,1))

Figure4c
```

And combine the figures

```
#and combine in the figure
Figure4Ext <- grid.arrange(FigOUSS_A, FigOUSS_B, FigOUSS_C, 
                          ncol = 1)
```

```
## Warning: Removed 11 rows containing missing values or values outside the scale range
## (`geom_segment()`).
## Removed 11 rows containing missing values or values outside the scale range
## (`geom_segment()`).
```

```
## Warning: Removed 11 rows containing missing values or values outside the scale range
## (`geom_point()`).
```

```
## Warning: Removed 11 rows containing missing values or values outside the scale range
## (`geom_segment()`).
## Removed 11 rows containing missing values or values outside the scale range
## (`geom_segment()`).
```

```
## Warning: Removed 11 rows containing missing values or values outside the scale range
## (`geom_point()`).
```

```
## Warning: Removed 11 rows containing missing values or values outside the scale range
## (`geom_segment()`).
## Removed 11 rows containing missing values or values outside the scale range
## (`geom_segment()`).
```

```
## Warning: Removed 11 rows containing missing values or values outside the scale range
## (`geom_point()`).
```

```
#lets saved it with good proportions
ggsave("results/Figure4_Extended_OUSS_EGSS_5-10-25yrsSimulated.pdf", 
       plot = Figure4Ext, dpi = 300, width = 12, height = 10, units = "in")

ggsave("results/Figure4_Extended_OUSS_EGSS_5-10-25yrsSimulated.png", 
       plot = Figure4Ext, dpi = 300, width = 12, height = 10, units = "in")
```

```
#and combine in the figure
Figure4 <- grid.arrange(Figure4a, Figure4b, Figure4c, 
                          ncol = 1)
```

```
#lets saved it with good proportions
ggsave("results/Figure4_PLocalPersi_5-10-25yrsSimulated.pdf", 
       plot = Figure4, dpi = 300, width = 5.5, height = 9, units = "in")

ggsave("results/Figure4_PLocalPersi_5-10-25yrsSimulated.png", 
       plot = Figure4, dpi = 300, width = 5.5, height = 9, units = "in")
```

#### 6.2 \(\hat{\varphi}\) Correlation and RMSE between datasets and simulated windows

How correlated are the \(\hat{\varphi}\)?

```
correlations <- phi.eBird |> 
  dplyr::select(Time.t,
         Timeframe,
         phi.hat) |>
  full_join(phi.SM |>
              dplyr::select(Time.t,
                     Timeframe,
                     phi.hatSM)) |>
  group_by(Timeframe) |>
  summarise(correlation = cor(phi.hatSM, 
                              phi.hat, 
                              use = "pairwise.complete.obs"))
```

```
## Joining with `by = join_by(Time.t, Timeframe)`
```

```
correlations
```

All simulated windows include high Pearson correlation (\(>0.9\)). Now, we calculated the Root Mean Square Error (RMSE).

```
phi.eBird |> 
  dplyr::select(Time.t,
         Timeframe,
         phi.hat) |>
  full_join(phi.SM |>
              dplyr::select(Time.t,
                     Timeframe,
                     phi.hatSM)) |>
  group_by(Timeframe) |>
  drop_na() |>
  summarise(rmse = sqrt(mean(phi.hatSM - phi.hat)^2))
```

```
## Joining with `by = join_by(Time.t, Timeframe)`
```

Lower RMSE in \(S\_{1250}\) indicates that, on average, close persistence estimates between the two datasets. However, a good compromise between high correlation and low RMSE is provided by the window simulation of \(S\_{500}\). We can plot the Absolute error difference for overlapping weeks of counts.

```
datesPP |>
  full_join(phi.eBird |> 
  dplyr::select(Time.t,
         Timeframe,
         phi.hat)) |>
  full_join(phi.SM |>
              dplyr::select(Time.t,
                     Timeframe,
                     phi.hatSM)) |>
  group_by(Timeframe) |>
  drop_na() |>
  mutate(Abs.diff = (phi.hatSM - phi.hat)) |>
  ggplot(aes(x = observation.date, y = Abs.diff)) +
  geom_hline(yintercept = 0,
             linetype = "dashed")+
    geom_segment(aes(x = observation.date,
                     xend = observation.date, 
                     y = 0, 
                     yend = Abs.diff),
                 color = "#909090")+
  geom_point(color = "#838383")+
  scale_x_date(limits = c(snailkites.PP$observation.date[90],
                          max(snailkites.PP$observation.date)),
               breaks = seq(snailkites.PP$observation.date[90], 
                            max(snailkites.PP$observation.date), 
                            by = "12 months"), date_labels="%b\n%Y")+
  facet_wrap(~Timeframe, ncol = 1)+
  labs(x = "Observation date",
       y = "Absolute error difference")+
  theme_classic()+
  theme(strip.background = element_blank())
```

```
## Joining with `by = join_by(Time.t)`
## Joining with `by = join_by(Time.t, Timeframe)`
```

#### 6.3 Model parameters time series and RMSE

In each iteration \(i\), we save the four parameters of the models. We can see the timeseries of estimation of these parameters for the two datasets.

```
phi_combined |>
  filter(dataset %in% c("eBird.EGSS", "SM")) |>
  mutate(N_0 = round(exp(x0)),
         original.dataset = if_else(dataset == "SM",
                           "Standardized monitoring",
                           dataset)) |>
  rename(Trend.parameter = ln.lambda,
         Env.noise = sigma.sqr,
         Obs.noise = tau.sqr) |>
  pivot_longer(cols = c(Trend.parameter, 
                        Env.noise,
                        Obs.noise,
                        N_0),
               names_to = "Parameter",
               values_to = "Estimate") |>
  unique() |>
  filter(Timeframe == "500 weeks or ~10 years") |>
  left_join(datesPP) |> 
  ggplot(aes(x = observation.date, 
             y = Estimate, 
             color = original.dataset))+
    geom_point(alpha = 0.6)+
    facet_wrap(~factor(Parameter,
                        levels = c("Trend.parameter",
                                   "Env.noise",
                                   "Obs.noise",
                                   "N_0"),
                       labels = c("ln(lambda)",
                                  "sigma^2",
                                  "tau^2",
                                  "N[0] == e^(X[0])")), 
               ncol = 2, 
               scales = "free_y", labeller = label_parsed) +
    scale_color_manual(values = c("#91bfdb95",
                                  "#fc8d5995")) +
  labs(x = "Observation date",
       color = "Dataset")+
  theme_classic()+
  theme(legend.position = "bottom",
        strip.background = element_blank())
```

```
## Joining with `by = join_by(Time.t)`
```

Very similar trends for the different parameters! We can see the evidently difference of the observation noise parameter (\(\hat{\tau}^2\)), way higher for eBird data. We can also explore the difference calculating the Root Mean Square Error between the two time series. Recall that there are four parameters in the EGSS:

```
phi.SM |>
  dplyr::select(Time.t,
                ln.lambda,
                sigma.sqr,
                tau.sqr,
                x0) |>
    mutate(init.pop = round(exp(x0))) |>
    unique() |>
  left_join(phi.eBird |>
              dplyr::select(Time.t,
                            ln.lambda,
                            sigma.sqr, 
                            tau.sqr,
                            x0) |>
              mutate(init.pop = round(exp(x0))) |>
              unique(),
            by = c("Time.t")) |> 
  #summary() #to see the means and ranges for each parameter; 
    #x = SM, y = eBird
  summarise(rmse.ln.lambda = sqrt(mean(ln.lambda.x - ln.lambda.y)^2),
            rmse.sigma.sqr = sqrt(mean(sigma.sqr.x - sigma.sqr.y)^2),
            rmse.tau.sqr = sqrt(mean(tau.sqr.x - tau.sqr.y)^2),
            rmse.N_0 = sqrt(mean(init.pop.x - init.pop.y)^2))
```

After converting back the \(\hat{x\_0}\) parameter to individuals (\(N\_0\)), the estimation for the two datasets is the same (\(\text{RMSE}\_{\hat{x\_0}} = 0\)). Similarly, lower values of \(\text{RMSE}\_{\hat{\sigma}^2}=0.0014\) indicate similar environmental noise estimated for the two datasets in weeks of overlap in monitoring. The following parameter with similar estimation is the trend parameter \(\text{RMSE}\_{\hat{\ln{\lambda}}}=0.0087\). Finally, the parameter of more difference between the datasets is the the observation noise error, \(\text{RMSE}\_{\hat{\tau}^2}=0.2007\).

We can, additionally, plot the Absolute error difference for overlapping weeks of counts for these four parameters.

```
phi.SM |>
  dplyr::select(Time.t,
                ln.lambda,
                sigma.sqr,
                tau.sqr,
                x0) |>
    mutate(init.pop = round(exp(x0))) |>
    unique() |>
  left_join(phi.eBird |>
              dplyr::select(Time.t,
                            ln.lambda,
                            sigma.sqr, 
                            tau.sqr,
                            x0) |>
              mutate(init.pop = round(exp(x0))) |>
              unique(),
            by = c("Time.t")) |>
  left_join(datesPP) |>
  mutate(Trend = (ln.lambda.x - ln.lambda.y),
         Env.noise = (sigma.sqr.x - sigma.sqr.y),
         Obs.noise = (tau.sqr.x - tau.sqr.y),
         N_0 = (init.pop.x - init.pop.y)) |>
  pivot_longer(cols = c(Trend,
                        Env.noise,
                        Obs.noise,
                        N_0), 
               names_to = "Parameter",
               values_to = "Abs.diff") |>
  ggplot(aes(x = observation.date, 
             y = Abs.diff)) +
    geom_segment(aes(x = observation.date,
                     xend = observation.date, 
                     y = 0, 
                     yend = Abs.diff),
                 color = "#909090")+
  geom_point(color = "#838383")+
  geom_hline(yintercept = 0, color = "#838383",
             linetype = "dotted")+
  scale_x_date(limits = c(snailkites.PP$observation.date[78],
                          max(snailkites.PP$observation.date)),
               breaks = seq(snailkites.PP$observation.date[90], 
                            max(snailkites.PP$observation.date), 
                            by = "12 months"), date_labels="%b\n%Y")+
  facet_wrap(~factor(Parameter,
                        levels = c("Trend",
                                   "Env.noise",
                                   "Obs.noise",
                                   "N_0"),
                       labels = c("ln(lambda)",
                                  "sigma^2",
                                  "tau^2",
                                  "N[0] == e^(X[0])")), 
               ncol = 2, 
               scales = "free_y", labeller = label_parsed)+
  labs(x = "Observation date",
       y = "Absolute error difference")+
  theme_classic()+
  theme(strip.background = element_blank())
```

```
## Joining with `by = join_by(Time.t)`
```

Negative values in the absolute error difference indicate a higher estimate in the parameter using eBird data compared to using standardized monitoring.

#### 6.4 Results for simulation window of ~5 years (250 weeks)

```
#Extended figures

phi_data_plot <- snailkites.PP |> 
  left_join(phi.eBird |>
              full_join(phi.SM)  |> 
              filter(Timeframe == "250 weeks or ~5 years")) |>
  ungroup() |>
    mutate(phi.hat = phi.hat * coeff,
           SD.phi = SD.phi * coeff,
           phi.hatSM = phi.hatSM * coeffSM,
           SD.phiSM = SD.phiSM * coeffSM) |> 
  pivot_longer(cols = !c(cell, 
                         year, 
                         week, 
                         Timeframe, 
                         Time.t, 
                         observation.date, 
                         Model,
                         ln.lambda,
                         sigma.sqr,
                         tau.sqr,
                         x0),
               names_to = "group",
               values_to = "phi_values") |> 
  drop_na(phi_values)
```

```
## Joining with `by = join_by(Time.t, Timeframe, Model, ln.lambda, sigma.sqr,
## tau.sqr, x0)`
## Joining with `by = join_by(Time.t)`
```

```
phi_data_plot
```

```
#Ribbon lo-hi phi (SD) for the two datasets
ribbon_phi_eBird <- phi_data_plot |>
  filter(group %in% c("phi.hat", "SD.phi")) |>
  pivot_wider(names_from = group, values_from = phi_values, values_fn = {mean}) |>
  mutate(phi_values = phi.hat)

ribbon_phi_SM <- phi_data_plot |>
  filter(group %in% c("phi.hatSM", "SD.phiSM")) |>
  pivot_wider(names_from = group, values_from = phi_values, values_fn = {mean})|>
  mutate(phi_values = phi.hatSM)

  #figurea
Fig3aExt <- ggplot(phi_data_plot |> 
                  filter(group %in% c("phi.hat", "Observed.y")), 
                aes(x = observation.date,
                    y = phi_values,
                    fill = group)) +
  geom_vline(xintercept = snailkites.PP$observation.date[92], 
             color = "gray", linetype = "dashed") +
  geom_ribbon(data = ribbon_phi_eBird,
              aes(ymin = phi.hat-SD.phi, ymax = phi.hat+SD.phi),
              fill = "#91bfdb25") +
#  geom_line(aes(color = group, linetype = group)) +
  geom_segment(data = phi_data_plot |> 
                  filter(group %in% c("Observed.y")),
               aes(x = observation.date,
                   xend = observation.date,
                   y = 0,
                   yend = phi_values),
             color = "#4575b445",
             fill = "#4575b445") +
  geom_point(data = phi_data_plot |> 
                  filter(group %in% c("Observed.y")),
             color = "#4575b445", size = 0.5, fill = "#4575b445", shape = 21)+
  geom_point(data = phi_data_plot |> 
                  filter(group %in% c("phi.hat")) |>
               drop_na(Model),
             shape = 21, fill = "#91bfdb95", color = "black") +
  labs(x = "Observation date",
       y = "Observed weekly high counts",
       tag = expression(bold("(a)")),
       fill = "",
       color = "",
       linetype = "",
       shape = "") +
  scale_y_continuous(sec.axis = sec_axis(trans = ~./coeff, 
                                         name = expression("eBird " * italic(varphi)))) +
  coord_cartesian(ylim = c(0, coeff))+
  scale_x_date(limits = c(min(datesPP$observation.date),
                          max(snailkites.PP$observation.date)),
               breaks = seq(min(datesPP$observation.date),
                          max(snailkites.PP$observation.date), 
                            by = "12 months"), date_labels="%b\n%Y")+
  scale_linetype_manual(values = c("solid",
                                   "solid"),
                     labels = c("eBird weekly high count", 
                                expression("P(local persistence) = " * italic(varphi))))+
  scale_color_manual(values = c("#4575b445", 
                                "#91bfdb45"),
                     labels = c("eBird weekly high count", 
                                expression("P(local persistence) = " * italic(varphi)))) +
  geom_hline(yintercept = N.critical, linetype = "dashed", color = "red")+
  theme_classic() +
  theme(legend.title = element_blank(),
        legend.position = c(0.125,0.7),
        legend.background = element_blank(),
        legend.box.background = element_rect(colour = "black"),
        axis.line.y.right = element_line(color = "#91bfdb"),
        axis.ticks.y.right = element_line(color = "#91bfdb"))
```

```
## Warning in geom_segment(data = filter(phi_data_plot, group %in%
## c("Observed.y")), : Ignoring unknown parameters: `fill`
```

```
## Warning: The `trans` argument of `sec_axis()` is deprecated as of ggplot2 3.5.0.
## ℹ Please use the `transform` argument instead.
## This warning is displayed once every 8 hours.
## Call `lifecycle::last_lifecycle_warnings()` to see where this warning was
## generated.
```

```
Fig3aExt
```

```
#SM figure

  #figurea
Fig3bExt <- ggplot(phi_data_plot |> 
                  filter(group %in% c("phi.hatSM", "abundance.monitored")), 
                aes(x = observation.date,
                    y = phi_values,
                    fill = group)) +
  geom_vline(xintercept = snailkites.PP$observation.date[92], 
             color = "gray", linetype = "dashed") +
  geom_ribbon(data = ribbon_phi_SM,
              aes(ymin = phi.hatSM-SD.phiSM, ymax = phi.hatSM+SD.phiSM),
              fill = "#fc8d5925") +
#  geom_line(aes(color = group, linetype = group)) +
  geom_segment(data = phi_data_plot |> 
                  filter(group %in% c("abundance.monitored")),
               aes(x = observation.date,
                   xend = observation.date,
                   y = 0,
                   yend = phi_values),
             color = "#d7302745", 
             fill = "#d7302745")+
  geom_point(data = phi_data_plot |> 
                  filter(group %in% c("abundance.monitored")),
             color = "#d7302745", size = 0.5, fill = "#d7302745", shape = 21)+
  geom_point(data = phi_data_plot |> 
                  filter(group %in% c("phi.hatSM")) |>
               drop_na(Model),
             shape = 21, fill = "#fc8d5995", color = "black") +
  labs(x = "Observation date",
       y = "Observed weekly high counts",
       tag = expression(bold("(b)")),
       fill = "",
       color = "",
       linetype = "",
       shape = "") +
  scale_y_continuous(sec.axis = sec_axis(trans = ~./coeffSM, 
                                         name = expression("Standardized Monitored " * italic(varphi)))) +
  coord_cartesian(ylim = c(0, coeffSM))+
  scale_x_date(limits = c(min(datesPP$observation.date),
                          max(snailkites.PP$observation.date)),
               breaks = seq(min(datesPP$observation.date),
                          max(snailkites.PP$observation.date), 
                            by = "12 months"), date_labels="%b\n%Y")+
  scale_linetype_manual(values = c("solid",
                                   "solid"),
                     labels = c("Standardized Monitored", 
                                expression("P(local persistence) = " * italic(varphi))))+
  scale_color_manual(values = c("#d7302745", 
                                "#fc8d5945"),
                     labels = c("Standardized Monitored", 
                                expression("P(local persistence) = " * italic(varphi)))) +
  geom_hline(yintercept = N.criticalSM, 
             linetype = "dashed", color = "red")+
  theme_classic() +
  theme(legend.title = element_blank(),
        legend.position = c(0.125,0.7),
        legend.background = element_blank(),
        legend.box.background = element_rect(colour = "black"),
        axis.line.y.right = element_line(color = "#fc8d59"),
        axis.ticks.y.right = element_line(color = "#fc8d59"))
```

```
## Warning in geom_segment(data = filter(phi_data_plot, group %in%
## c("abundance.monitored")), : Ignoring unknown parameters: `fill`
```

```
Fig3bExt
```

```
#Performance comparison
phidata <- phi.eBird |>
  dplyr::select(Time.t, 
                Timeframe,
                phi.hat, SD.phi) |>
              full_join(phi.SM |>
                          dplyr::select(Time.t, 
                                        Timeframe, 
                                        phi.hatSM, SD.phiSM)) |>
  filter(Timeframe == "250 weeks or ~5 years")
```

```
## Joining with `by = join_by(Time.t, Timeframe)`
```

```
#relationship?
lm(phi.hat~phi.hatSM, data = phidata)
```

```
## 
## Call:
## lm(formula = phi.hat ~ phi.hatSM, data = phidata)
## 
## Coefficients:
## (Intercept)    phi.hatSM  
##      0.2725       0.5769
```

```
Fig3cExt <- ggplot(phidata)+
    geom_abline(slope = 1)+
    geom_pointrange(aes(x = phi.hatSM, y = phi.hat, 
                        ymin = phi.hat-SD.phi, 
                        ymax = phi.hat+SD.phi),
                    shape = 23, fill = "gray", color = "gray",
                    alpha = 0.25) +
    geom_errorbarh(aes(x = phi.hatSM, y = phi.hat,
                       xmax = phi.hatSM+SD.phiSM, 
                       xmin = phi.hatSM-SD.phiSM, height = 0),
                   color = "gray",
                   alpha = 0.25)+
    geom_point(aes(x = phi.hatSM, y = phi.hat),
                    shape = 23, fill = "gray", 
               color = "black", size = 2, alpha = 0.5)+
    geom_text(x = 0.5, y = 0.05, 
            label = lm_eqn(df = phidata,
                           x = phidata$phi.hatSM, 
                           y = phidata$phi.hat), 
            parse = TRUE, color = "black")+
      geom_smooth(aes(x=phi.hatSM, y=phi.hat),
              method = "lm", fullrange=TRUE,
              color = "black", se = F, linetype = "dashed", size = 1)+
    scale_y_continuous(limits = c(0,1))+
    scale_x_continuous(limits = c(0,1))+
    labs(x = expression("Standardized Monitored "*italic(varphi)),
         y = expression("eBird "*italic(varphi)),
         tag = expression(bold("(c)")),
         title = "250 weeks or ~5 years")+
  coord_fixed()+
  theme_classic()
```

```
## Warning in geom_errorbarh(aes(x = phi.hatSM, y = phi.hat, xmax = phi.hatSM + :
## Ignoring unknown aesthetics: x
```

```
Fig3cExt
```

```
## `geom_smooth()` using formula = 'y ~ x'
```

```
## Warning: Removed 226 rows containing non-finite outside the scale range
## (`stat_smooth()`).
```

```
## Warning: Removed 226 rows containing missing values or values outside the scale range
## (`geom_pointrange()`).
```

```
## Warning: Removed 232 rows containing missing values or values outside the scale range
## (`geom_errorbarh()`).
```

```
## Warning: Removed 226 rows containing missing values or values outside the scale range
## (`geom_point()`).
```

```
#and combine in the figure
Fig3ab <- grid.arrange(Fig3aExt, Fig3bExt, ncol = 1)
```

```
Fig3 <- grid.arrange(Fig3ab, Fig3cExt, ncol = 2, widths = c(1.5,1))
```

```
## `geom_smooth()` using formula = 'y ~ x'
```

```
## Warning: Removed 226 rows containing non-finite outside the scale range
## (`stat_smooth()`).
```

```
## Warning: Removed 226 rows containing missing values or values outside the scale range
## (`geom_pointrange()`).
```

```
## Warning: Removed 232 rows containing missing values or values outside the scale range
## (`geom_errorbarh()`).
```

```
## Warning: Removed 226 rows containing missing values or values outside the scale range
## (`geom_point()`).
```

```
#It looks not good in the `Rmd`, but it is saved with good proportions
ggsave("results/Figure3_timeseries_PLocalPersi_250weeks.pdf", 
       plot = Fig3, dpi = 300, width = 16, height = 6, units = "in")

ggsave("results/Figure3_timeseries_PLocalPersi250weeks.png", 
       plot = Fig3, dpi = 300, width = 16, height = 6, units = "in")
```

#### 6.5 Results for simulation window of ~10 years (500 weeks) - higher RMSE and \(\rho\)

```
#Extended figures

phi_data_plot <- snailkites.PP |> 
  left_join(phi.eBird |>
              full_join(phi.SM) |> 
              filter(Timeframe == "500 weeks or ~10 years")) |>
  ungroup() |>
    mutate(phi.hat = phi.hat * coeff,
           SD.phi = SD.phi * coeff,
           phi.hatSM = phi.hatSM * coeffSM,
           SD.phiSM = SD.phiSM * coeffSM) |> 
  pivot_longer(cols = !c(cell, 
                         year, 
                         week, 
                         Timeframe, 
                         Time.t, 
                         observation.date, 
                         Model,
                         ln.lambda,
                         sigma.sqr,
                         tau.sqr,
                         x0),
               names_to = "group",
               values_to = "phi_values") |> 
  drop_na(phi_values)
```

```
## Joining with `by = join_by(Time.t, Timeframe, Model, ln.lambda, sigma.sqr,
## tau.sqr, x0)`
## Joining with `by = join_by(Time.t)`
```

```
phi_data_plot
```

```
#Ribbon lo-hi phi (SD) for the two datasets
ribbon_phi_eBird <- phi_data_plot |>
  filter(group %in% c("phi.hat", "SD.phi")) |>
  pivot_wider(names_from = group, values_from = phi_values, values_fn = {mean}) |>
  mutate(phi_values = phi.hat)

ribbon_phi_SM <- phi_data_plot |>
  filter(group %in% c("phi.hatSM", "SD.phiSM")) |>
  pivot_wider(names_from = group, values_from = phi_values, values_fn = {mean})|>
  mutate(phi_values = phi.hatSM)

  #figurea
Fig3aExt <- ggplot(phi_data_plot |> 
                  filter(group %in% c("phi.hat", "Observed.y")), 
                aes(x = observation.date,
                    y = phi_values,
                    fill = group)) +
  geom_vline(xintercept = snailkites.PP$observation.date[92], 
             color = "gray", linetype = "dashed") +
  geom_ribbon(data = ribbon_phi_eBird,
              aes(ymin = phi.hat-SD.phi, ymax = phi.hat+SD.phi),
              fill = "#91bfdb25") +
#  geom_line(aes(color = group, linetype = group)) +
  geom_segment(data = phi_data_plot |> 
                  filter(group %in% c("Observed.y")),
               aes(x = observation.date,
                   xend = observation.date,
                   y = 0,
                   yend = phi_values),
             color = "#4575b445",
             fill = "#4575b445") +
  geom_point(data = phi_data_plot |> 
                  filter(group %in% c("Observed.y")),
             color = "#4575b445", size = 0.5, fill = "#4575b445", shape = 21)+
  geom_point(data = phi_data_plot |> 
                  filter(group %in% c("phi.hat")) |>
               drop_na(Model),
             shape = 21, fill = "#91bfdb95", color = "black") +
  labs(x = "Observation date",
       y = "Observed weekly high counts",
       tag = expression(bold("(a)")),
       fill = "",
       color = "",
       linetype = "",
       shape = "") +
  scale_y_continuous(sec.axis = sec_axis(trans = ~./coeff, 
                                         name = expression("eBird " * italic(varphi)))) +
  coord_cartesian(ylim = c(0, coeff))+
  scale_x_date(limits = c(min(datesPP$observation.date),
                          max(snailkites.PP$observation.date)),
               breaks = seq(min(datesPP$observation.date),
                          max(snailkites.PP$observation.date), 
                            by = "12 months"), date_labels="%b\n%Y")+
  scale_linetype_manual(values = c("solid",
                                   "solid"),
                     labels = c("eBird weekly high count", 
                                expression("P(local persistence) = " * italic(varphi))))+
  scale_color_manual(values = c("#4575b445", 
                                "#91bfdb45"),
                     labels = c("eBird weekly high count", 
                                expression("P(local persistence) = " * italic(varphi)))) +
  geom_hline(yintercept = N.critical, linetype = "dashed", color = "red")+
  theme_classic() +
  theme(legend.title = element_blank(),
        legend.position = c(0.125,0.7),
        legend.background = element_blank(),
        legend.box.background = element_rect(colour = "black"),
        axis.line.y.right = element_line(color = "#91bfdb"),
        axis.ticks.y.right = element_line(color = "#91bfdb"))
```

```
## Warning in geom_segment(data = filter(phi_data_plot, group %in%
## c("Observed.y")), : Ignoring unknown parameters: `fill`
```

```
Fig3aExt
```

```
#SM figure

  #figurea
Fig3bExt <- ggplot(phi_data_plot |> 
                  filter(group %in% c("phi.hatSM", "abundance.monitored")), 
                aes(x = observation.date,
                    y = phi_values,
                    fill = group)) +
  geom_vline(xintercept = snailkites.PP$observation.date[92], 
             color = "gray", linetype = "dashed") +
  geom_ribbon(data = ribbon_phi_SM,
              aes(ymin = phi.hatSM-SD.phiSM, ymax = phi.hatSM+SD.phiSM),
              fill = "#fc8d5925") +
 #  geom_line(aes(color = group, linetype = group)) +
  geom_segment(data = phi_data_plot |> 
                  filter(group %in% c("abundance.monitored")),
               aes(x = observation.date,
                   xend = observation.date,
                   y = 0,
                   yend = phi_values),
             color = "#d7302745", 
             fill = "#d7302745")+
 geom_point(data = phi_data_plot |> 
                  filter(group %in% c("abundance.monitored")),
             color = "#d7302745", size = 0.5, fill = "#d7302745", shape = 21)+
  geom_point(data = phi_data_plot |> 
                  filter(group %in% c("phi.hatSM")) |>
               drop_na(Model),
             shape = 21, fill = "#fc8d5995", color = "black") +
  labs(x = "Observation date",
       y = "Observed weekly high counts",
       tag = expression(bold("(b)")),
       fill = "",
       color = "",
       linetype = "",
       shape = "") +
  scale_y_continuous(sec.axis = sec_axis(trans = ~./coeffSM, 
                                         name = expression("Standardized Monitored " * italic(varphi)))) +
  coord_cartesian(ylim = c(0, coeffSM))+
  scale_x_date(limits = c(min(datesPP$observation.date),
                          max(snailkites.PP$observation.date)),
               breaks = seq(min(datesPP$observation.date),
                          max(snailkites.PP$observation.date), 
                            by = "12 months"), date_labels="%b\n%Y")+
  scale_linetype_manual(values = c("solid",
                                   "solid"),
                     labels = c("Standardized Monitored", 
                                expression("P(local persistence) = " * italic(varphi))))+
  scale_color_manual(values = c("#d7302745", 
                                "#fc8d5945"),
                     labels = c("Standardized Monitored", 
                                expression("P(local persistence) = " * italic(varphi)))) +
  geom_hline(yintercept = N.criticalSM, 
             linetype = "dashed", color = "red")+
  theme_classic() +
  theme(legend.title = element_blank(),
        legend.position = c(0.125,0.7),
        legend.background = element_blank(),
        legend.box.background = element_rect(colour = "black"),
        axis.line.y.right = element_line(color = "#fc8d59"),
        axis.ticks.y.right = element_line(color = "#fc8d59"))
```

```
## Warning in geom_segment(data = filter(phi_data_plot, group %in%
## c("abundance.monitored")), : Ignoring unknown parameters: `fill`
```

```
Fig3bExt
```

```
#Performance comparison
phidata <- phi.eBird |>
  dplyr::select(Time.t, 
                Timeframe,
                phi.hat, SD.phi) |>
              full_join(phi.SM |>
                          dplyr::select(Time.t, 
                                        Timeframe, 
                                        phi.hatSM, SD.phiSM)) |>
  filter(Timeframe == "500 weeks or ~10 years")
```

```
## Joining with `by = join_by(Time.t, Timeframe)`
```

```
#relationship?
lm(phi.hat~phi.hatSM, data = phidata)
```

```
## 
## Call:
## lm(formula = phi.hat ~ phi.hatSM, data = phidata)
## 
## Coefficients:
## (Intercept)    phi.hatSM  
##      0.3292       0.5370
```

```
Fig3cExt <- ggplot(phidata)+
    geom_abline(slope = 1)+
    geom_pointrange(aes(x = phi.hatSM, y = phi.hat, 
                        ymin = phi.hat-SD.phi, 
                        ymax = phi.hat+SD.phi),
                    shape = 23, fill = "gray", color = "gray",
                    alpha = 0.25) +
    geom_errorbarh(aes(x = phi.hatSM, y = phi.hat,
                       xmax = phi.hatSM+SD.phiSM, 
                       xmin = phi.hatSM-SD.phiSM, height = 0),
                   color = "gray",
                   alpha = 0.25)+
    geom_point(aes(x = phi.hatSM, y = phi.hat),
                    shape = 23, fill = "gray", 
               color = "black", size = 2, alpha = 0.5)+
    geom_text(x = 0.5, y = 0.05, 
            label = lm_eqn(df = phidata,
                           x = phidata$phi.hatSM, 
                           y = phidata$phi.hat), 
            parse = TRUE, color = "black")+
      geom_smooth(aes(x=phi.hatSM, y=phi.hat),
              method = "lm", fullrange=TRUE,
              color = "black", se = F, linetype = "dashed", size = 1)+
    scale_y_continuous(limits = c(0,1))+
    scale_x_continuous(limits = c(0,1))+
    labs(x = expression("Standardized Monitored "*italic(varphi)),
         y = expression("eBird "*italic(varphi)),
         tag = expression(bold("(c)")),
         title = "500 weeks or ~10 years")+
  coord_fixed()+
  theme_classic()
```

```
## Warning in geom_errorbarh(aes(x = phi.hatSM, y = phi.hat, xmax = phi.hatSM + :
## Ignoring unknown aesthetics: x
```

```
Fig3cExt
```

```
## `geom_smooth()` using formula = 'y ~ x'
```

```
## Warning: Removed 226 rows containing non-finite outside the scale range
## (`stat_smooth()`).
```

```
## Warning: Removed 226 rows containing missing values or values outside the scale range
## (`geom_pointrange()`).
```

```
## Warning: Removed 229 rows containing missing values or values outside the scale range
## (`geom_errorbarh()`).
```

```
## Warning: Removed 226 rows containing missing values or values outside the scale range
## (`geom_point()`).
```

```
#and combine in the figure
Fig3ab <- grid.arrange(Fig3aExt, Fig3bExt, ncol = 1)
```

```
Fig3 <- grid.arrange(Fig3ab, Fig3cExt, ncol = 2, widths = c(1.5,1))
```

```
## `geom_smooth()` using formula = 'y ~ x'
```

```
## Warning: Removed 226 rows containing non-finite outside the scale range
## (`stat_smooth()`).
```

```
## Warning: Removed 226 rows containing missing values or values outside the scale range
## (`geom_pointrange()`).
```

```
## Warning: Removed 229 rows containing missing values or values outside the scale range
## (`geom_errorbarh()`).
```

```
## Warning: Removed 226 rows containing missing values or values outside the scale range
## (`geom_point()`).
```

```
#It looks not good in the `Rmd`, but it is saved with good proportions
ggsave("results/Figure3_timeseries_PLocalPersi_500weeks.pdf", 
       plot = Fig3, dpi = 300, width = 16, height = 6, units = "in")

ggsave("results/Figure3_timeseries_PLocalPersi500weeks.png", 
       plot = Fig3, dpi = 300, width = 16, height = 6, units = "in")
```

#### 6.6 Results for simulation window of ~25 years (1250 weeks)

```
#Extended figures

phi_data_plot <- snailkites.PP |> 
  left_join(phi.eBird |>
              full_join(phi.SM) |>
              filter(Timeframe == "1250 weeks or ~25 years")) |>
  ungroup() |>
    mutate(phi.hat = phi.hat * coeff,
           SD.phi = SD.phi * coeff,
           phi.hatSM = phi.hatSM * coeffSM,
           SD.phiSM = SD.phiSM * coeffSM) |> 
  pivot_longer(cols = !c(cell, 
                         year, 
                         week, 
                         Timeframe, 
                         Time.t, 
                         observation.date, 
                         Model,
                         ln.lambda,
                         sigma.sqr,
                         tau.sqr,
                         x0),
               names_to = "group",
               values_to = "phi_values") |> 
  drop_na(phi_values)
```

```
## Joining with `by = join_by(Time.t, Timeframe, Model, ln.lambda, sigma.sqr,
## tau.sqr, x0)`
## Joining with `by = join_by(Time.t)`
```

```
phi_data_plot
```

```
#Ribbon lo-hi phi (SD) for the two datasets
ribbon_phi_eBird <- phi_data_plot |>
  filter(group %in% c("phi.hat", "SD.phi")) |>
  pivot_wider(names_from = group, values_from = phi_values, values_fn = {mean}) |>
  mutate(phi_values = phi.hat)

ribbon_phi_SM <- phi_data_plot |>
  filter(group %in% c("phi.hatSM", "SD.phiSM")) |>
  pivot_wider(names_from = group, values_from = phi_values, values_fn = {mean})|>
  mutate(phi_values = phi.hatSM)

  #figurea
Fig3aExt <- ggplot(phi_data_plot |> 
                  filter(group %in% c("phi.hat", "Observed.y")), 
                aes(x = observation.date,
                    y = phi_values,
                    fill = group)) +
  geom_vline(xintercept = snailkites.PP$observation.date[92], 
             color = "gray", linetype = "dotted") +
  geom_ribbon(data = ribbon_phi_eBird,
              aes(ymin = phi.hat-SD.phi, ymax = phi.hat+SD.phi),
              fill = "#91bfdb25") +
#  geom_line(aes(color = group, linetype = group)) +
  geom_segment(data = phi_data_plot |> 
                  filter(group %in% c("Observed.y")),
               aes(x = observation.date,
                   xend = observation.date,
                   y = 0,
                   yend = phi_values),
             color = "#4575b445",
             fill = "#4575b445") +
  geom_point(data = phi_data_plot |> 
                  filter(group %in% c("Observed.y")),
             color = "#4575b445", 
             size = 0.5, 
             fill = "#4575b445", 
             shape = 21)+
  geom_point(data = phi_data_plot |> 
                  filter(group %in% c("phi.hat")) |>
               drop_na(Model),
              color = "black",
             shape = 21, fill = "#91bfdb95") +
  labs(x = "Observation date",
       y = "Observed weekly high count",
       tag = expression(bold("(a)")),
       fill = "",
       color = "",
       linetype = "",
       shape = "") +
  scale_y_continuous(sec.axis = sec_axis(trans = ~./coeff, 
                                         name = expression("eBird " * italic(varphi)))) +
  coord_cartesian(ylim = c(0, coeff))+
  scale_x_date(limits = c(min(datesPP$observation.date),
                          max(snailkites.PP$observation.date)),
               breaks = seq(min(datesPP$observation.date),
                          max(snailkites.PP$observation.date), 
                            by = "12 months"), date_labels="%b\n%Y")+
  scale_linetype_manual(values = c("solid",
                                   "solid"),
                     labels = c("eBird weekly high count", 
                                expression("P(local persistence) = " * italic(varphi))))+
  scale_color_manual(values = c("#4575b475", 
                                "#91bfdb75"),
                     labels = c("eBird weekly high count", 
                                expression("P(local persistence) = " * italic(varphi)))) +
  geom_hline(yintercept = N.critical, 
             linetype = "dashed", 
             color = "red")+
    theme_classic()+
  theme(legend.title = element_blank(),
        legend.position = c(0.125,0.7),
        legend.background = element_blank(),
        legend.box.background = element_rect(colour = "black"),
        axis.line.y.right = element_line(color = "#91bfdb"),
        axis.ticks.y.right = element_line(color = "#91bfdb"))
```

```
## Warning in geom_segment(data = filter(phi_data_plot, group %in%
## c("Observed.y")), : Ignoring unknown parameters: `fill`
```

```
Fig3aExt
```

```
#SM figure

  #figurea
Fig3bExt <- ggplot(phi_data_plot |> 
                  filter(group %in% c("phi.hatSM", "abundance.monitored")), 
                aes(x = observation.date,
                    y = phi_values,
                    fill = group)) +
  geom_vline(xintercept = snailkites.PP$observation.date[92], 
             color = "gray", linetype = "dashed") +
  geom_ribbon(data = ribbon_phi_SM,
              aes(ymin = phi.hatSM-SD.phiSM, 
                  ymax = phi.hatSM+SD.phiSM),
              fill = "#fc8d5925") +
#  geom_line(aes(color = group, linetype = group)) +
  geom_segment(data = phi_data_plot |> 
                  filter(group %in% c("abundance.monitored")),
               aes(x = observation.date,
                   xend = observation.date,
                   y = 0,
                   yend = phi_values),
             color = "#d7302745", 
             fill = "#d7302745")+
  geom_point(data = phi_data_plot |> 
                  filter(group %in% c("abundance.monitored")),
             color = "#d7302745", size = 0.5, fill = "#d7302745", 
             shape = 21)+
  geom_point(data = phi_data_plot |> 
                  filter(group %in% c("phi.hatSM")) |>
               drop_na(Model),
             shape = 21, color = "black",
             fill = "#fc8d5995") +
  labs(x = "Observation date",
       y = "Observed weekly high count",
       tag = expression(bold("(b)")),
       fill = "",
       color = "",
       linetype = "",
       shape = "") +
  scale_y_continuous(sec.axis = sec_axis(trans = ~./coeffSM, 
                                         name = expression("Standardized Monitored " * italic(varphi)))) +
  coord_cartesian(ylim = c(0, coeffSM))+
  scale_x_date(limits = c(min(datesPP$observation.date),
                          max(snailkites.PP$observation.date)),
               breaks = seq(min(datesPP$observation.date),
                          max(snailkites.PP$observation.date), 
                            by = "12 months"), date_labels="%b\n%Y")+
  scale_linetype_manual(values = c("solid",
                                   "solid"),
                     labels = c("Standardized Monitored", 
                                expression("P(local persistence) = " * italic(varphi))))+
  scale_color_manual(values = c("#d7302745", 
                                "#fc8d5945"),
                     labels = c("Standardized Monitored", 
                                expression("P(local persistence) = " * italic(varphi)))) +
  geom_hline(yintercept = N.criticalSM, 
             linetype = "dashed", color = "red")+
  theme_classic() +
  theme(legend.title = element_blank(),
        legend.position = c(0.125,0.7),
        legend.background = element_blank(),
        legend.box.background = element_rect(colour = "black"),
        axis.line.y.right = element_line(color = "#fc8d59"),
        axis.ticks.y.right = element_line(color = "#fc8d59"))
```

```
## Warning in geom_segment(data = filter(phi_data_plot, group %in%
## c("abundance.monitored")), : Ignoring unknown parameters: `fill`
```

```
Fig3bExt
```

```
#Performance comparison
phidata <- phi.eBird |>
  dplyr::select(Time.t, 
                Timeframe,
                phi.hat, SD.phi) |>
              full_join(phi.SM |>
                          dplyr::select(Time.t, 
                                        Timeframe, 
                                        phi.hatSM, SD.phiSM)) |>
  filter(Timeframe == "1250 weeks or ~25 years")
```

```
## Joining with `by = join_by(Time.t, Timeframe)`
```

```
#relationship?
lm(phi.hat~phi.hatSM, data = phidata)
```

```
## 
## Call:
## lm(formula = phi.hat ~ phi.hatSM, data = phidata)
## 
## Coefficients:
## (Intercept)    phi.hatSM  
##      0.4424       0.4577
```

```
Fig3cExt <- ggplot(phidata)+
    geom_abline(slope = 1)+
    geom_pointrange(aes(x = phi.hatSM, y = phi.hat, 
                        ymin = phi.hat-SD.phi, 
                        ymax = phi.hat+SD.phi),
                    shape = 23, fill = "gray", color = "gray",
                    alpha = 0.25) +
    geom_errorbarh(aes(x = phi.hatSM, y = phi.hat,
                       xmax = phi.hatSM+SD.phiSM, 
                       xmin = phi.hatSM-SD.phiSM, height = 0),
                   color = "gray",
                   alpha = 0.25)+
    geom_point(aes(x = phi.hatSM, y = phi.hat),
                    shape = 23, fill = "gray", 
               color = "black", size = 2, alpha = 0.5)+
    geom_text(x = 0.5, y = 0.05, 
            label = lm_eqn(df = phidata,
                           x = phidata$phi.hatSM, 
                           y = phidata$phi.hat), 
            parse = TRUE, color = "black")+
      geom_smooth(aes(x=phi.hatSM, y=phi.hat),
              method = "lm", fullrange=TRUE,
              color = "black", se = F, linetype = "dashed", size = 1)+
    scale_y_continuous(limits = c(0,1))+
    scale_x_continuous(limits = c(0,1))+
    labs(x = expression("Standardized Monitored "*italic(varphi)),
         y = expression("eBird "*italic(varphi)),
         tag = expression(bold("(c)")),
         title = "1250 weeks or ~25 years")+
  coord_fixed()+
  theme_classic()
```

```
## Warning in geom_errorbarh(aes(x = phi.hatSM, y = phi.hat, xmax = phi.hatSM + :
## Ignoring unknown aesthetics: x
```

```
#and combine in the figure
Fig3ab <- grid.arrange(Fig3aExt, Fig3bExt, ncol = 1)
```

```
Fig3 <- grid.arrange(Fig3ab, Fig3cExt, ncol = 2, widths = c(1.5,1))
```

```
## `geom_smooth()` using formula = 'y ~ x'
```

```
## Warning: Removed 226 rows containing non-finite outside the scale range
## (`stat_smooth()`).
```

```
## Warning: Removed 226 rows containing missing values or values outside the scale range
## (`geom_pointrange()`).
```

```
## Warning: Removed 226 rows containing missing values or values outside the scale range
## (`geom_errorbarh()`).
```

```
## Warning: Removed 226 rows containing missing values or values outside the scale range
## (`geom_point()`).
```

```
#It looks not good in the `Rmd`, but it is saved with good proportions
ggsave("results/Figure3_timeseries_PLocalPersi_150weeks.pdf", 
       plot = Fig3, dpi = 300, width = 16, height = 6, units = "in")

ggsave("results/Figure3_timeseries_PLocalPersi150weeks.png", 
       plot = Fig3, dpi = 300, width = 16, height = 6, units = "in")
```

### 7 Sensitivity analysis

#### 7.1 Temporal thinning

As a sensitivity analysis, we randomly removed 5% of the data in the eBird time series for each year, and projected under the simulation window of ~10 years (500 weeks). This process last ~13 hours.

```
#the original data
snailkites.PP <- readRDS("data_tmp/snailkitesPP.rds")

#number of counts per year in eBird (44 to 52)
snailkites.PP |> 
  dplyr::select(observation.date,Observed.y) |> 
  mutate(year = year(observation.date)) |> 
  group_by(year) |> 
  summarise(n_weeks = n())

#number of counts per year in SM (3 to 10)
snailkites.PP |> 
  dplyr::select(observation.date,abundance.monitored) |> 
  drop_na() |>
  mutate(year = year(observation.date)) |> 
  group_by(year) |> 
  summarise(n_weeks = n())

#recall the number of trajectories
ntraj =50000

# Define percentages for sensitivity analysis
percentages <- seq(0.95, 0.05, -0.05)

# Initialize lists to store results of the sensitivity analysis
results <- list()

# Loop over each percentage
for (p in percentages) {

  # Start timing for each percentage
  StartTime <- Sys.time()

  # Sample tt and yt for the current percentage p
  sampled_data <- snailkites.PP |>
    mutate(year = year(observation.date)) |> 
    group_by(year) |>
    drop_na(Observed.y) |>
    sample_frac(p) |>
    arrange(Time.t)

  # Extract tt and yt
  tt_sampled <- sampled_data$Time.t
  yt_sampled <- log(sampled_data$Observed.y)

  # Define N_critical
  N.critical <- round((1/2) * mean(exp(yt_sampled)),0)

  # End positions to modeling
  end_positions <- which(tt_sampled >= 101) #having the initial phi fixed

  #save φ (SD)
  phi_results <- vector("list", length = length(end_positions))

  #save model parameters
  parms <- vector("list", length = length(end_positions))

  for (i in seq_along(end_positions)) {
    last.tt <- end_positions[i]
    m <- tt_sampled[last.tt]

    #only for ~ten years
    #Forcing EGSS
    
    EGSS.partial <- egss_reml(yt = yt_sampled[1:last.tt],
                                 tt = tt_sampled[1:last.tt],
                                 fguess = guess_egss(yt = yt_sampled[1:last.tt],
                                                     tt = tt_sampled[1:last.tt]));
    
    parms[[i]] <- EGSS.partial$remls

      thres.times <- as.numeric(0:(500)) #~10 years
      len <- max(thres.times) + 1

      sim.mat.eBird <- egss_sim(ntraj,
                                tt = thres.times,
                                parms = EGSS.partial$remls)

      phi <- rep(0, ntraj)
      last.points <- rep(0, ntraj)

      for(n in 1:ntraj){
        Pop.sim <- exp(c(yt_sampled[last.tt], sim.mat.eBird[-1,n]));
        last.points[n] <- Pop.sim[len]

        # How many times each trajectory go below the threshold?
        below.threshold <- sum(Pop.sim < N.critical)
        phi[n] <- 1-(below.threshold/len)
      } #n simulated trajectories

      #Expected value of probability of local persistence, and SD
      phi_mean <- mean(phi)
      phi_SD <- sqrt(var(phi))
      phi_1st <- as.numeric(quantile(phi, probs = 0.25))
      phi_3rd <- as.numeric(quantile(phi, probs = 0.75))

    #see advance by printing results
    print(cbind(c(p*100),
                (phi_mean-phi_SD),
                phi_1st,
                phi_mean,
                phi_3rd,
                (phi_mean+phi_SD)))

    # Store results in lists for each percentage
     phi_results[[i]] <- cbind(phi_1st, phi_mean, phi_3rd, phi_SD)
  }# i: each week of estimation

  # End timing for this percentage
  EndTime <- Sys.time()
  timelast <- EndTime - StartTime

  # Store results for this percentage
  results[[paste0("p", round(p*100))]] <- list(
    Time.t = tt_sampled[end_positions],
    phi_sensitivity = phi_results,
    parameters = parms,
    time_taken = timelast
  )
}
```

Recover and save the results as data frame for comparison (with an iterative process for each percentage).

```
# Initialize an empty data frame to store results
phiSD_df_sensitivity <- data.frame(
  Percentage = numeric(),
  Time.t = numeric(),
  phi.1st = numeric(),
  phi.hat = numeric(),
  phi.3rd = numeric(),
  SD.phi = numeric(),
  Time_Taken = character(),
  ln.lambda = numeric(),
  sigma.sqr = numeric(),
  tau.sqr = numeric(),
  x0 = numeric()
)

# Loop through each percentage in the results
for (p in names(results)) {
  for (i in seq_along(results[[p]]$phi_sensitivity)) {
    # Extract data for this specific percentage p and end position = week i
    phi.1st = results[[p]]$phi_sensitivity[[i]][1]
    phi.hat = results[[p]]$phi_sensitivity[[i]][2]
    phi.3rd = results[[p]]$phi_sensitivity[[i]][3]
    SD.phi <- results[[p]]$phi_sensitivity[[i]][4]
    time_t <- results[[p]]$Time.t[i]
    time_taken <- results[[p]]$time_taken
    ln.lambda <- results[[p]]$parameters[[i]][1]
    sigma.sqr <- results[[p]]$parameters[[i]][2]
    tau.sqr <- results[[p]]$parameters[[i]][3]
    x0 <- results[[p]]$parameters[[i]][4]
    
    # Combine the extracted data into a data frame
    temp_df <- data.frame(
      Percentage = p,
      Time.t = time_t,
      phi.1st = phi.1st,
      phi.hat = phi.hat,
      phi.3rd = phi.3rd,
      SD.phi = SD.phi,
      Time_Taken = time_taken,
      Trend = ln.lambda,
      Env.noise = sigma.sqr,
      Obs.noise = tau.sqr,
      N_0 = round(exp(x0))
    )
    
    # Bind the temp_df to the final data frame
    phiSD_df_sensitivity <- rbind(phiSD_df_sensitivity, temp_df)
  }
}

head(phiSD_df_sensitivity)
tail(phiSD_df_sensitivity)

# Save the data frame to a file
saveRDS(phiSD_df_sensitivity, "results/phi_SD_Sensitivity_model_parameters.rds")
```

##### 7.1.1 Correlation of reduced data and standardized monitoring data

With the data saved, we can compare how the reduction of data affect the inference.

```
#Load data saved
  #data organized
  datesPP <- readRDS("data_tmp/datesTimeseriesPaynesPrairie.rds")
  snailkites.PP <- readRDS("data_tmp/snailkitesPP.rds")
  #local persistence from Standardized Monitored
  phi.SM.500 <-readRDS("results/phi_hat_SD_SM_model_parameters.rds") |>
    left_join(datesPP) |>
    filter(Timeframe == "500 weeks or ~10 years")
```

```
## Joining with `by = join_by(Time.t)`
```

```
  #local persistence from eBird
  phi.eBird.500 <- readRDS("results/phi_hat_SD_eBird_model_parameters.rds") |>
    left_join(datesPP) |>
    filter(Timeframe == "500 weeks or ~10 years")
```

```
## Joining with `by = join_by(Time.t)`
```

```
  #data reduced
  phiSD_df_sensitivity <- readRDS("results/phi_SD_Sensitivity_model_parameters.rds") |>
    left_join(datesPP) |>
    full_join(phi.eBird.500 |>
                mutate(Percentage = "All",
                       x0 = round(exp(x0))) |>
                rename(Trend = ln.lambda,
                       Env.noise = sigma.sqr,
                       Obs.noise = tau.sqr,
                       N_0 = x0))
```

```
## Joining with `by = join_by(Time.t)`
## Joining with `by = join_by(Percentage, Time.t, phi.1st, phi.hat, phi.3rd,
## SD.phi, Trend, Env.noise, Obs.noise, N_0, observation.date)`
```

For example, using Pearson correlation coefficient

```
#generate a dataframe to estimate correlation and make a figure
cor_sen <- phiSD_df_sensitivity |>
  left_join(phi.SM.500 |>
              mutate(x0 = round(exp(x0))) |>
              rename(Trend = ln.lambda,
                       Env.noise = sigma.sqr,
                       Obs.noise = tau.sqr,
                       N_0 = x0),
            by = c("Time.t", "observation.date"))

#correlation estimation for each percentage
cor_sen |>
  group_by(Percentage) |>
  summarise(correlation = cor(phi.hatSM, 
                              phi.hat, 
                              use = "pairwise.complete.obs"),
            n_obs = n())
```

```
#overlapping weeks
cor_sen |> 
  filter(Timeframe.y == "500 weeks or ~10 years") |> #this is SM
  group_by(Percentage) |>
  summarise(correlation = cor(phi.hatSM, 
                              phi.hat, 
                              use = "pairwise.complete.obs"),
            n_obs = n())
```

```
#labeller object for the `facet_wrap()`
sensilab <- c(
  'All' = "Complete (n = 258; 32)",
  'p95' = "95% dataset (n = 245; 31)",
  'p90' = "90% dataset (n = 233; 30)",
  'p85' = "85% dataset (n = 218; 29)",
  'p80' = "80% dataset (n = 209; 26)",
  'p75' = "75% dataset (n = 195; 24)",
  'p70' = "70% dataset (n = 181; 24)",
  'p65' = "65% dataset (n = 169; 25)",
  'p60' = "60% dataset (n = 154; 20)",
  'p55' = "55% dataset (n = 144; 20)",
  'p50' = "50% dataset (n = 127; 14)",
  'p45' = "45% dataset (n = 112; 14)",
  'p40' = "40% dataset (n = 104; 12)",
  'p35' = "35% dataset (n = 89; 11)",
  'p30' = "30% dataset (n = 79; 12)",
  'p25' = "25% dataset (n = 66; 7)",
  'p20' = "20% dataset (n = 50; 4)",
  'p15' = "15% dataset (n = 38; 4)",
  'p10' = "10% dataset (n = 26; 2)",
  'p5' = "5% dataset (n = 13; 2)"
)

Fig5Ext_a <- ggplot(cor_sen,
       aes(x = phi.hatSM, y = phi.hat))+
  geom_abline(slope = 1)+
  facet_wrap(~factor(Percentage,
                     levels = c("All","p95","p90","p85","p80",
                                "p75","p70","p65","p60","p55",
                                "p50","p45","p40","p35","p30",
                                "p25","p20","p15","p10","p5")),
             ncol = 5, 
             labeller = as_labeller(sensilab))+
  geom_point()+
  geom_smooth(method = "lm", fullrange = T)+
  stat_regline_equation(label.x = 0.35, label.y = 0.2, size = 2.5) +
  stat_cor(method = "pearson", 
         aes(label = paste("rho == ", ..r.., "*','~~p == ", ..p..)), 
         label.x = 0.25, label.y = 0.1, size = 2.5, parse = TRUE)+
  labs(x = expression(varphi[SM]),
       y = expression(varphi[eBird]),
       title = expression("Persistence probability ("~varphi~") comparison"))+
  scale_x_continuous(limits = c(0,1))+
  scale_y_continuous(limits = c(0,1))+
  coord_equal(ratio = 1)+
  theme_bw() +
  theme(panel.grid.major = element_blank(),
        panel.grid.minor = element_blank(),
        strip.background = element_blank(),
        panel.border = element_rect(colour = "black", fill = NA))

Fig5Ext_a
```

```
## `geom_smooth()` using formula = 'y ~ x'
```

```
#lets saved it with good proportions
ggsave("results/Figure5ext_a_Sensitivity_correlations.pdf", 
       plot = Fig5Ext_a, dpi = 300, width = 9, height = 8, units = "in")
```

```
## `geom_smooth()` using formula = 'y ~ x'
```

```
ggsave("results/Figure5ext_a_Sensitivity_correlations.png", 
       plot = Fig5Ext_a, dpi = 300, width = 9, height = 8, units = "in")
```

```
## `geom_smooth()` using formula = 'y ~ x'
```

##### 7.1.2 Visual inspection of trend after data reduction

And we can see the differences in a figure

```
Fig5Ext_b <- ggplot(data = phiSD_df_sensitivity,
                    aes(x = observation.date))+
  facet_wrap(~factor(Percentage,
                     levels = c("All","p95","p90","p85","p80",
                                "p75","p70","p65","p60","p55",
                                "p50","p45","p40","p35","p30",
                                "p25","p20","p15","p10","p5")),
             ncol = 5,
             labeller = as_labeller(sensilab))+
      geom_vline(xintercept = snailkites.PP$observation.date[92], 
             color = "gray", linetype = "dotted") +
    geom_linerange(aes(ymin = phi.hat-SD.phi,
                       ymax = phi.hat+SD.phi,
                      y = phi.hat),
                   alpha = 0.15, color = "#91bfdb95") +
    geom_linerange(aes(ymin = phi.1st,
                       ymax = phi.3rd,
                      y = phi.hat),
                   alpha = 0.15, size = 1, color = "#91bfdb95") +
    geom_point(aes(y = phi.hat),
               alpha = 0.65, color = "#91bfdb95", shape = 21,
               fill = "#91bfdb95") +
    geom_linerange(data = phi.SM.500,
                   aes(y = phi.hatSM,
                       ymin = phi.hatSM-SD.phiSM,
                       ymax = phi.hatSM+SD.phiSM),
                   alpha = 0.15, color = "#fc8d5995") +
    geom_linerange(data = phi.SM.500,
                   aes(y = phi.hatSM, 
                       ymin = phi.1stSM,
                       ymax = phi.3rdSM),
                   alpha = 0.15, size = 1, color = "#fc8d5995") +
    geom_point(data = phi.SM.500,
               aes(y = phi.hatSM),
                alpha = 0.65, color = "#fc8d5995", shape = 24,
               fill = "#fc8d5995") +
    scale_y_continuous(expand = c(0,0))+
   labs(x = "Observation date",
       y = expression(paste("Persistence (",italic(varphi),")")))  +
    coord_cartesian(ylim = c(0,1))+
  theme_classic() +
  theme(panel.grid.major = element_blank(),
        panel.grid.minor = element_blank(),
        strip.background = element_blank(),
        panel.border = element_rect(colour = "black", fill = NA),
        legend.position = "none")

Fig5Ext_b
```

```
#lets saved it with good proportions
ggsave("results/Figure5ext_b_sensitivity_datareduction.pdf", 
       plot = Fig5Ext_b, dpi = 300, width = 10, height = 6, units = "in")

ggsave("results/Figure5ext_b_sensitivity_datareduction.png", 
       plot = Fig5Ext_b, dpi = 300, width = 10, height = 6, units = "in")
```

##### 7.1.3 Root Mean Square Error for each list in the reduced dataset

The trends are very similar and include high Pearson correlation (\(>0.9\)). We calculated the Root Mean Square Error (RMSE) for each percentage to see, on average, how the overlapping weeks differ from each other (lower values indicate close estimate).

```
phiSD_df_sensitivity |> 
  dplyr::select(Time.t,
         Percentage,
         phi.hat) |> 
  left_join(phi.SM.500  |>
              dplyr::select(Time.t,
                     phi.hatSM)) |>
  group_by(Percentage) |>
  drop_na(phi.hatSM) |>
  summarise(rmse = sqrt(mean(phi.hatSM - phi.hat)^2)) |> #
  ggplot(aes(x = factor(Percentage,
                     levels = c("All","p95","p90","p85","p80",
                                "p75","p70","p65","p60","p55",
                                "p50","p45","p40","p35","p30",
                                "p25","p20","p15","p10","p5")),
             y = rmse))+
    geom_segment(aes(y = 0, yend = rmse), 
                 color = "#909090")+
    geom_point(color = "#838383") +
    geom_hline(yintercept = 0.1, 
               linetype = "dashed")+
    labs(x = "Data reduction",
         y = "Root Mean Square Error")+
    theme_classic()
```

```
## Joining with `by = join_by(Time.t)`
```

Most reduced datasets have low RMSE (<0.1), which indicates that, on average, the close persistence estimates between the Standardized Monitoring and eBird is maintained even with observation reduced data. We can plot the Absolute error difference for overlapping weeks of counts.

```
#labeller object for the `facet_wrap()`
sensilab2 <- c(
  'All' = "All (258; 32)",
  'p95' = "95% (245; 31)",
  'p90' = "90% (233; 30)",
  'p85' = "85% (218; 29)",
  'p80' = "80% (209; 26)",
  'p75' = "75% (195; 24)",
  'p70' = "70% (181; 24)",
  'p65' = "65% (169; 25)",
  'p60' = "60% (154; 20)",
  'p55' = "55% (144; 20)",
  'p50' = "50% (127; 14)",
  'p45' = "45% (112; 14)",
  'p40' = "40% (104; 12)",
  'p35' = "35% (89; 11)",
  'p30' = "30% (79; 12)",
  'p25' = "25% (66; 7)",
  'p20' = "20% (50; 4)",
  'p15' = "15% (38; 4)",
  'p10' = "10% (26; 2)",
  'p5' = "5% (13; 2)"
)

datesPP |>
  full_join(phiSD_df_sensitivity |> 
  dplyr::select(Time.t,
         Percentage,
         phi.hat)) |>
  left_join(phi.SM.500  |>
              dplyr::select(Time.t,
                     phi.hatSM)) |>
  group_by(Percentage) |>
  drop_na() |>
  mutate(Abs.diff = (phi.hatSM - phi.hat)) |>
  ggplot(aes(x = observation.date, y = Abs.diff)) +
    geom_segment(aes(x = observation.date,
                     xend = observation.date, 
                     y = 0, 
                     yend = Abs.diff),
                 color = "#909090")+
    geom_point(color = "#838383")+
    geom_hline(yintercept = 0, color = "#838383",
             linetype = "dotted")+
    scale_x_date(limits = c(snailkites.PP$observation.date[92],
                          max(snailkites.PP$observation.date)),
               breaks = seq(min(datesPP$observation.date),
                          max(snailkites.PP$observation.date), 
                            by = "24 months"), date_labels="%Y")+
    facet_wrap(~factor(Percentage,
                     levels = c("All","p95","p90","p85","p80",
                                "p75","p70","p65","p60","p55",
                                "p50","p45","p40","p35","p30",
                                "p25","p20","p15","p10","p5")), 
             ncol = 5,
             labeller = as_labeller(sensilab2))+
    labs(x = "Observation date",
       y = "Absolute error difference")+
    theme_classic() +
    theme(strip.background = element_blank())
```

```
## Joining with `by = join_by(Time.t)`
## Joining with `by = join_by(Time.t)`
```

Again, more negative values indicate a higher estimate of \(\phi\) for a week \(i\) using eBird data when compared with standardized monitoring data across different observation reduced data sets.

#### 7.2 List-level thinning

Another approach to conduct a sensitivity analysis is thinning at the list level. We randomly removed 5% of the data in eBird for each year, and projected under the simulation window of ~10 years (500 weeks). This process last ~13 hours.

```
snailkites.week <- readRDS("data_tmp/SnailKiteCellsWeek.rds")

#recall the number of trajectories
ntraj =50000

# Define percentages for sensitivity analysis
percentages <- seq(0.95, 0.05, -0.05)

# Initialize lists to store results of the sensitivity analysis
results.S2 <- list()

# Loop over each percentage
for (p in percentages) {

  # Start timing for each percentage
  StartTime <- Sys.time()

  # Sample tt and yt for the current percentage p
  sampled_data <- snailkites.week |>
    sample_frac(p) |>
    group_by(cell, year, month, week) |>
    mutate(max_count = max(observation_count, na.rm = T)) |>
    group_by(cell, Time.t) |>
    summarise(Observed.y = round(max(max_count),0)) |>
    filter(cell == CellTop$cell) |>
    drop_na(Observed.y) |>
    arrange(Time.t)
  head(sampled_data)

  # Extract tt and yt
  tt_sampled <- sampled_data$Time.t
  yt_sampled <- log(sampled_data$Observed.y)

  # Define N_critical
  N.critical <- round((1/2) * mean(exp(yt_sampled)),0)

  # End positions to modeling
  end_positions <- which(tt_sampled >= 101) #having the initial phi fixed

  #save φ (SD) IQR
  phi_results <- vector("list", length = length(end_positions))

  #save model parameters
  parms <- vector("list", length = length(end_positions))

  for (i in seq_along(end_positions)) {
    last.tt <- end_positions[i]
    m <- tt_sampled[last.tt]

    # only for ~ten years window simulation
    # Using EGSS
    
    EGSS.partial <- egss_reml(yt = yt_sampled[1:last.tt],
                                 tt = tt_sampled[1:last.tt],
                                 fguess = guess_egss(yt = yt_sampled[1:last.tt],
                                                     tt = tt_sampled[1:last.tt]));
    
    parms[[i]] <- EGSS.partial$remls

      thres.times <- as.numeric(0:(500)) #~10 years
      len <- max(thres.times) + 1

      sim.mat.eBird <- egss_sim(ntraj,
                                tt = thres.times,
                                parms = EGSS.partial$remls)

      phi <- rep(0, ntraj)
      last.points <- rep(0, ntraj)

      for(n in 1:ntraj){
        Pop.sim <- exp(c(yt_sampled[last.tt], sim.mat.eBird[-1,n]));
        last.points[n] <- Pop.sim[len]

        # How many times each trajectory go below the threshold?
        below.threshold <- sum(Pop.sim < N.critical)
        phi[n] <- 1-(below.threshold/len)
      } #n simulated trajectories

      #Expected value of probability of local persistence, and SD
      phi_mean <- mean(phi)
      phi_SD <- sqrt(var(phi))
      phi_1st <- as.numeric(quantile(phi,probs = 0.25))
      phi_3rd <- as.numeric(quantile(phi,probs = 0.75))

      phi_results[[i]] <- cbind(phi_mean,phi_SD,phi_1st,phi_3rd)

    #see advance by printing results
    print(cbind(c(p*100),
                phi_mean-phi_SD,
                phi_1st,
                phi_mean,
                phi_3rd,
                phi_mean+phi_SD))

    # Store results in lists for each percentage
    phi_results[[i]] <- cbind(phi_mean,phi_SD, phi_1st, phi_3rd)
  }# i: each week of estimation

  # End timing for this percentage
  EndTime <- Sys.time()
  timelast <- EndTime - StartTime

  # Store results for this percentage
  results.S2[[paste0("p", round(p*100))]] <- list(
    Time.t = tt_sampled[end_positions],
    phi_sensitivity = phi_results,
    parameters = parms,
    time_taken = timelast
  )
}
```

Recover and save the results as data frame for comparison (with an iterative process for each percentage).

```
# Initialize an empty data frame to store results
phiSD_df_sensitivity.S2 <- data.frame(
  Percentage = numeric(),
  Time.t = numeric(),
  phi.1st = numeric(),
  phi.hat = numeric(),
  phi.3rd = numeric(),
  SD.phi = numeric(),
  Time_Taken = character(),
  ln.lambda = numeric(),
  sigma.sqr = numeric(),
  tau.sqr = numeric(),
  x0 = numeric()
)

# Loop through each percentage in the results
for (p in names(results.S2)) {
  for (i in seq_along(results.S2[[p]]$phi_sensitivity)) {
    # Extract data for this specific percentage p and end position = week i
    phi.1st = results.S2[[p]]$phi_sensitivity[[i]][1],
    phi.hat = results.S2[[p]]$phi_sensitivity[[i]][2],
    phi.3rd = results.S2[[p]]$phi_sensitivity[[i]][3],
    SD.phi <- results.S2[[p]]$phi_sensitivity[[i]][4]
    time_t <- results.S2[[p]]$Time.t[i]
    time_taken <- results.S2[[p]]$time_taken
    ln.lambda <- results.S2[[p]]$parameters[[i]][1]
    sigma.sqr <- results.S2[[p]]$parameters[[i]][2]
    tau.sqr <- results.S2[[p]]$parameters[[i]][3]
    x0 <- results.S2[[p]]$parameters[[i]][4]
    
    # Combine the extracted data into a data frame
    temp_df <- data.frame(
      Percentage = p,
      Time.t = time_t,
      phi.1st = phi.1st,
      phi.hat = phi.hat,
      phi.3rd = phi.3rd,
      SD.phi = SD.phi,
      Time_Taken = time_taken,
      Trend = ln.lambda,
      Env.noise = sigma.sqr,
      Obs.noise = tau.sqr,
      N_0 = round(exp(x0))
    )
    
    # Bind the temp_df to the final data frame
    phiSD_df_sensitivity.S2 <- rbind(phiSD_df_sensitivity.S2, temp_df)
  }
}

head(phiSD_df_sensitivity.S2)
tail(phiSD_df_sensitivity.S2)

# Save the data frame to a file
saveRDS(phiSD_df_sensitivity.S2, "results/phi_SD_SensitivityS2_model_parameters.rds")
```

With the data saved, we can compare how the reduction of data affect the inference.

```
#Load data saved
  #data organized
  datesPP <- readRDS("data_tmp/datesTimeseriesPaynesPrairie.rds")
  #local persistence from Standardized Monitored
  phi.SM.500 <-readRDS("results/phi_hat_SD_SM_model_parameters.rds") |>
    left_join(datesPP) |>
    filter(Timeframe == "500 weeks or ~10 years")
```

```
## Joining with `by = join_by(Time.t)`
```

```
  #local persistence from eBird
  phi.eBird.500 <- readRDS("results/phi_hat_SD_eBird_model_parameters.rds") |>
    left_join(datesPP) |>
    filter(Timeframe == "500 weeks or ~10 years")
```

```
## Joining with `by = join_by(Time.t)`
```

```
  #data reduced
  phiSD_df_sensitivity.S2 <- readRDS("results/phi_SD_SensitivityS2_model_parameters.rds") |>
    left_join(datesPP) |>
    full_join(phi.eBird.500 |>
                mutate(Percentage = "All",
                       x0 = round(exp(x0))) |>
                rename(Trend = ln.lambda,
                       Env.noise = sigma.sqr,
                       Obs.noise = tau.sqr,
                       N_0 = x0))
```

```
## Joining with `by = join_by(Time.t)`
## Joining with `by = join_by(Percentage, Time.t, phi.1st, phi.hat, phi.3rd,
## SD.phi, Trend, Env.noise, Obs.noise, N_0, observation.date)`
```

For example, using Pearson correlation coefficient

```
#generate a dataframe to estimate correlation and make a figure
cor_sen2 <- phiSD_df_sensitivity.S2 |>
  left_join(phi.SM.500 |>
              mutate(x0 = round(exp(x0))) |>
              rename(Trend = ln.lambda,
                       Env.noise = sigma.sqr,
                       Obs.noise = tau.sqr,
                       N_0 = x0),
            by = c("Time.t", "observation.date"))

#correlation estimation for each percentage
cor_sen2 |>
  group_by(Percentage) |>
  summarise(correlation = cor(phi.hatSM, 
                              phi.hat, 
                              use = "pairwise.complete.obs"),
            n_obs = n())
```

```
#overlapping weeks
cor_sen2 |> 
  filter(Timeframe.y == "500 weeks or ~10 years") |> #this is SM
  group_by(Percentage) |>
  summarise(correlation = cor(phi.hatSM, 
                              phi.hat, 
                              use = "pairwise.complete.obs"),
            n_obs = n())
```

```
#labeller object for the `facet_wrap()`
sensilab2 <- c(
  'All' = "All (258; 32)",
  'p95' = "95% (258; 32)",
  'p90' = "90% (257; 32)",
  'p85' = "85% (258; 32)",
  'p80' = "80% (256; 32)",
  'p75' = "75% (258; 32)",
  'p70' = "70% (255; 32)",
  'p65' = "65% (257; 32)",
  'p60' = "60% (256; 32)",
  'p55' = "55% (256; 32)",
  'p50' = "50% (254; 31)",
  'p45' = "45% (253; 32)",
  'p40' = "40% (252; 32)",
  'p35' = "35% (254; 32)",
  'p30' = "30% (254; 32)",
  'p25' = "25% (242; 32)",
  'p20' = "20% (241; 32)",
  'p15' = "15% (213; 30)",
  'p10' = "10% (192; 29)",
  'p5' = "5% (149; 24)"
  )

Fig5Ext_a2 <- ggplot(cor_sen2,
       aes(x = phi.hatSM, y = phi.hat))+
  geom_abline(slope = 1)+
  facet_wrap(~factor(Percentage,
                     levels = c("All","p95","p90","p85","p80",
                                "p75","p70","p65","p60","p55",
                                "p50","p45","p40","p35","p30",
                                "p25","p20","p15","p10","p5")),
             ncol = 5, 
             labeller = as_labeller(sensilab2))+
  geom_point()+
  geom_smooth(method = "lm", fullrange = T)+
  stat_regline_equation(label.x = 0.35, label.y = 0.2, size = 2.5) +
  stat_cor(method = "pearson", 
         aes(label = paste("rho == ", ..r.., "*','~~p == ", ..p..)), 
         label.x = 0.25, label.y = 0.1, size = 2.5, parse = TRUE)+
  labs(x = expression(varphi[SM]),
       y = expression(varphi[eBird]),
       title = expression("Persistence probability ("~varphi~") 2nd comparison"))+
  scale_x_continuous(limits = c(0,1))+
  scale_y_continuous(limits = c(0,1))+
  coord_equal(ratio = 1)+
  theme_bw() +
  theme(panel.grid.major = element_blank(),
        panel.grid.minor = element_blank(),
        strip.background = element_blank(),
        panel.border = element_rect(colour = "black", fill = NA))

Fig5Ext_a2
```

```
## `geom_smooth()` using formula = 'y ~ x'
```

```
#lets saved it with good proportions
ggsave("results/Figure5ext_a_Sensitivity2_correlations.pdf", 
       plot = Fig5Ext_a2, dpi = 300, width = 9, height = 8, units = "in")
```

```
## `geom_smooth()` using formula = 'y ~ x'
```

```
ggsave("results/Figure5ext_a_Sensitivity2_correlations.png", 
       plot = Fig5Ext_a2, dpi = 300, width = 9, height = 8, units = "in")
```

```
## `geom_smooth()` using formula = 'y ~ x'
```

##### 7.2.1 Visual inspection of trend after data reduction

And we can see the differences in a figure

```
Fig5Ext_b2 <- ggplot(data = phiSD_df_sensitivity.S2,
         aes(x = observation.date))+
  facet_wrap(~factor(Percentage,
                     levels = c("All","p95","p90","p85","p80",
                                "p75","p70","p65","p60","p55",
                                "p50","p45","p40","p35","p30",
                                "p25","p20","p15","p10","p5")),
             ncol = 5,
             labeller = as_labeller(sensilab2))+
      geom_vline(xintercept = snailkites.PP$observation.date[92], 
             color = "gray", linetype = "dotted") +
    geom_linerange(aes(y = phi.hat, 
                       ymin = phi.hat-SD.phi,
                       ymax = phi.hat+SD.phi),
                   alpha = 0.15, color = "#91bfdb95") +
    geom_linerange(aes(y = phi.hat, 
                       ymin = phi.1st,
                       ymax = phi.3rd),
                   alpha = 0.15, size = 1, color = "#91bfdb95") +
    geom_point(aes(y = phi.hat),
               alpha = 0.65, color = "#91bfdb95", fill =  "#91bfdb95",shape = 21) +
    geom_linerange(data = phi.SM.500,
                   aes(ymin = phi.hatSM-SD.phiSM,
                       ymax = phi.hatSM+SD.phiSM),
                   alpha = 0.15, color = "#fc8d5995") +
    geom_linerange(data = phi.SM.500,
                   aes(ymin = phi.1stSM,
                       ymax = phi.3rdSM),
                   alpha = 0.15, size = 1, color = "#fc8d5995") +
    geom_point(data = phi.SM.500,
               aes(y = phi.hatSM),
                alpha = 0.65, color = "#fc8d5995", fill = "#fc8d5995", shape = 24) +
    scale_y_continuous(expand = c(0,0))+
   labs(x = "Observation date",
       y = expression(paste("Persistence (",italic(varphi),")")))  +
    coord_cartesian(ylim = c(0,1))+
  theme_classic() +
  theme(panel.grid.major = element_blank(),
        panel.grid.minor = element_blank(),
        strip.background = element_blank(),
        panel.border = element_rect(colour = "black", fill = NA),
        legend.position = "none")

Fig5Ext_b2
```

```
#lets saved it with good proportions
ggsave("results/Figure5ext_b_sensitivity2_datareduction.pdf", 
       plot = Fig5Ext_b2, dpi = 300, width = 10, height = 7, units = "in")

ggsave("results/Figure5ext_b_sensitivity2_datareduction.png", 
       plot = Fig5Ext_b2, dpi = 300, width = 10, height = 7, units = "in")
```

##### 7.2.2 Root Mean Square Error for each list in the (list) reduced dataset

The trends are very similar and include high Pearson correlation (\(>0.9\)). We calculated the Root Mean Square Error (RMSE) for each percentage to see, on average, how the overlapping weeks differ from each other (lower values indicate close estimate).

```
phiSD_df_sensitivity.S2 |> 
  dplyr::select(Time.t,
         Percentage,
         phi.hat) |> 
  left_join(phi.SM.500  |>
              dplyr::select(Time.t,
                     phi.hatSM)) |>
  group_by(Percentage) |>
  drop_na(phi.hatSM) |>
  summarise(rmse = sqrt(mean(phi.hatSM - phi.hat)^2)) |> #
  ggplot(aes(x = factor(Percentage,
                     levels = c("All","p95","p90","p85","p80",
                                "p75","p70","p65","p60","p55",
                                "p50","p45","p40","p35","p30",
                                "p25","p20","p15","p10","p5")),
             y = rmse))+
    geom_segment(aes(y = 0, yend = rmse), 
                 color = "#909090")+
    geom_point(color = "#838383") +
    geom_hline(yintercept = 0.1, 
               linetype = "dashed")+
    labs(x = "Data reduction - list level",
         y = "Root Mean Square Error")+
    theme_classic()
```

```
## Joining with `by = join_by(Time.t)`
```

All reduced datasets have low RMSE (<0.1), which indicates that, on average, the close persistence estimates between the Standardized Monitoring and eBird is maintained even with observation reduced data. We can plot the Absolute error difference for overlapping weeks of counts.

```
datesPP |>
  full_join(phiSD_df_sensitivity.S2 |> 
  dplyr::select(Time.t,
         Percentage,
         phi.hat)) |>
  left_join(phi.SM.500  |>
              dplyr::select(Time.t,
                     phi.hatSM)) |>
  group_by(Percentage) |>
  drop_na() |>
  mutate(Abs.diff = (phi.hatSM - phi.hat)) |>
  ggplot(aes(x = observation.date, y = Abs.diff)) +
    geom_segment(aes(x = observation.date,
                     xend = observation.date, 
                     y = 0, 
                     yend = Abs.diff),
                 color = "#909090")+
    geom_point(color = "#838383")+
    geom_hline(yintercept = 0, color = "#838383",
             linetype = "dotted")+
    scale_x_date(limits = c(snailkites.PP$observation.date[92],
                          max(snailkites.PP$observation.date)),
               breaks = seq(min(datesPP$observation.date),
                          max(snailkites.PP$observation.date), 
                            by = "24 months"), date_labels="%Y")+
    facet_wrap(~factor(Percentage,
                     levels = c("All","p95","p90","p85","p80",
                                "p75","p70","p65","p60","p55",
                                "p50","p45","p40","p35","p30",
                                "p25","p20","p15","p10","p5")), 
             ncol = 5,
             labeller = as_labeller(sensilab2))+
    labs(x = "Observation date",
       y = "Absolute error difference")+
    theme_classic() +
    theme(strip.background = element_blank())
```

```
## Joining with `by = join_by(Time.t)`
## Joining with `by = join_by(Time.t)`
```

Again, more negative values indicate a higher estimate of \(\phi\) for a week \(i\) using eBird data when compared with standardized monitoring data across different observation reduced data sets.

#### 7.3 Figure 5

Data reduction in time series and correlation for some percentages: 95%, 75%, 50%, 25%, 5%.

```
# Load necessary libraries
library(ggplot2)
library(patchwork)
```

```
## 
## Attaching package: 'patchwork'
```

```
## The following object is masked from 'package:MASS':
## 
##     area
```

```
# Define the sensilab object
sensilab_fig5 <- c(
  'p95' = expression(bold("(a)")~"95% - temporal (n = 245; 31)"),
  'p75' = expression(bold("(b)")~"75% - temporal (n = 195; 24)"),
  'p50' = expression(bold("(c)")~"50% - temporal (n = 129; 12)"),
  'p25' = expression(bold("(d)")~"25% - temporal (n = 66; 7)"),
  'p5' = expression(bold("(e)")~"5% - temporal (n = 13; 2)")
)

# Filter the data for the required percentages
cor_sen_filtered <- cor_sen[cor_sen$Percentage %in% c("p95",
                                                      "p75",
                                                      "p50",
                                                      "p25",
                                                      "p5"), ]

# we can use a function that creates the time series plots
time_series_plot <- function(percentage) {
  ggplot(cor_sen_filtered[cor_sen_filtered$Percentage == percentage, ],
         aes(x = observation.date)) +
    geom_linerange(aes(ymin = phi.hat-SD.phi,
                       ymax = phi.hat+SD.phi, 
             y = phi.hat),
                   alpha = 0.15, color = "#91bfdb95",
             fill = "#91bfdb95", shape = 21) +
    geom_linerange(aes(ymin = phi.1st,
                       ymax = phi.3rd, 
             y = phi.hat),
                   alpha = 0.15, size = 1, color = "#91bfdb95") +
    geom_point(aes(y = phi.hat),
               alpha = 0.65, color = "#91bfdb95") +
    geom_linerange(data = phi.SM.500,
                   aes(y = phi.hatSM,
                       ymin = phi.hatSM-SD.phiSM,
                       ymax = phi.hatSM+SD.phiSM),
                   alpha = 0.15, color = "#fc8d5995") +
    geom_linerange(data = phi.SM.500,
                   aes(y = phi.hatSM, 
                       ymin = phi.1stSM,
                       ymax = phi.3rdSM),
                   alpha = 0.15, size = 1, color = "#fc8d5995") +
    geom_point(data = phi.SM.500,
               aes(y = phi.hatSM),
                alpha = 0.65, color = "#fc8d5995",
               fill = "#fc8d5995", shape = 24) +
    scale_x_date(limits = c(datesPP$observation.date[99],
                            max(datesPP$observation.date)),
                 breaks = seq(datesPP$observation.date[99],
                              max(datesPP$observation.date),
                              by = "24 months"), 
                 date_labels="%b\n%Y")+
    labs(x = "Observation date", 
         y = expression(paste("Persistence (", varphi,")"))) +
    theme_classic() +
    theme(legend.position = "none", 
          strip.background = element_blank()) +
    ggtitle(sensilab_fig5[percentage])
}

# and another funtion that creates correlation plots
correlation_plot <- function(percentage) {
  ggplot(cor_sen_filtered[cor_sen_filtered$Percentage == percentage, ],
         aes(x = phi.hatSM, 
             y = phi.hat)) +
    geom_abline(slope = 1) +
    geom_point() +
    geom_smooth(method = "lm", 
                fullrange = TRUE) +
    stat_regline_equation(label.x = 0.25, 
                          label.y = 0.2, 
                          size = 3) +
    stat_cor(method = "pearson",
             aes(label = paste("rho == ", ..r.., "*','~~p == ", ..p..)),
             label.x = 0.15, 
             label.y = 0.1, 
             size = 3, 
             parse = TRUE) +
    labs(x = expression(paste("Persistence (", varphi[SM],")")), 
         y = expression(paste("Persistence (", varphi[eBird],")"))) +
    scale_x_continuous(limits = c(0, 1)) +
    scale_y_continuous(limits = c(0, 1)) +
    coord_equal(ratio = 1) +
    theme_bw() +
    theme(strip.background = element_blank(),
          plot.margin = unit(c(0.25, 0.25, 0.25, 0.25), "cm"),
          axis.title.x = element_text(margin = margin(t = 10)),
          panel.grid.major = element_blank(),
          panel.grid.minor = element_blank(),
          panel.border = element_rect(colour = "black", fill = NA))
}

# Combine plots using patchwork
Figure5 <- (
  (time_series_plot("p95") | correlation_plot("p95")) /
  (time_series_plot("p75") | correlation_plot("p75")) /
  (time_series_plot("p50") | correlation_plot("p50")) /
  (time_series_plot("p25") | correlation_plot("p25")) /
  (time_series_plot("p5") | correlation_plot("p5"))
)

ggsave("results/Figure5_sensitivity_datareduction.pdf", 
       plot = Figure5, dpi = 300, width = 6.5, height = 12.5, units = "in")
```

```
## `geom_smooth()` using formula = 'y ~ x'
```

```
## `geom_smooth()` using formula = 'y ~ x'
## `geom_smooth()` using formula = 'y ~ x'
## `geom_smooth()` using formula = 'y ~ x'
## `geom_smooth()` using formula = 'y ~ x'
```

```
ggsave("results/Figure5_sensitivity_datareduction.png", 
       plot = Figure5, dpi = 300, width = 6.5, height = 12.5, units = "in")
```

```
## `geom_smooth()` using formula = 'y ~ x'
## `geom_smooth()` using formula = 'y ~ x'
## `geom_smooth()` using formula = 'y ~ x'
## `geom_smooth()` using formula = 'y ~ x'
## `geom_smooth()` using formula = 'y ~ x'
```

```
# Define the second sensilab object
sensilab_fig5b <- c(
  'p95' = expression(bold("(f)")~"95% - sampling (n = 258; 32)"),
  'p75' = expression(bold("(g)")~"75% - sampling (n = 258; 32)"),
  'p50' = expression(bold("(h)")~"50% - sampling (n = 254; 31)"),
  'p25' = expression(bold("(i)")~"25% - sampling (n = 242; 32)"),
  'p5' = expression(bold("(j)")~"5% - sampling (n = 149; 24)")
)

# Filter the data for the required percentages
cor_sen_filtered2 <- cor_sen2[cor_sen2$Percentage %in% c("p95",
                                                      "p75",
                                                      "p50",
                                                      "p25",
                                                      "p5"), ]

# we can use a function that creates the time series plots
time_series_plot2 <- function(percentage) {
  ggplot(cor_sen_filtered2[cor_sen_filtered2$Percentage == percentage, ],
         aes(x = observation.date)) +
    geom_linerange(aes(ymin = phi.hat-SD.phi,
                       ymax = phi.hat+SD.phi),
                   alpha = 0.15, color = "#91bfdb95") +
    geom_linerange(aes(ymin = phi.1st,
                       ymax = phi.3rd),
                   alpha = 0.15, size = 1, color = "#91bfdb95") +
    geom_point(aes(y = phi.hat),
               alpha = 0.65, color = "#91bfdb95",
               fill = "#91bfdb95", shape = 21) +
    geom_linerange(data = phi.SM.500,
                   aes(ymin = phi.hatSM-SD.phiSM,
                       ymax = phi.hatSM+SD.phiSM),
                   alpha = 0.15, color = "#fc8d5995") +
    geom_linerange(data = phi.SM.500,
                   aes(ymin = phi.1stSM,
                       ymax = phi.3rdSM),
                   alpha = 0.15, size = 1, color = "#fc8d5995") +
    geom_point(data = phi.SM.500,
               aes(y = phi.hatSM),
                alpha = 0.65, color = "#fc8d5995", 
               fill = "#fc8d5995", shape = 24) +
    scale_x_date(limits = c(datesPP$observation.date[99],
                            max(datesPP$observation.date)),
                 breaks = seq(datesPP$observation.date[99],
                              max(datesPP$observation.date),
                              by = "24 months"), 
                 date_labels="%b\n%Y")+
    labs(x = "Observation date", 
         y = expression(paste("Persistence (", varphi,")"))) +
    theme_classic() +
    theme(legend.position = "none", 
          strip.background = element_blank()) +
    ggtitle(sensilab_fig5b[percentage])
}

# and another funtion that creates correlation plots
correlation_plot2 <- function(percentage) {
  ggplot(cor_sen_filtered2[cor_sen_filtered2$Percentage == percentage, ],
         aes(x = phi.hatSM, 
             y = phi.hat)) +
    geom_abline(slope = 1) +
    geom_point() +
    geom_smooth(method = "lm", 
                fullrange = TRUE) +
    stat_regline_equation(label.x = 0.25, 
                          label.y = 0.2, 
                          size = 3) +
    stat_cor(method = "pearson",
             aes(label = paste("rho == ", ..r.., "*','~~p == ", ..p..)),
             label.x = 0.15, 
             label.y = 0.1, 
             size = 3, 
             parse = TRUE) +
    labs(x = expression(paste("Persistence (", varphi[SM],")")), 
         y = expression(paste("Persistence (", varphi[eBird],")"))) +
    scale_x_continuous(limits = c(0, 1)) +
    scale_y_continuous(limits = c(0, 1)) +
    coord_equal(ratio = 1) +
    theme_classic() +
    theme(strip.background = element_blank(),
          plot.margin = unit(c(0.25, 0.25, 0.25, 0.25), "cm"),
          axis.title.x = element_text(margin = margin(t = 10)),
          panel.grid.major = element_blank(),
          panel.grid.minor = element_blank(),
          panel.border = element_rect(colour = "black", fill = NA))
}

# Combine plots using patchwork
Figure5b <- (
  (time_series_plot2("p95") | correlation_plot2("p95")) /
  (time_series_plot2("p75") | correlation_plot2("p75")) /
  (time_series_plot2("p50") | correlation_plot2("p50")) /
  (time_series_plot2("p25") | correlation_plot2("p25")) /
  (time_series_plot2("p5") | correlation_plot2("p5"))
)

ggsave("results/Figure5_sensitivity2_datareduction.pdf", 
       plot = Figure5b, dpi = 300, width = 6.5, height = 12.5, units = "in")
```

```
## `geom_smooth()` using formula = 'y ~ x'
## `geom_smooth()` using formula = 'y ~ x'
## `geom_smooth()` using formula = 'y ~ x'
## `geom_smooth()` using formula = 'y ~ x'
## `geom_smooth()` using formula = 'y ~ x'
```

```
ggsave("results/Figure5_sensitivity2_datareduction.png", 
       plot = Figure5b, dpi = 300, width = 6.5, height = 12.5, units = "in")
```

```
## `geom_smooth()` using formula = 'y ~ x'
## `geom_smooth()` using formula = 'y ~ x'
## `geom_smooth()` using formula = 'y ~ x'
## `geom_smooth()` using formula = 'y ~ x'
## `geom_smooth()` using formula = 'y ~ x'
```

Combine the figures of the two sensitivity analysis

```
Fig5All <- (Figure5 | Figure5b)

#lets saved it with good proportions
ggsave("results/Figure5_sensitivity_datareductionx2.pdf", 
       plot = Fig5All, dpi = 300, width = 12, height = 13, units = "in")
```

```
## `geom_smooth()` using formula = 'y ~ x'
## `geom_smooth()` using formula = 'y ~ x'
## `geom_smooth()` using formula = 'y ~ x'
## `geom_smooth()` using formula = 'y ~ x'
## `geom_smooth()` using formula = 'y ~ x'
## `geom_smooth()` using formula = 'y ~ x'
## `geom_smooth()` using formula = 'y ~ x'
## `geom_smooth()` using formula = 'y ~ x'
## `geom_smooth()` using formula = 'y ~ x'
## `geom_smooth()` using formula = 'y ~ x'
```

```
ggsave("results/Figure5_sensitivity_datareductionx2.png", 
       plot = Fig5All, dpi = 300, width = 12, height = 14, units = "in")
```

```
## `geom_smooth()` using formula = 'y ~ x'
## `geom_smooth()` using formula = 'y ~ x'
## `geom_smooth()` using formula = 'y ~ x'
## `geom_smooth()` using formula = 'y ~ x'
## `geom_smooth()` using formula = 'y ~ x'
## `geom_smooth()` using formula = 'y ~ x'
## `geom_smooth()` using formula = 'y ~ x'
## `geom_smooth()` using formula = 'y ~ x'
## `geom_smooth()` using formula = 'y ~ x'
## `geom_smooth()` using formula = 'y ~ x'
```

### 8 Aditional example for other populations of snail kite and monthly \(\hat{\varphi}\)

Let’s explore our approach at a different temporal resolution (month) with four different populations:

- Central Everglades (declining)
- West site of Lake Okeechobee (stable)
- North Lake Tohopekaliga (increasing)
- Western Florida (not monitored)

```
#load saved data
SnailKite <- readRDS("data_tmp/SnailKiteCellsID_filtered.rds")

SnailKite |>
  group_by(cell) |>
  filter(year >= 2018) |>
  summarise(count = n(), 
            mu_lat = mean(latitude), 
            mu_lon = mean(longitude)) |> 
  filter(count >= 40) |>
  arrange(mu_lat) |> 
  as.data.frame()
```

```
#search for the cells ID with high counts and our interest location

  # 2702385: Central Everglades1: 25.75671, -80.76607; 
  # 2702386: Central Everglades2: 25.75906, -80.67722; 
  # 2701656: Central Everglades3: 25.76546, -80.79460; 

  # 2689989: West site of Lake Okeechobee1: 26.98044 -81.09575;
  # 2690719: West site of Lake Okeechobee2: 26.99305 -81.06428;
  # 2690722: East site of Lake Okeechobee: 27.12027 -80.67136;

  # 2678323: Northeast Tohopeliga (Toho1): 28.23773 -81.36858;
  # 2678324: East Lake Tohopeliga (Toho2): 28.26645 -81.27500;
  # 2677594: Northcentral Tohopekaliga (Toho3): 28.28354, -81.39557;

  # 2696545: Western Florida1 (Naples): 26.01485, -81.62422
  # 2690713: Western Florida2 (Harns Marsh): 26.64986, -81.68672
  # 2677588: Western Florida3 (Lakeland): 28.07533, -81.94395

sk.others <- SnailKite |>
  filter(cell %in% c(#Central Everglades
                      "2702385","2702386", "2701656", 
                     # Lake Okeechobee
                      "2689989", "2690719","2690722", 
                     # Lake Tohopeliga
                      "2678323","2678324","2677594",
                     # Western Florida
                      "2696545","2690713","2677588")) |> 
  mutate(population = case_when(cell == "2702385"~"Everglades",
                                cell == "2702386"~"Everglades",
                                cell == "2701656"~"Everglades",
                                cell == "2689989"~"Okeechobee",
                                cell == "2690719"~"Okeechobee",
                                cell == "2690722"~"Okeechobee",
                                cell == "2678323"~"Tohopekaliga",
                                cell == "2678324"~"Tohopekaliga",
                                cell == "2677594"~"Tohopekaliga",
                                cell == "2696545"~"Western Florida",
                                cell == "2690713"~"Western Florida",
                                cell == "2677588"~"Western Florida"),
         seqnum = cell,
         cell = case_when(cell == "2702385"~"Everglades 1",
                                cell == "2702386"~"Everglades 2",
                                cell == "2701656"~"Everglades 3",
                                cell == "2689989"~"Okeechobee W1",
                                cell == "2690719"~"Okeechobee W2",
                                cell == "2690722"~"Okeechobee E",
                                cell == "2678323"~"Toho - NE",
                                cell == "2678324"~"Toho - East Lake Toho",
                                cell == "2677594"~"Toho - NC",
                                cell == "2696545"~"WF - Naples",
                                cell == "2690713"~"WF - Harns Marsh",
                                cell == "2677588"~"WF - Lakeland"))

sk.others.ts <- sk.others |>
  group_by(cell) |>
  filter(year >= 2018) |>
  mutate(Time.t = case_when(year == 2018 ~ month,
                            year > 2018 ~ month+(12*(year-2018)))) |> 
  group_by(population, cell, Time.t) |>
  summarise(Observed.y = round(max(max_count),0),
            observation_date = min(observation_date))
```

```
## `summarise()` has grouped output by 'population', 'cell'. You can override
## using the `.groups` argument.
```

```
ggplot(sk.others.ts, aes(x = observation_date, 
                         y = Observed.y))+
  geom_segment(aes(color = population,
                   y = 0, yend = Observed.y), alpha = 0.5)+
  geom_point(aes(fill = population), shape = 21)+
  labs(x = "Observation date",
       y = "eBird monthly high-counts",
       color = "Population",
       fill = "Population")+
  geom_hline(yintercept = c(2,5), 
             linetype = "dashed",
             color = "red")+
  scale_color_manual(values = c("#FF1493", 
                                "#00bfff", 
                                "#9acd32", 
                                "#800080"))+
  scale_fill_manual(values = c("#FF1493",
                               "#00bfff",
                               "#9acd32",
                               "#800080"))+
  facet_wrap(~cell, ncol = 3, scales = "free_y")+
  theme_classic() +
  theme(legend.position = "none",
        strip.background = element_blank())
```

This figure deployed the monthly high count in eBird for three cells per population, with the \(N\_{c}^{eBird} = 5\) used for Payne’s Prairie (the cell with more data in eBird dataset) and \(N\_{c}^{eBird} = 2\) as one-half the mean observed counts in the nine cells analyzed here.

Where are these localities in Florida?

```
#A global map to make figures ###
world1 <- sf::st_as_sf(maps::map(database = 'world', plot = FALSE, fill = TRUE))
world1
```

```
wrapped_gridSnailKite <- readRDS("data_tmp/wrapped_gridSnailKite.rds") |>
  left_join(sk.others)
```

```
## Joining with `by = join_by(seqnum)`
```

```
ggplot() +
  geom_sf(data = world1)+
  geom_sf(data=wrapped_gridSnailKite, 
          color = "gray") +
  geom_sf(data = wrapped_gridSnailKite |>
                    filter(population %in% c("Everglades",
                                             "Okeechobee",
                                             "Tohopekaliga", 
                                             "Western Florida")),
          aes(color = population, 
              fill = population), alpha = 0.6) +
    coord_sf(xlim = c(-84.5, -79.5), 
           ylim =  c(24.1, 30.9)) +
  scale_color_manual(values = c("#FF1493", 
                                "#00bfff", 
                                "#9acd32", 
                                "#800080"))+
  scale_fill_manual(values = c("#FF1493",
                               "#00bfff",
                               "#9acd32",
                               "#800080"))+
  labs(x = "Longitude",
       y = "Latitude",
       color = "Population",
       fill = "Population")+
  theme_classic() +
  theme(legend.position = "bottom")
```

#### 8.1 Iterate the process for months for each cell - two values of \(N\_{c}\)

And iterate for each cell and each month with data of eBird high count

```
# Initialize lists to store results of the sensitivity analysis
sk.others.results <- list()

ntraj = 50000

cells <- c(unique(sk.others.ts$cell))

# Loop over each percentage
for (k in cells) {
  
  # Sample tt and yt for the current percentage p
  cell_data <- sk.others.ts |>
    ungroup() |>
    drop_na(Observed.y) |>
    filter(cell == k) |>
    arrange(Time.t)
  
  # Extract tt and yt
  tt_sampled <- cell_data$Time.t
  yt_sampled <- log(cell_data$Observed.y)
  
  # Using the same N_critical estimated in Payne's Prairie (location with more data)
  N.critical <- 5 #2 # we can change from the N_critical in PP, 
                      # to the half mean of the populations evaluated = 2
  
  #Initial time vector (for fist model fitting) - first 12 observations
  init.tt <- min(tt_sampled) + 12
  
  # End positions to modeling
  end_positions <- which(tt_sampled >= init.tt)
  
  #to save φ (SD) 
  phi_results <- vector("list", length = length(end_positions))

  #to save model used
  modelSS <- vector("list", length = length(end_positions))
  
  #to save the time last in each week
  timelast <- vector("list", length = length(end_positions))
  
  for (i in seq_along(end_positions)) {
    
    # Start timing for this week i
    StartTime <- Sys.time()
    
    last.tt <- end_positions[i]
    l <- tt_sampled[last.tt]
    
    #only for ~10 years
  
    OUSS.partial <- ouss_reml(yt = yt_sampled[1:last.tt],
                               tt = tt_sampled[1:last.tt],
                               fguess = guess_ouss(yt = yt_sampled[1:last.tt],
                                                   tt = tt_sampled[1:last.tt]))
    
    model <- if(OUSS.partial$remls[2] < 0.025){
      "EGSS"
    }else{
      "OUSS"
    }
    modelSS[[i]] <- model
    
    if(model == "OUSS"){
      
      thres.times <- as.numeric(0:(120)) #~10 years
      len <- max(thres.times) + 1
      
      sim.mat.eBird <- ouss_sim(ntraj, 
                                tt = thres.times, 
                                parms = OUSS.partial$remls)
      
      phi <- rep(0, ntraj)
      last.points <- rep(0, ntraj)
      
      for(n in 1:ntraj){
        Pop.sim <- exp(c(yt_sampled[last.tt], sim.mat.eBird[-1,n]));
        last.points[n] <- Pop.sim[len] 
      
      # How many times each trajectory go below the threshold?
        below.threshold <- sum(Pop.sim < N.critical)
        phi[n] <- 1-(below.threshold/len)
      }

      #Expected value of probability of local persistence, and SD
      phi_mean <- mean(phi)
      phi_SD <- sqrt(var(phi))
      
    }
    else{
      
      EGSS.partial <- egss_reml(yt = yt_sampled[1:last.tt],
                                 tt = tt_sampled[1:last.tt],
                                 fguess = guess_egss(yt = yt_sampled[1:last.tt],
                                                     tt = tt_sampled[1:last.tt]));

      thres.times <- as.numeric(0:(120)) #~10 years
      len <- max(thres.times) + 1
      
      sim.mat.eBird <- egss_sim(ntraj, 
                                tt = thres.times, 
                                parms = EGSS.partial$remls)
      
      phi <- rep(0, ntraj)
      last.points <- rep(0, ntraj)
      
      for(n in 1:ntraj){
        Pop.sim <- exp(c(yt_sampled[last.tt], sim.mat.eBird[-1,n]));
        last.points[n] <- Pop.sim[len] 
      
      # How many times each trajectory go below the threshold?
        below.threshold <- sum(Pop.sim < N.critical)
        phi[n] <- 1-(below.threshold/len)
      }
      
      #Expected value of probability of local persistence, and SD
      phi_mean <- mean(phi)
      phi_SD <- sqrt(var(phi))
  
    }
    
    #see advance by printing results
    print(cbind(k, #cell ID
                l, #month of estimation in the time series
                model, #model selected
                (phi_mean-phi_SD), #lower ribbon
                phi_mean, #Expected value
                (phi_mean+phi_SD))) #higher ribbon
  
      # Store results in lists for each percentage
      phi_results[[i]] <- cbind(phi_mean,phi_SD)
      modelSS[[i]] <- model
      
        # End timing for this week i in the cell k
      EndTime <- Sys.time()
      timelast[[i]] <- EndTime - StartTime
  }

  
  # Store results for this percentage
  sk.others.results[[paste0("cell - ", k)]] <- list(
    Time.t = tt_sampled[end_positions],
    phi_others = phi_results,
    modelSS = modelSS,
    time_taken = timelast
  )
}
```

Recover and save the results as data frame (with an iterative process for each location).

```
# Initialize an empty data frame to store results
phiSD_df_pops <- data.frame(
  Population = numeric(),
  Time.t = numeric(),
  phi.hat = numeric(),
  SD.phi = numeric(),
  Model = character(),
  Time_Taken = character()
)

# Loop through each percentage in the results
for (k in seq_along(cells)) {
  for (i in seq_along(sk.others.results[[k]]$phi_others)) {
    # Extract data for this specific percentage and end position
    phi_hat <- sk.others.results[[k]]$phi_others[[i]][1]
    SD_phi <- sk.others.results[[k]]$phi_others[[i]][2]
    model <- sk.others.results[[k]]$modelSS[[i]]
    time_t <- sk.others.results[[k]]$Time.t[i]
    time_taken <- sk.others.results[[k]]$time_taken[[i]]
    
    # Combine the extracted data into a data frame
    temp_df <- data.frame(
      cell = cells[k],
      Time.t = time_t,
      phi.hat = phi_hat,
      SD.phi = SD_phi,
      Model = model,
      Time_Taken = time_taken
    )
    
    # Bind the temp_df to the final data frame
    phiSD_df_pops <- rbind(phiSD_df_pops, temp_df)
  }
}

head(phiSD_df_pops)
tail(phiSD_df_pops)

# Save the data frame to a file - N_critical 5
saveRDS(phiSD_df_pops, "results/phi_SD_OtherPopulations.rds")
# with N_critical 2
#saveRDS(phiSD_df_pops, "results/phi_SD_OtherPopulations_2.rds")
```

#### 8.2 Results of monthly \(\hat{\phi}\) in three populations of snail kites \(N\_{c} = 5\)

Call the saved results (data frame) to generate the figure of \(\hat{\phi}\) for each sampling unit.

```
phiSD_df_pops <- readRDS("results/phi_SD_OtherPopulations.rds")

phiSD_df_pops |> 
  left_join(sk.others.ts, by = c("cell", 
                                 "Time.t")) |>
  ggplot(aes(x = observation_date, y = phi.hat))+
    geom_line(aes(color = population))+
    geom_ribbon(aes(ymin = phi.hat-SD.phi,
                    ymax = phi.hat+SD.phi,
                    fill = population),
                alpha = 0.25) +
    geom_point(aes(fill = population, shape = Model),
               color = "black")+
  coord_cartesian(ylim = c(0, 1))+
  scale_x_date(breaks = seq(as.Date("2018-01-01"), 
                            as.Date(Sys.Date( )), 
                            by = "24 months"), date_labels="%b-\n%Y")+
  scale_color_manual(values = c("#FF1493", 
                                "#00bfff", 
                                "#9acd32", 
                                "#800080"))+
  scale_fill_manual(values = c("#FF1493",
                               "#00bfff",
                               "#9acd32",
                               "#800080"))+
  scale_shape_manual(values = c(21,24))+
  labs(x = "Observation date",
       y = expression(phi*" - estimated with monthly high-counts in eBird"),
       title = expression("Threshold from Payne's Prairie - "*N[c] ~ "= 5"))+
  facet_wrap(~cell, ncol = 3)+
  theme_classic() +
  theme(legend.position = "bottom",
        strip.background = element_blank()) +
  guides(color = "none",
         fill = "none")
```

This figure confirms the declining trend, with very low persistence for the Everglades subpopulation. The subpopulation in Lake Okeechobee seems more stable in the east (Okeechobee\_E), and there is a potential decline for Okeechobee\_W2. Finally, the recent expanded subpopulation in Lake Tohopekaliga, south of Kissimmee, has lower persistence, but with a potential increase, specially in the NE of the lake. These output can be helpful for researchers and managers to intensify standardized monitoring.

#### 8.3 Results of monthly \(\hat{\phi}\) in three populations of snail kites \(N\_{c} = 2\)

Adjusting the \(N\_{c}^{eBird}=2\), using one-half of the mean of the 9 sampling cells of three subpopulations

```
phiSD_df_pops <- readRDS("results/phi_SD_OtherPopulations_2.rds")

phiSD_df_pops |> 
  left_join(sk.others.ts, by = c("cell", 
                                 "Time.t")) |>
  ggplot(aes(x = observation_date, y = phi.hat))+
    geom_line(aes(color = population))+
    geom_ribbon(aes(ymin = phi.hat-SD.phi,
                    ymax = phi.hat+SD.phi,
                    fill = population),
                alpha = 0.25) +
    geom_point(aes(fill = population, shape = Model),
               color = "black")+
  coord_cartesian(ylim = c(0, 1))+
  scale_x_date(breaks = seq(as.Date("2018-01-01"), 
                            as.Date(Sys.Date( )), 
                            by = "24 months"), date_labels="%b-\n%Y")+
  scale_color_manual(values = c("#FF1493", 
                                "#00bfff", 
                                "#9acd32", 
                                "#800080"))+
  scale_fill_manual(values = c("#FF1493",
                               "#00bfff",
                               "#9acd32",
                               "#800080"))+
  scale_shape_manual(values = c(21,24))+
  labs(x = "Observation date",
       y = expression(phi*" - estimated with monthly high-counts in eBird"),
       title = expression("Threshold from cells evaluated - "*N[c] ~ "= 2"))+
  facet_wrap(~cell, ncol = 3)+
  theme_classic() +
  theme(legend.position = "bottom",
        strip.background = element_blank()) +
  guides(color = "none",
         fill = "none")
```

Although the lower threshold (\(N\_{c}^{eBird}=2\)) provide higher overall values of persistence estimates, the trend for the different supbopulations stands: concerning decline and lower persistence in the Everglades subpopulation; some stable dynamics in the Lake Okeechobee; and, increase in persistence for the subpopulation in Lake Tohopekaliga. Further extension of our approach could help to evaluate other subpopulations (e.g., Cape Coral channels) and include them in robust systematic monitoring.
